# The EGFR-STYK1-FGF1 axis sustains functional drug tolerance to EGFR inhibitors in EGFR-mutant non-small cell lung cancer

**DOI:** 10.1101/2022.04.26.489539

**Authors:** Carolien Eggermont, Philippe Giron, Maxim Noeparast, Hugo Vandenplas, Pedro Aza-Blanc, Gustavo J. Gutierrez, Jacques De Grève

## Abstract

Non-small cell lung cancer (NSCLC) patients harboring activating mutations in epidermal growth factor receptor (EGFR) are sensitive to therapy with EGFR tyrosine kinase inhibitors (TKI). Despite remarkable clinical responses using EGFR TKI, surviving drug tolerant cells serve as a reservoir from which drug resistant tumors may emerge. This study addresses the need for improved efficacy of EGFR TKI by identifying targets involved in functional drug tolerance against them. To this aim, a high-throughput siRNA kinome screen was performed using two EGFR TKI-sensitive EGFR-mutant NSCLC cell lines in the presence/absence of the second-generation EGFR TKI afatinib. From the screen, Serine/Threonine/Tyrosine Kinase 1 (STYK1) was identified as a target that when downregulated potentiates the effects of EGFR inhibition *in vitro*. We found that chemical inhibition of EGFR combined with the siRNA-mediated knockdown of STYK1 led to a significant decrease in cancer cell viability and anchorage-independent cell growth. Further, we show that STYK1 selectively interacts with mutant EGFR and that the interaction is disrupted upon EGFR inhibition. Finally, we identified fibroblast growth factor 1 (FGF1) as a downstream effector of STYK1 in NSCLC cells. Accordingly, downregulation of STYK1 counteracted the afatinib-induced upregulation of FGF1. Altogether, we unveil STYK1 as a valuable target to repress the pool of surviving drug tolerant cells arising upon EGFR inhibition. Co-targeting of EGFR and STYK1 could lead to a better overall outcome for NSCLC patients.

## Introduction

A significant proportion (14-40%) of non-small cell lung cancers (NSCLC) displays activating mutations in the tyrosine kinase domain of epidermal growth factor receptor (EGFR) (1). Lung cancer patients harboring hyperactivating EGFR mutations are eligible for treatment with EGFR tyrosine kinase inhibitors (TKI). Three generations of EGFR TKI are currently approved as first-line treatment for advanced EGFR-mutant NSCLC patients (2–5) and have increased patients’ survival compared to regular chemotherapy. However, cancer recurrence inexorably occurs in these patients due to baseline or acquired drug resistance. Recent studies indicate that a population of functionally drug tolerant ‘persister’ cells survive the drug treatment and eventually acquires mutations resulting in constitutive resistance and therapy failure (6). In contrast to acquired drug resistance, drug tolerance is a reversible state mediated by non-genetic changes sustaining cell survival (7). Elucidating and targeting drug tolerance mechanisms could improve the efficacy of initial EGFR-related treatments by diminishing or eradicating the pool of drug tolerant cells in which constitutive resistance can arise.

With about 500 members, human kinases form one of the most prominent protein families in human cells. Overexpressed and mutated kinases exert crucial roles in cancer-promoting signaling pathways and are intensively investigated as potential drug targets (8). Here, we performed a kinome-wide RNA interference (RNAi) screen to identify kinases responsible for functional tolerance to EGFR TKI in NSCLC cells. Our rationale was that co-targeting of an additional kinase could enhance the sensitivity of NSCLC cells to EGFR TKI.

Serine/Threonine/Tyrosine Kinase 1 (STYK1) has been described as a tyrosine kinase possessing a transmembrane domain, an intracellular kinase domain and a truncated extracellular domain that lacks ligand binding capacity. STYK1 shares 20-30% homology with fibroblast and platelet-derived growth factor receptors and has been reported to display cellular-transforming capabilities and enhance tumor progression by promoting invasion and metastasis (9). However, STYK1 remains a protein with limited functional characterization that may signal through the PI3K/Akt, MAPK, and JAK/STAT pathways (10), but no direct substrates for STYK1 have been described. STYK1 is overexpressed in various cancer types including NSCLC (11–17), and immunohistochemical analyses in tumors have shown that STYK1 is more expressed in NSCLC samples compared to adjacent non-cancerous tissues. Further, NSCLC patients with STYK1-positive cancerous lesions have reduced survival rates (14,18).

We found STYK1 as a potential target for a combination therapy with EGFR inhibition in NSCLC and demonstrate that STYK1 downregulation improves the anti-cancer effects of EGFR TKI *in vitro*. Mechanistically, STYK1 downregulation counteracts the upregulation of FGF1 induced by EGFR TKI in NSCLC cells. Overall, depletion of STYK1 from cancer cells may diminish persistence mechanisms elicited by EGFR inhibition, such as FGF1 upregulation, and in that way reduce/eliminate the pool of drug tolerant cells in which constitutive, genomic resistance may arise.

## Results

### Kinome-wide RNAi screen defines a role for STYK1 in the intrinsic tolerance of EGFR-mutant NSCLC cells to afatinib

To unveil mechanisms of functional tolerance to EGFR TKI, we performed an unbiased RNAi screen targeting 659 known and putative human kinases. To avoid cell context-specific hits, two afatinib-sensitive NSCLC cell lines were included in the screen, PC9 and HCC827, both harboring an oncogenic deletion in exon 19 of EGFR. This ΔE746-A750 mutation represents 45% of all activating EGFR mutations in NSCLC and makes the cells sensitive to EGFR TKI (19,20). In the screen, we used a sub-lethal dose of afatinib adapted to each cell line (as previously determined performing dose-response experiments (21)) (Figure 1A). This methodology allowed us to identify kinases that when downregulated potentiated the effects of afatinib in decreasing the viability of the two NSCLC cell lines (**Error! Reference source not found.**A and Suppl. Table 1).

**Figure 1:**
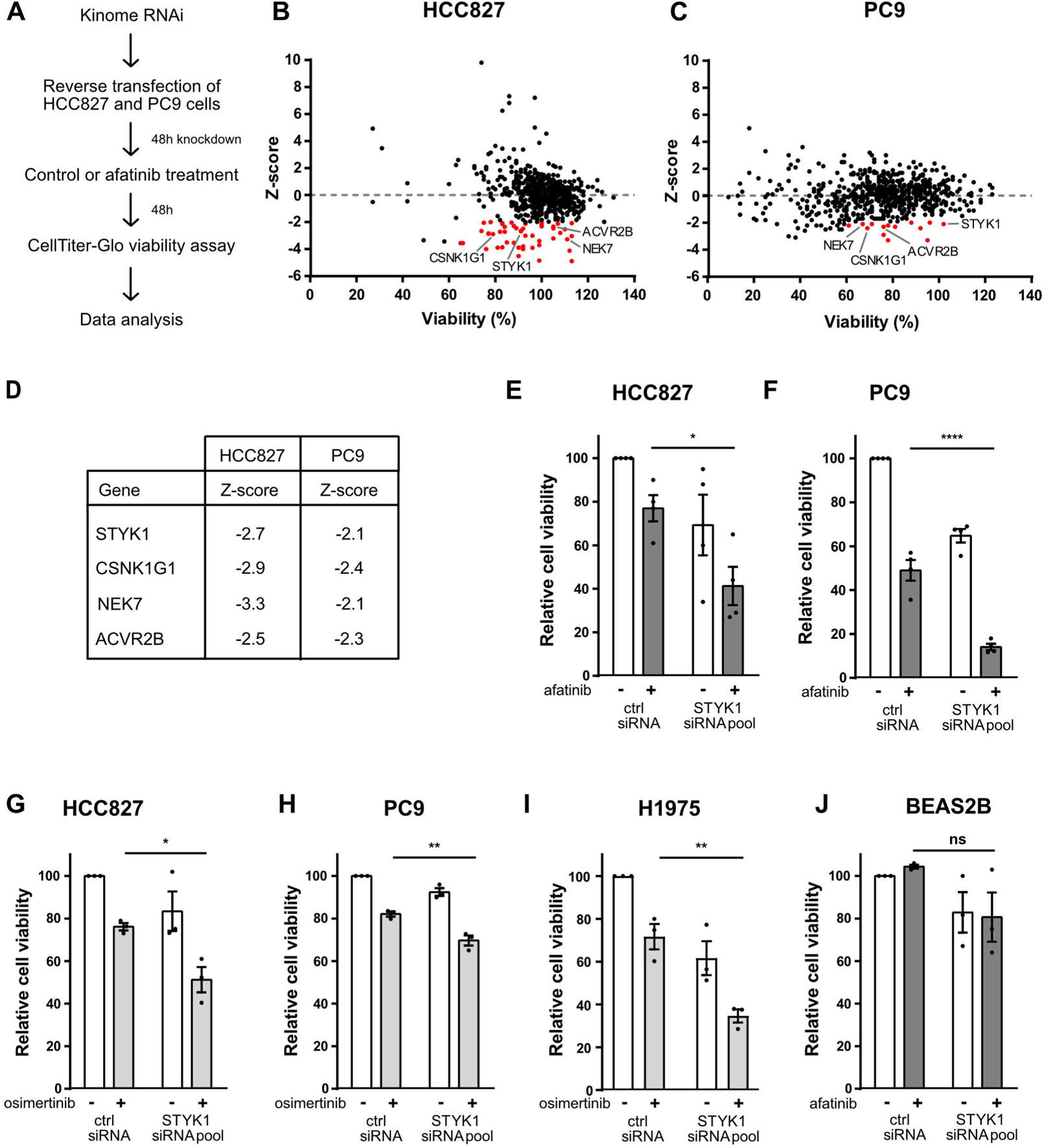
STYK1 supports a prime mechanism of innate tolerance to afatinib in EGFR mutant NSCLC cells. (A) Schematic representation of the kinome-wide RNAi screen. HCC827 and PC9 were reverse transfected with a custom-made siRNA kinome library and 48 hours post-transfection cells were treated with control (DMSO) or afatinib (1nM or 5nM for HCC827 or PC9, respectively). Cell viability was measured after an additional 48 hours of incubation. (B & C) RNAi screen results in HCC827 (B) and PC9 (C) cells. siRNA pools with Z-scores <-2.00 and effects on viability >60% are indicated as red dots. (D) Common hit targets between HCC827 and PC9 with their respective Z-scores. (E-J) Cell viability analysis after reverse transfection of non-targeting control or STYK1 (pool) siRNAs treated for 48 hours with control, afatinib or osimertinib. (E) HCC827 and (F) PC9 cells were treated with 1nM and 5nM afatinib, respectively. (G) HCC827 and (H) PC9 were treated with osimertinib (5nM). (I) H1975 cells were treated with 10nM osimertinib. (J) BEAS-2B cells were treated with 35nM afatinib. Bars represent mean viability ± SEM. At least three independent experiments were performed.

Stringent parameters were used to select candidate hits that were considered for further evaluation: (i) knockdown of target should substantially enhance the sensitivity to afatinib with Z-scores ≤ −2 (see Methods); and (ii) knockdown of target alone should only modestly affect cell viability (less than 40% reduction). Based on these criteria, we identified 50 and 14 siRNA pools (each comprised of four different siRNAs) that sensitized HCC827 and PC9 cells to afatinib, respectively (**Error! Reference source not found.**B, C). Four candidate targets were shared by both cell lines: ACVR2B, NEK7, STYK1 and CSNK1G1 (**Error! Reference source not found.**D). Interestingly, STYK1 (with Z-scores of −2.1 and −2.67 in PC9 and HCC827, respectively) has been reported to be overexpressed in NSCLC where it contributes to tumor growth and metastasis (9,11–17,22). This led us to further explore the exact involvement of STYK1 in NSCLC and its newly discovered potential interplay with mutated EGFR.

We first validated the results from the siRNA screen for STYK1 in dedicated low-throughput experiments and found that combined targeting of EGFR (by afatinib) and STYK1 (by an siRNA pool) caused an extra 35% reduction in cell viability in both HCC827 and PC9 cells, compared to the single afatinib treatment (**Error! Reference source not found.**E, F). Given the current clinical relevance of the third-generation EGFR TKI osimertinib (5), we also investigated the potential of targeting STYK1 combined with osimertinib. We found that the STYK1 siRNA pool readily enhanced osimertinib efficacy in HCC827 and PC9 cells (Figure 1G, H). In addition, in H1975 cells harboring the EGFR L858R/T790M double mutant the STYK1 siRNA pool also significantly enhanced the reduction in cell viability in the presence of osimertinib (Figure 1I). We confirmed these results using single siRNAs from the siRNA pool and found that siRNAs #2 and #3 were the most efficient at knocking down STYK1 mRNA (Suppl. Figure 1A) and protein levels (Suppl. Figure 1B), and accordingly enhanced the effects of afatinib and osimertinib in HCC827 cells (Suppl. Figures 1C, D). Conversely, we found that STYK1 knockdown in BEAS-2B, a normal lung epithelial cell line with wild-type (WT) EGFR, resulted in minimal changes in cell viability, and combination with afatinib did not further affect their viability (**Error! Reference source not found.**J). This highlights the potential selectivity of the combined targeting of STYK1 and EGFR for EGFR-mutant lung cancer cells.

We hypothesized that downregulation of STYK1 would enhance the capacity of afatinib to induce apoptosis in NSCLC cells. We utilized flow cytometry to quantify the apoptotic fraction obtained in cells with STYK1 downregulation in both the presence and absence of afatinib. We observed a trend of enhanced apoptosis when STYK1 siRNAs #2 and #3 were used in afatinib-treated HCC827 and PC9 cells, although those differences were not statistically significant (Suppl. Figures 1E, F).

In agreement with a role for STYK1 in the emergence of drug tolerance mechanisms, we found that STYK1 has been reported to be overexpressed in gefitinib- and WZ4002-tolerant PC9 cells (GSE75602 dataset) (Suppl. Figure 1G) (23). Overall, our data support a role for STYK1 in drug tolerance to EGFR TKI, making it a potential candidate target for increasing the sensitivity of EGFR mutant NSCLC cells to EGFR inhibition and preventing the emergence of drug tolerant cells.

### STYK1 knockdown reduces anchorage-independent growth of EGFR mutant lung cancer cells upon EGFR TKI treatment

Anchorage-independent growth is the gold standard 3D cell culture *in vitro* model for oncogenic transformation and tumorigenesis. To further assess the combinatorial effects of STYK1 knockdown and EGFR inhibition, we employed a classical soft agar colony formation assay as a surrogate for anchorage-independent growth. We generated PC9 cells stably expressing STYK1 short-hairpin RNAs (shRNAs) (Figures 2A, B) and determined the colony formation capacity of STYK1-downregulated cells upon afatinib or osimertinib treatments during a three-week period. As predicted, we found that afatinib and osimertinib hampered the anchorage-independent cell growth of PC9 cells, and this effect was strengthened when combined with STYK1 knockdown (Figures 2C, D). Interestingly, the downregulation of STYK1 alone also compromised colony growth similarly to the EGFR TKI (Figures 2C, D), indicating that STYK1 acts as a tumor-promoting factor in EGFR-mutant cells.

**Figure 2:**
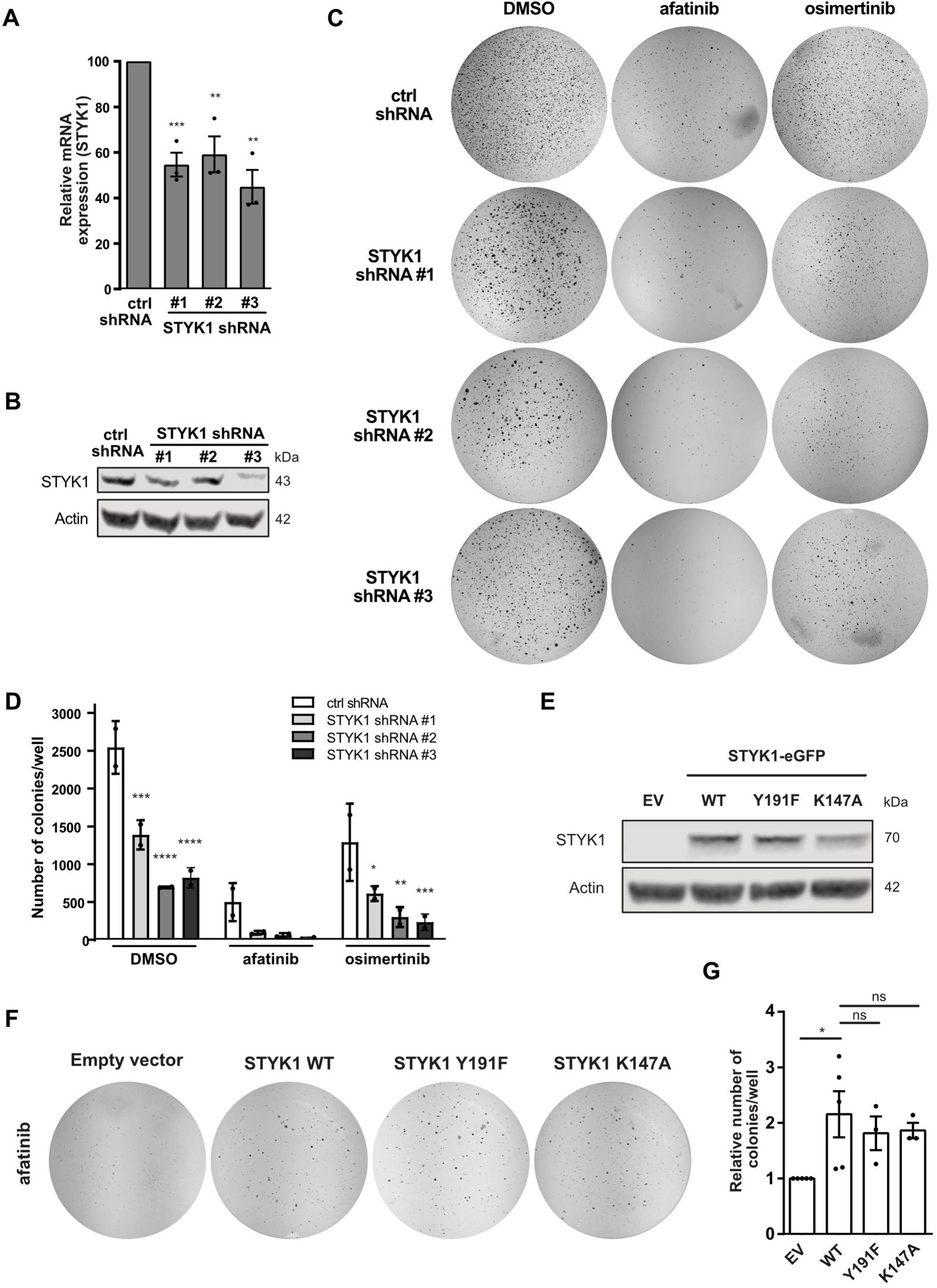
STYK1 alters the anchorage-independent growth of EGFR TKI-treated NSCLC cells. (A-D) PC9 cells were stably transduced with control or STYK1 shRNA plasmids. (A-B) STYK1 knockdown efficiency in three independent experiments as determined by qRT-PCR (mean ± SEM) (A) and Western blotting (representative images shown) (B). β-actin was used as a loading control. (C) The cells were seeded in soft agar medium containing afatinib (5nM) or osimertinib (10nM). Representative images of individual wells stained with nitro blue tetrazolium chloride after 21 days in culture are shown. (D) The average number of colonies (mean ± SEM) of 2 independent experiments is shown as calculated by the OpenCFU software. Statistics are shown for STYK1 shRNA compared to ctrl shRNA for the respective treatments. (E-G) PC9 cells with stable overexpression of empty vector (EV), STYK1-eGFP wild-type (WT), or STYK1-eGFP catalytically inactive mutants Y191F and K147A as confirmed by Western blotting (E). α-tubulin was used as the loading control. (F) The STYK1 overexpressing cells were grown in soft agar in the presence of afatinib (5nM). Representative whole-well images of nitro blue tetrazolium chloride-stained colonies are shown. (G) Quantified results of 3 (or 5 in case of STYK1-WT) independent experiments (mean ± SEM).

We next examined whether enhanced levels of STYK1 could reduce the sensitivity of NSCLC cells to afatinib by performing soft agar assays using PC9 cells constitutively overexpressing STYK1 (Figure 2E). We found a diminished effect of afatinib on the anchorage-independent growth of cells overexpressing STYK1 compared to empty vector controls (Figures 2F, G). In addition, stable overexpression of a kinase-dead version of STYK1 (K147A mutant that does not bind to ATP) (13,24) or a dimerization-impaired mutant of STYK1 (Y191F) also associated with reduced kinase activity (24) increased the formation of colonies in the presence of afatinib treatment in a similar fashion as the WT STYK1 (Figures 2E-G) (24). These observations suggest that the putative kinase activity of STYK1 and its dimerization are not required to enhance the anchorage-independent growth in EGFR-mutant NSCLC cells treated with EGFR TKI. Taken together, STYK1 has an important impact on the tumorigenic potential of NSCLC cells upon EGFR TKI treatment, likely acting in a kinase-independent manner.

### STYK1 colocalizes and interacts with mutant EGFR

According to a previous report, WT EGFR and STYK1 colocalize in endosomes, and STYK1 participates in EGFR trafficking to late endosomes (25). Since mutant EGFR variants have been reported to have a different interactome compared to WT EGFR (21,26), we investigated whether mutant EGFR and STYK1 also colocalize in cells. An immunofluorescence assay was conducted in HCC827 cells, stably producing a STYK1-eGFP fusion protein. We detected colocalization of mutant EGFR with STYK1 at the plasma membrane and in perinuclear vesicles in these cells. Remarkably, all EGFR-containing vesicles appeared positive for STYK1 (Figures 3A, B).

**Figure 3:**
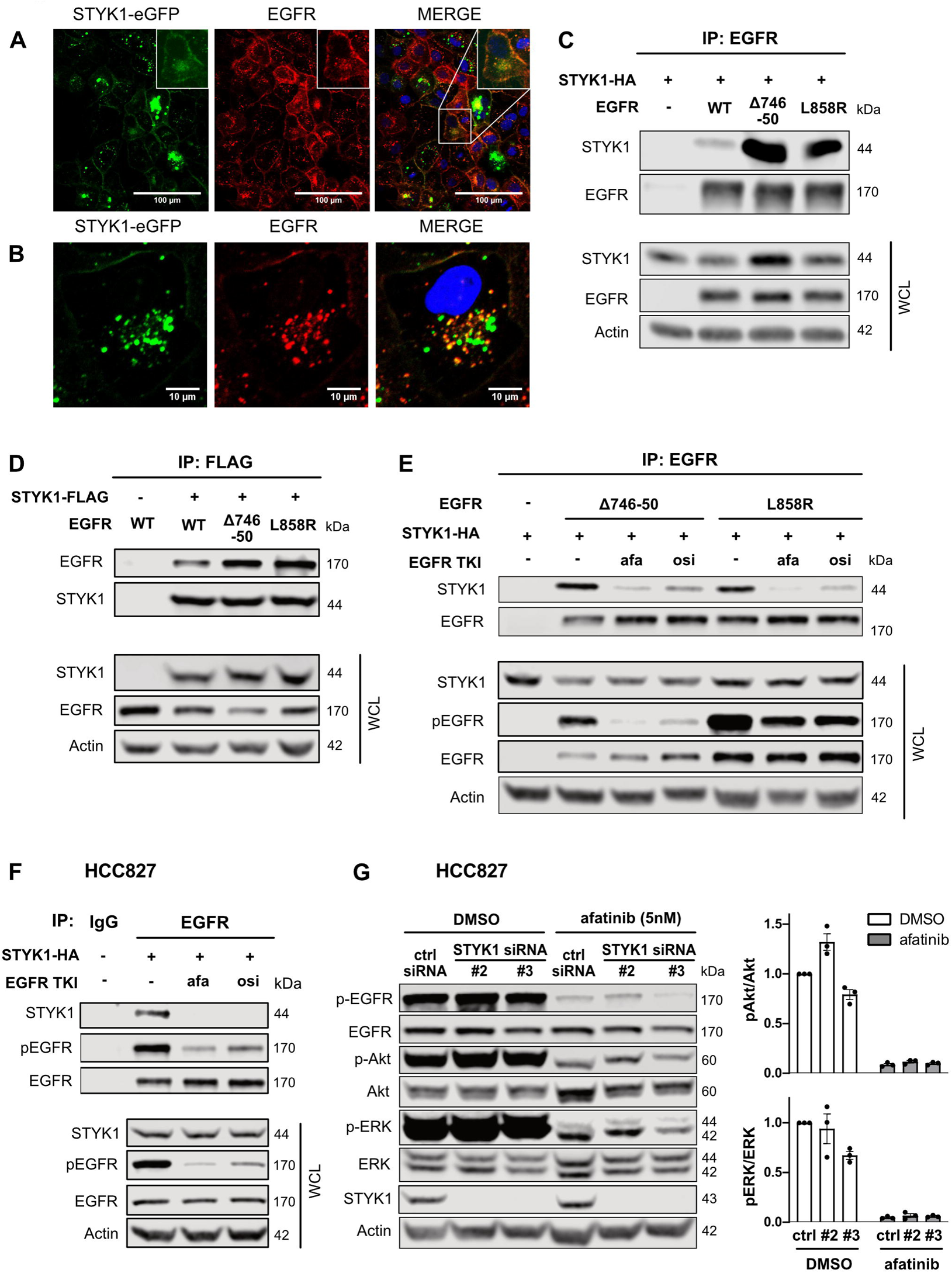
STYK1 colocalizes and preferentially interacts with mutant EGFR in a manner dependent on EGFR activation. (A & B) HCC827 cells were stably transduced with STYK1-eGFP. After fixation, the cells were stained using an anti-EGFR antibody. STYK1-eGFP, EGFR and nuclei (Hoechst) were visualized by Confocal microscopy. Insets show magnifications. (C & D) HEK293T cells were transfected with STYK1-HA or STYK1-FLAG and EGFR (wild-type (WT), Δ746-750, or L858R) and incubated for 24 hours. Cell lysates were subjected to immunoprecipitation with EGFR- (C) and FLAG- (D) antibodies. Immunoprecipitates and whole-cell lysates (WCL) were subjected to Western blotting using the indicated antibodies. (E) HEK293T were transfected with STYK1-HA and EGFR (wild-type (WT), Δ746-50, or L858R) and treated overnight with afatinib (10nM) or osimertinib (10nM). Twenty-four hours post-transfection, cell lysates were subjected to immunoprecipitation using anti-EGFR antibodies. Immunoprecipitates and WCL were subjected to Western blotting using the indicated antibodies. (F) STYK1-HA was transfected in HCC827 cells. After an overnight treatment with afatinib (10nM) or osimertinib (10nM) and a total incubation of 24 hours post-transfection, lysates were prepared and subjected to immunoprecipitation using anti-EGFR or IgG control antibodies. Precipitates and WCL were immunoblotted with the indicated antibodies. (G) HCC827 cells were reverse transfected with control or STYK1 siRNAs, and 48 hours post-transfection 5nM afatinib was added for an additional 24 hours. Total lysates were immunoblotted with the indicated antibodies. pAkt/Akt and pERK/ERK levels were quantified with the help of the LI-COR Odyssey software. Normalized values of three independent experiments are shown as mean ± SEM.

Since mutant EGFR and STYK1 colocalized in the same subcellular compartments, we evaluated their possible interaction by performing co-immunoprecipitation experiments. HEK293T cells were transfected with STYK1 and WT EGFR, EGFR Δ746-750 or L858R activated mutants. We detected a strong interaction between STYK1 and the mutant variants of EGFR by immunoprecipitating either EGFR or STYK1, and while STYK1 also interacted with WT EGFR it was with reduced affinity compared to mutant EGFR (Figures 3C, D). Interestingly, EGFR inhibition by afatinib or osimertinib disrupted the interaction between mutant EGFR (ΔE746-A750 and L858R) and STYK1 (Figures 3E-F). The interaction between STYK1 and endogenous mutant EGFR was confirmed by immunoprecipitating mutant EGFR from NSCLC cells overexpressing STYK1 (Figure 3F & Suppl. Figure 2A). Altogether, our findings indicate that STYK1 and (mutant) EGFR colocalize and interact in a manner dependent on the activation of (mutant) EGFR since the interaction is disrupted in the presence of EGFR TKI.

Since STYK1 and (mutant) EGFR interact in cells, we investigated whether their interplay affected MAPK and PI3K/Akt, the most common EGFR downstream signaling pathways. Previous reports have also shown links between STYK1 and these pathways (9,16,22). We assessed whether MAPK and/or PI3K/Akt pathways were modulated by co-targeting of STYK1 and EGFR in NSCLC cells and found that while afatinib inhibited the activation of ERK and Akt, phosphorylated (active) levels of these kinases were not affected by STYK1 knockdown alone, or further reduced by the combination of STYK1 siRNA and afatinib, compared to afatinib alone (Figure 3G & Suppl. Figure 2B). Overall, our data indicate that the interaction of EGFR and STYK1 does not alter the activation of EGFR downstream signaling and that the interplay between STYK1 downregulation and EGFR inhibition in NSCLC cells is independent of the MAPK and PI3K/Akt pathways.

### Unraveling the effects of STYK1 knockdown on the transcriptome of EGFR mutant NSCLC cells treated with afatinib

To dissect the exact molecular mechanisms involved in the combined effect of STYK1 knockdown and EGFR TKI treatment in NSCLC cells, we performed RNA sequencing (RNAseq) analyses. We assessed the changes in the transcriptome of PC9 cells upon STYK1 siRNA-mediated knockdown in the presence or absence of afatinib (Figure 4A). We identified 87 differentially expressed genes (DEG) which displayed at least a two-fold up- or down-regulation in PC9 cells exposed to afatinib plus STYK1 siRNA *versus* afatinib alone (67 upregulated and 20 downregulated) (Figure 4B & Suppl. Table 2). Importantly, these DEG were shared by the two siRNAs used in the experiment. A KEGG analysis was performed on the DEG revealing two statistically significant enriched pathways: *cAMP signaling* and *Regulation of actin cytoskeleton* (Suppl. Figure 3A). From the top down- and up-regulated DEG (Figure 4C), four genes (ADAMTS1, FGF1, G0S2 and TP63) were also closely associated with cancer pathways according to the literature. We independently evaluated changes in expression levels of these four genes by qRT-PCR using the varying conditions of the RNAseq experiment, and mostly confirmed their differential regulation in PC9 and HCC827 cells (Figures 4D, E & Suppl. Figures 3B-E). The expression levels of G0S2 and TP63 (TAp63 and ΔNp63 isoforms) did not show the same trend in HCC827 (Suppl. Figures 3C-E), while ADAMTS1 and FGF1 clearly appeared to be genes which afatinib-mediated upregulation was blocked by STYK1 siRNAs (Figures 4D, E & Suppl. Figure 4B). Since STYK1 shares approximately 30% homology with members of the fibroblast growth factor receptor (FGFR) family, we focused on Fibroblast Growth Factor 1 (FGF1) (9) and confirmed that STYK1 knockdown alone was able to reduce the baseline FGF1 mRNA levels (Figures 4D-E). More importantly, we also found that STYK1 depletion reduced FGF1 protein levels and counteracted the upregulation of FGF1 induced by afatinib in PC9 cells (Figure 4F). To confirm the regulation of FGF1 by STYK1, we overexpressed STYK1-WT in PC9 cells and evidenced increased FGF1 mRNA levels (Figure 4G). Finally, to assess the role of FGF1 in drug tolerance to EGFR inhibition we determined the viability of PC9 cells treated with FGF1 siRNAs in the presence or absence of EGFR TKI and found that the dual targeting of FGF1 and EGFR enhanced the reduction in cell viability compared to the single treatments (Figure 4H & Suppl. Figure 3F). In addition, in the published GSE75602 dataset (23), we also noticed that FGF1 expression levels were increased in gefitinib- and WZ4002-tolerant PC9 cells (Suppl. Figure 3G), strongly suggesting a role for FGF1 in survival of drug tolerant cells upon EGFR inhibition. Overall, our data demonstrate that FGF1 mRNA and protein expression levels are controlled by STYK1 in untreated NSCLC cells and especially, upon EGFR inhibition. In addition, we establish a role for FGF1 in the drug tolerance mechanisms linked to the use of EGFR TKI.

**Figure 4:**
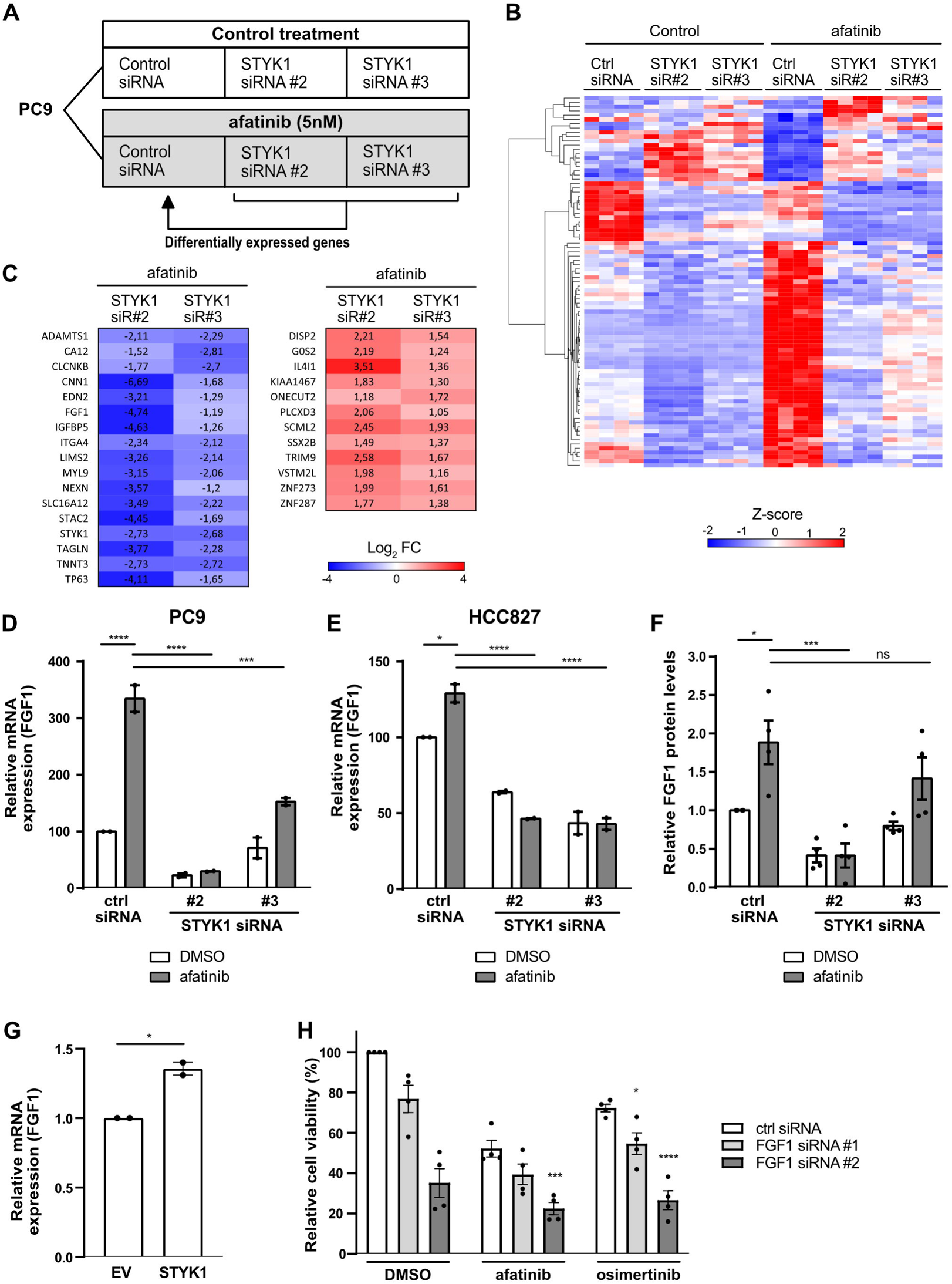
STYK1 knockdown counteracts afatinib-induced increase of FGF1 levels in NSCLC cells. (A) Schematic diagram of the RNAseq experiments performed. PC9 cells were transfected with control or STYK1 siRNAs. Forty-eight hours post-transfection, cells were treated with DMSO (control) or afatinib (5nM) for an additional 24 hours. Differentially expressed genes between STYK1 siRNA + afatinib versus control siRNA + afatinib were determined (|log2FoldChange|= 2, p adj. <0.05) from the RNAseq data. (B) Heatmap of DEG for STYK1 siRNA + afatinib versus control siRNA + afatinib based on Z-scores. (C) Expression of top DEG between STYK1 siRNA + afatinib relative to control siRNA + afatinib as determined by RNAseq. (D & E) Validation by qRT-PCR in PC9 (D) and HCC827 (E) cells of the RNAseq results for FGF1. Data are presented as mean ± SEM from two independent experiments. (F) Validation at the protein level of the differential expression of FGF1, in the experimental conditions of the RNAseq screen, as determined by ELISA. The graph represents mean ± SEM from three independent experiments. (G) PC9 cells were transfected with empty vector (EV) or STYK1 and incubated for 48h. FGF1 mRNA levels were determined by qRT-PCR and plotted as mean ± SEM (n=2). (H) PC9 cells were reverse transfected with non-targeting control or FGF1 siRNAs. 48h after transfection, cells were treated with control (DMSO), afatinib (5nM) or osimertinib (10nM). Cell viability was measured after an additional 48h of incubation. Data are presented as mean ± SEM from four independent experiments. Statistics are shown for FGF1 siRNA compared to control siRNA for the respective treatments.

## Discussion

Numerous strategies have been explored to increase the efficacy of EGFR TKI in EGFR-mutant NSCLC. While some have shown that the efficacy of EGFR TKI can be improved by targeting specific acquired resistance mechanisms, none of these efforts have readily resulted in the betterment of clinical outcomes for patients, except for a modest effect by co-targeting VEGF and EGFR (27).

Here, we uncovered STYK1 as a prime therapeutic target involved in immediate drug tolerance of EGFR mutant NSCLC cells to the EGFR TKI afatinib and osimertinib. STYK1 depletion potentiated EGFR TKI effects on cell viability and colony formation in EGFR mutant NSCLC cells while having minimal effects on EGFR WT cells.

We hypothesize that STYK1 depletion blocks a functional resistance mechanism elicited by EGFR inhibition. Few kinases such as IGF-1R (7), FGFR3 (28), AXL (29) and AURKA (30) have previously been shown to cause drug tolerance to EGFR TKI. Alternative mechanisms associated with drug tolerance also include alteration of chromatin epigenetic state (7), evading apoptosis (31), or interfering with EGFR degradation itself *via* USP13, as we have recently reported (21).

We found that STYK1 overexpression increased colony formation of afatinib-treated cells. This finding supports the role of STYK1 in modulating the sensitivity of NSCLC cells to EGFR TKI and suggests that expression levels of STYK1 could be a valuable biomarker to predict the long-term efficacy of EGFR TKI. Our results agree with previously published studies indicating that increased STYK1 expression correlates with enhanced tumor growth and invasiveness in NSCLC and with poorer prognosis for NSCLC patients, making STYK1 also a potential prognostic marker (14,18,32). Future analyses in EGFR mutant NSCLC patients could evaluate whether the expression of STYK1 can be used as a biomarker to predict progression-free survival and prognosis in response to EGFR inhibitors.

We also found that STYK1 colocalizes with mutated EGFR in NSCLC cells and we proved the existence of a preferential interaction between STYK1 and mutant EGFR. Remarkably, EGFR kinase inhibition disrupted the interaction between STYK1 and mutated EGFR. Colocalization of STYK1 and WT EGFR was previously shown in STYK1-overexpressing HeLa cells (25), especially in early endosomes upon EGF stimulation (25). Overall, these data indicate that STYK1 is recruited to an active EGFR complex, but how precisely the interaction with EGFR affects STYK1 function and *vice versa* remains to be determined. Nevertheless, we speculate that the fraction of STYK1 released from EGFR upon its inhibition is probably the STYK1 sub-population that contributes the most to the drug tolerance mechanisms to EGFR TKI in NSCLC cells.

STYK1 was initially classified as a pseudokinase, with no or weak activity based on the absence of some critical residues which are usually present and conserved in protein kinases. Some groups, however, have reported a residual catalytic activity for STYK1 (13,24). Experiments with a kinase-dead version of STYK1 have confirmed that STYK1 residual kinase activity is required to promote the growth of prostate cancer cells *in vitro* (13), although no direct substrates of STYK1 have been reported until now. In our study, introducing point mutations in STYK1, eliminating its (residual) kinase activity, did not abrogate the enhanced colony formation in STYK1-overexpressing cells treated with afatinib. Our experiments rather unveil a kinase-independent function for STYK1 and imply that the effect triggered by STYK1 in EGFR-inhibited NSCLC cells must be due to a scaffolding role. Interestingly, it has been previously reported that STYK1 can act as a scaffold to enhance the phosphorylation of GSK3β through activated Akt, which likely promotes cancer cell survival (33).

The oncogenic function of STYK1 has been linked to signaling through PI3K/Akt and MAPK pathways in multiple models, including Baf3 cells (9), gallbladder carcinoma (22), and hepatocellular carcinoma (16). However, in EGFR mutant NSCLC cells, we did not observe significant effects on the activation of PI3K/Akt and ERK pathways upon STYK1 downregulation. Also, upon afatinib treatment which severely reduces ERK and Akt activations, STYK1 knockdown did not display any additional effects in the activation levels of these two pathways. One possible explanation is that the activation of the PI3K and MAPK pathways by the mutant EGFR in NSCLC cells does not require an additional modulation by STYK1 or that the modulation is ‘masked’ considering the strong signal promoted by mutant EGFR itself.

Finally, we performed RNAseq experiments aimed to determine downstream effectors of STYK1 in NSCLC cells and to unravel the underlying mechanisms sustaining the potentiation of the combined treatment with EGFR inhibitors and knockdown of STYK1. By doing so, we identified Fibroblast Growth Factor 1 (FGF1) as a downstream effector of STYK1 in NSCLC cells. We found that a short treatment with afatinib caused an acute upregulation of FGF1 in EGFR mutant lung cancer cells. This observation agrees with the findings by Raoof *et al*. reporting upregulation of multiple FGF ligands, including FGF1, in the drug tolerant mechanisms arising in NSCLC cells treated with EGFR TKI for two weeks (28). FGF1 is a potent cytokine with a major role in cell proliferation, differentiation, and survival, and is expressed in various cell types, including NSCLC cells (34). In lung tissue from NSCLC patients, high levels of FGF1 present in both the cytoplasm and nuclei correlated with lower overall survival of NSCLC patients. FGF1 can function through FGF receptors (FGFR) in an autocrine or paracrine manner, but it can also have intracellular FGFR-independent signaling routes (35). FGF1 binds to all members of the FGFR family, resulting in signaling through MAPK, PI3K/Akt and STAT pathways. FGFR signaling has been associated with acquired resistance to EGFR inhibitors and FGFR TKI inhibition in combination with EGFR TKI are known to diminish resistance to EGFR TKI (36–38). Furthermore, it has been demonstrated that FGFR signaling is implicated in the survival of drug tolerant cells (28). Since we observed that STYK1 knockdown did not alter PI3K or MAPK activation in NSCLC cells, it is unlikely that the classical FGFR-dependent signaling pathways are involved in the mechanism we identified. Remarkably, FGF1 effects can also be mediated via intracrine pathways, and the intracellular pools of non-secreted FGF ligands can have mitotic and anti-apoptotic functions independent of FGFR binding (40–42).

In summary, our data provide a strong rationale for further exploring a combined targeting therapy involving STYK1 and EGFR in EGFR-mutant NSCLC in the pre-clinical setting. FGF1 seems to be an important effector in this model, presumably through an intracrine pathway. As such, current FGFR inhibitors may not be suited to target the intracrine effects of FGF1 in NSCLC cells. In that context, the development of STYK1 inhibitors or degraders (that also target the kinase-independent functions of STYK1) could emerge as potentially interesting precision medicine drugs in the future.

## Materials and methods

### Cell cultures and drugs

The human NSCLC cell lines PC9 (RIDD: CVCL_B260; Sigma-Aldrich, Saint-Louis, MO, USA) and HCC827 (RIDD: CVCL_2063; ATCC, Manassas, VA, USA), harboring an exon 19 deletion (ΔE746-A750) in EGFR, and H1975 (RIDD: CVCL_1511; ATCC), harboring EGFR L858R/T790M mutations, were cultured in RPMI-1640 medium (Thermo Fisher, Waltham, MA, USA). Human embryonic kidney 293T (HEK293T) (RIDD: CVCL_0063; ATCC) cells and BEAS-2B (RIDD: CVCL_0168; kindly provided by Prof. Didier Cataldo, Université de Liège, Belgium) cells were cultured in DMEM. All media were supplemented with 10% fetal bovine serum (FBS) and 1% penicillin-streptomycin. The cells were kept in a humidified incubator at 37°C and 5% CO_2_ and were passaged every three days using Versene and 0.25% Trypsin. All cell lines were authenticated using STR profiling. Afatinib (S1011, Selleckchem, Houston, TX, USA) and osimertinib (S7297, Selleckchem) were used as EGFR tyrosine kinase inhibitors (TKI) and used at final concentrations mentioned in the respective figure legends. Stock solutions for all drugs were prepared in dimethyl-sulphoxide (DMSO). The final concentration of DMSO in all the assays was <0.1%.

### Kinome siRNA screen

An arrayed, custom-produced siRNA library, consisting of siRNA pools (4 siRNA sequences per gene) targeting 659 known and putative human kinases, was employed (ON-target plus SMART pool custom made, Horizon, Cambridge, UK). A final siRNA concentration of 10nM was used, and the experiments were performed in duplicates. The viability assay itself was performed as described in the cell viability section. Analysis of the viability results was based on a robust Z-score (43). The Z-score = ([mean viability ratio of afatinib and DMSO (siRNA pool)] − [mean viability ratio afatinib and DMSO of the 384 well-plate])/mean absolute deviation (MAD). The MAD was calculated as follows: MAD = 1.4826 (k is a constant scale factor for normal distribution) × absolute value ([mean viability ratio of afatinib and DMSO (siRNA pool)] − [mean viability ratio afatinib and DMSO of the 384-well plate]).

### Plasmids and siRNA transfections

Four different siRNA sequences (J-003113-09, J-003113-10, J-003113-11 and J-003113-12) targeting STYK1 and two FGF1 siRNA sequences (J-011172-05 and J-011172-08) were purchased from Horizon. The cells were transiently transfected with the siRNAs using Lipofectamine® RNAiMAX (13778-030, Thermo Fisher) according to the manufacturer’s protocol. Final siRNA concentrations were 6nM.

pcDNA3.1-NOK-HA, pcDNA3.1-NOK-FLAG and pEF-BOS-NOK were kindly provided by Prof. Li Liu (Institute of Basic Medical Sciences, Chinese Academy of Medical Sciences & School of Basic Medicine, Peking Union Medical College, Beijing, China). Site-directed mutagenesis on these constructs was performed to convert them to STYK1 (by introducing the P203L point mutation (10)). After PCR amplification, the template DNA was removed by DpnI digestion. To create pLentipuro-STYK1-eGFP, eGFP-tagged STYK1 was PCR-amplified from the pEF-BOS-STYK1 vector and subcloned in the pLentipuro backbone (39481, Addgene, Watertown, MA, USA) using XbaI and MluI restriction enzymes. The K147A and Y191F mutants in pLentipuro STYK1-eGFP were generated by site-directed mutagenesis. pcDNA4-EGFR-myc-his B was kindly provided by Dr. Yi-Rong Chen (National Health Research Institutes, Taiwan). pcDNA4-EGFR-L858R-myc-his B and pcDNA4-EGFR-Δ746-750-myc-his B were previously generated by site-directed mutagenesis (21). Primers used for cloning and site-directed mutagenesis are listed in Supplemental Table 3. Plasmids inserts and desired mutations were all confirmed by sequencing (Eurofins, Luxembourg). DNA plasmids transfections were performed using Lipofectamine® 2000 (11668-019, Thermo Fisher).

### Lentiviral particles production and cell transduction

To produce the lentiviral vectors (LV), HEK293T cells were plated at 12 × 10^6^ cells per 175cm^2^ flask. The following day, polyethyleneimine (Polysciences, Eppelheim, Germany) was used to transfect the transgene encoding plasmid (37.5μ together with the envelope VSV-G (9µg) and packaging GAG (12.5 µg) and REV (6.25µg) coding plasmids (kindly provided by Prof. Brian Brown, Mount Sinai Icahn School of Medicine, NY, USA). LV-containing supernatants were collected on days 2 and 3 after transfection and subsequently filtered through a 0.22µm filter. The target cells were transduced using the LV-containing medium with the addition of 10µg/ml protamine sulfate (P4020, Sigma-Aldrich). Lentiviral particles containing pLentipuro-STYK1-eGFP WT/K147A/Y191F were used to infect PC9 and HCC827 cells to generate stable STYK1-eGFP (WT or mutant) overexpressing cells. Seventy-two hours after infection, the transduced PC9 and HCC827 cells were selected with respectively 4 and 1µg/ml puromycin. Stable knockdown of STYK1 in PC9 cells was achieved by lentiviral transduction using PLKO.1-puro-STYK1 shRNA constructs (TRCN0000001742, TRCN0000001744 and TRCN0000001746, Horizon), followed by 1 µg/ml puromycin selection.

### Cell viability

Cells were reverse transfected with either control, STYK1, or FGF1 siRNAs and plated at a density of 500-1 000 cells in a 384-well plate. After culturing for 48 hours, EGFR TKI (afatinib, osimertinib) were added to the cells at the indicated doses. After an incubation time of 96 hours, cell viability was measured using the CellTiter-Glo® luminescent kit (G7571, Promega, Madison, WI, USA) with the help of a Spectromax® M3 (Molecular Devices, San Jose, CA, USA).

### Apoptosis analysis by flow cytometry

Cells were reverse transfected with either control or STYK1 siRNAs, seeded in 6-well plates, and the indicated drugs were added after 48 hours. The cells were ultimately harvested after 96 hours of incubation. After harvesting, the cells were washed, resuspended in Annexin V binding buffer (556454, BD Biosciences, Franklin Lakes, NJ, USA), and then stained with 5 µl APC-Annexin V (550475, BD Biosciences) and 5 µl 7-AAD (559925, BD Biosciences) according to the manufacturer’s instructions. The measurements were done by a FACScanto flow cytometer (BD Biosciences). FACS data were analyzed for percentages of viable cells (AnnexinV-/7-AAD-), early-apoptotic cells (AnnexinV+/7-AAD-), and late apoptotic cells (AnnexinV+/7-AAD+) using *BD FACSDiva 8.0.1* (RRID:SCR_001456).

### Quantitative real-time PCR

Total mRNA extraction was performed using the Nucleospin RNA plus kit (740984, Macherey-Nagel, Bethlehem, PA, USA) according to the manufacturer’s protocol. First-strand cDNA was synthesized using SuperScript™ II Reverse Transcriptase (18064014, Thermo Fisher). Target and reference genes were quantified in duplicates on a LightCycler®480 (Roche, Basel, Switzerland) using SYBR Green I Mastermix (04707516001, Roche). The geometric mean of housekeeping genes TBP and SDHA was used for the normalization of input cDNA. Primer sequences are listed in Supplemental Table 3.

### Western blotting

Cells were lysed using a buffer containing 1% Triton X-100, 20mM Tris-HCl (pH=7.5), 150mM NaCl, 1mM EDTA, 2.5mM sodium-pyrophosphate, and 1mM sodium orthovanadate, supplemented with 1% phosphatase inhibitors (P8340, Sigma-Aldrich) and 1% protease inhibitors (P5726, Sigma-Aldrich). Protein concentrations were calculated using the Bradford protein assay kit (Bio-Rad, Hercules, CA, USA). Equal amounts of proteins (10-30µg) were loaded and separated by SDS-PAGE using 10%-15% resolving acrylamide gels. Proteins were transferred overnight onto nitrocellulose membranes. After blocking the membranes with 5% non-fat milk, primary antibodies diluted in 5% bovine serum albumin containing Tris-buffered saline Tween-20 (TBST) were incubated overnight at 4°C. The appropriate secondary infrared-conjugated antibodies (LI-COR, Lincoln, NE, USA) were incubated for one hour at room temperature protected from light. Detection was performed using the LI-COR Biosciences Odyssey® Fc Imaging System and analyzed with the Image Studio™ software (LI-COR). Western blots are representative images from experiments that were repeated at least two times.

Primary antibodies for western blot used in the study are: phospho-EGFR (Tyr1068, #3777, RRID:AB_2096270), total ERK (#4695, RRID:AB_390779), phospho-ERK1/2 (Thr202/Tyr204, #4370, RRID:AB_2315112), total Akt (#9272, RRID:AB_329827), and phospho-Akt (Ser473, #4058, RRID:AB_331168) from Cell Signaling Technology (Danvers, MA, USA); total EGFR (AMAb90816, RRID:AB_2665679), β-actin (A1978, RRID:AB_476692) and α-tubulin (T9026, RRID:AB_477593) from Sigma-Aldrich; and STYK1 (Ab97451 (RRID:AB_10679828), Abcam, Cambridge, UK and PA521695 (RRID:AB_11154125), Thermo Fisher).

### Co-immunoprecipitation

Cells were lysed 24 hours after transfection with the indicated plasmids using the Triton X-100 containing lysis buffer. Equal amounts of proteins were subjected to immunoprecipitation, and 20 µg of each condition were saved as control inputs. Immunoprecipitation was performed for three hours at 4°C using anti-EGFR antibodies (GRO1, Sigma-Aldrich) combined with protein G coupled Sepharose (17-0618-01, GE Healthcare, Chicago, IL, USA) or anti-FLAG coupled agarose (A2220, Sigma-Aldrich). After incubation, the beads were washed three times using ice-cold PBS. Proteins were eluted by the addition of a 2X SDS-containing loading buffer and subjected to Western blotting.

### Soft agar assay

A total of 10,000 cells were suspended in 1.5mL of 0.3% agar supplemented with complete RPMI-1640 medium containing the appropriate concentrations of afatinib or osimertinib and layered on top of a 0.5% base agar in a 6-well plate. Every week, a freshly prepared drug-containing medium was added on top. After a total incubation time of three weeks, nitro blue tetrazolium chloride was added to visualize the cell colonies. Twenty-four hours later, the wells were photographed, and the colonies were quantified with the help of the OpenCFU software (44). Experiments were performed in triplicates.

### Confocal Microscopy

Stable STYK1-eGFP overexpressing HCC827 and empty vector control cells were seeded in a black Microclear 96-well plate (655090, Greiner, Frickenhausen, Germany). At 90% confluency, the cells were fixed using 4% paraformaldehyde for 15 min at 4°C and subsequently permeabilized with 0.25% Triton X-100 in PBS for 15 min. Subsequent washes were performed with 0.05% Tween-20 in PBS. Primary antibody for EGFR (GRO1, Sigma-Aldrich) was diluted in 0.05% Tween-20, 1% BSA and 22.5mg/mL Glycine, and was incubated for one hour, followed by one hour staining with an AlexaFluor 647-labeled secondary antibody. Nuclei were stained with Hoechst3342. Images were acquired using a ZEISS LSM710 NLO Confocal microscope using the ZEN 2009 software (Carl Zeiss, Oberkochen, Germany).

### RNA sequencing

Total mRNA extraction was performed using the Nucleospin RNA plus kit (740984, Macherey-Nagel) according to the manufacturer’s protocol. Four biological repeats for each condition were processed by the BRIGHTcore platform (Brussels University Alliance, VUB-ULB, Brussels, Belgium). RNA sequencing libraries were constructed using the TruSeq RNA Library kit (Illumina, San Diego, CA, USA) and sequenced on an Illumina NovaSeq 6000 instrument. The DeSeq2 tool (RRID:SCR_015687) was used in R to determine the differentially expressed genes (DEG) (45). The cut-off values for DEG selection were ≥ 2-fold change and p<0.05. KEGG analysis was performed using KOBAS3.0 (http://kobas.cbi.pku.edu.cn/kobas3) (RRID:SCR_006350) (46), and a heatmap was created by the Morpheus tool (https://software.broadinstitute.org/morpheus) (RRID:SCR_017386).

### ELISA

Cells were lysed using Triton X-100 containing lysis buffer. 20µg of total cell lysates were loaded in the FGF1 coated wells. The FGF1 ELISA was performed with the Quantikine FGF1 ELISA kit (DFA00B, R&D systems, Minneapolis, MN, USA) according to the manufacturer’s instructions.

### Statistics

Graphs are represented by means and SEM. Statistical analysis was performed using GraphPad Prism (RRID:SCR_002798). Sample size was not predetermined using statistical methods, but based on previous experience. The two-tailed *student’s* t-test was used to assess the significance of STYK1 and FGF1 knockdown (qRT-PCR) and other data consisting of two groups. One-way ANOVA was used to determine significance in relative number of colonies upon STYK1 (WT or mutant) overexpression. Two-way ANOVA with Sidak posthoc analysis was performed on all data with two independent variables. P-value less than 0.05 was considered statistically significant (*<0.05; **<0.01; ***<0.001; and ****<0.0001).

## Acknowledgments

We thank BrightCore (UZB/VUB) for the RNA sequencing experiments; Profs. Karine Breckpot and Cleo Goyvaerts (VUB) for the use of the lentiviral core facility; Prof. Luc Bouwens and Eddy Himpe (VUB) for the use of the Confocal microscope; and Prof. Luc Leyns (VUB) for access to his laboratory equipment.

## Data availability statement

The datasets used and/or analyzed during the current study are available from the corresponding authors on reasonable request.

