## Supplemental Information for "The EGFR-STYK1-FGF1 axis sustains functional drug tolerance to EGFR inhibitors in EGFR-mutant non-small cell lung cancer"

**Table of contents:**

Supplemental Figure 1 - STYK1 knockdown potentiates the EGFR TKI effects on cell viability and apoptosis in NSCLC cells	2

Supplemental Figure 2 - STYK1 interacts with mutant EGFR and STYK1 knockdown does not alter downstream EGFR signaling in NSCLC cells	4

Supplemental Figure 3 – RNA sequencing KEGG pathway analysis and validation of top hits	5

Supplemental Table 1 – Kinome RNAi screen data	7 & data provided as a separate file

Supplemental Table 2 – RNA sequencing data	7 & data provided as a separate file

Supplemental Table 3 – Primer sequences	8

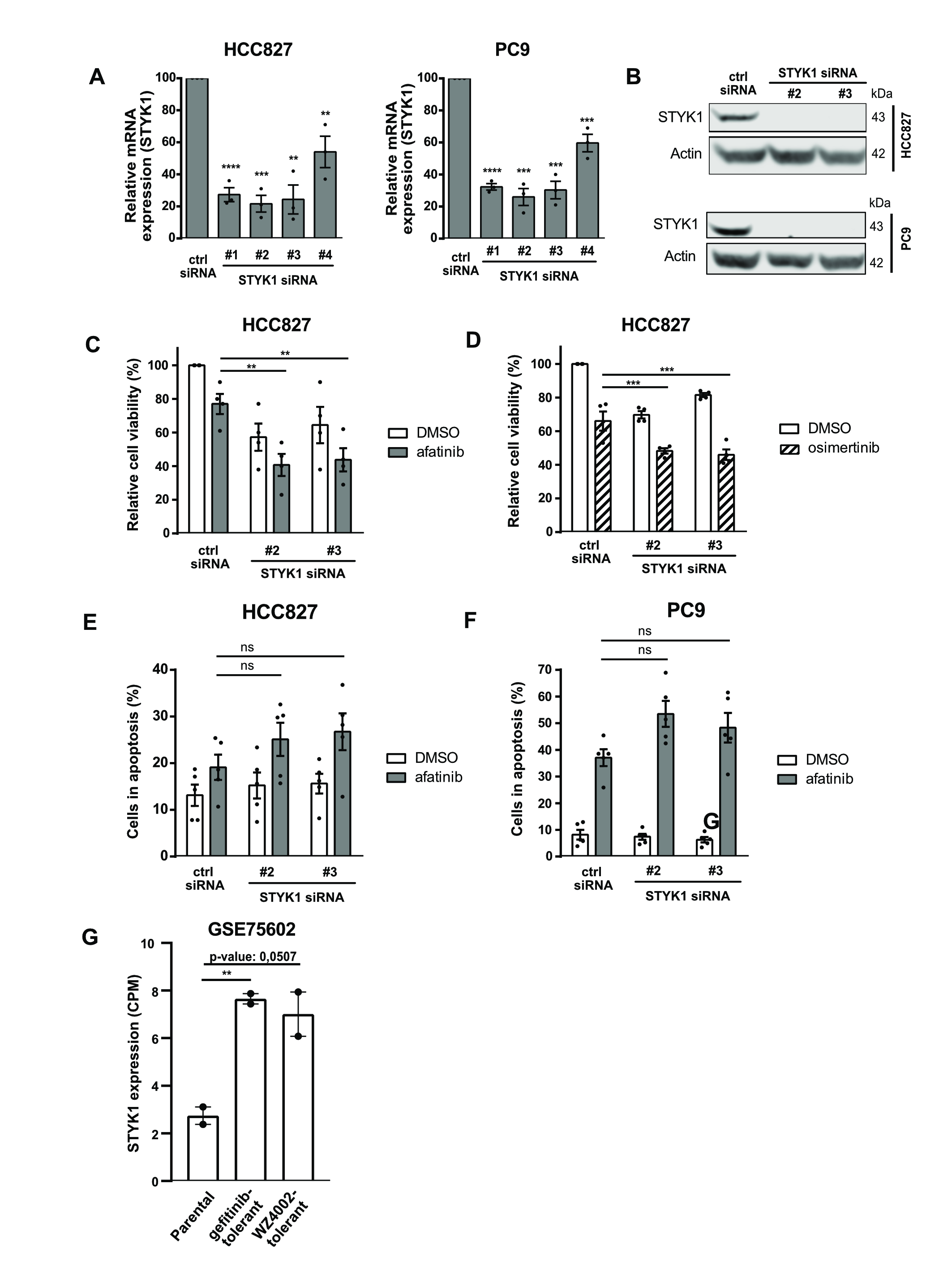

***Supplemental Figure 1: STYK1 knockdown potentiates the EGFR TKI effects on cell viability and apoptosis in NSCLC cells.*** *(A) HCC827 and PC9 were reverse transfected with control or individual STYK1 siRNAs. qRT-PCR analysis was performed after 24 hours to determine knockdown efficiency (mean ± SEM, n=3). (B) Western blotting for STYK1 72 hours after reverse transfection of the indicated siRNAs. (C & D) HCC827 cells were reverse transfected with control or indicated STYK1 siRNAs, and 48 hours post-transfection 5nM afatinib (C) or 10nM osimertinib (D) were added. Cell viability was measured after 96 hours. Data are shown relative to the control siRNA-DMSO condition and represented as mean ± SEM, with at least four independent repeats. (E-F) HCC827 and PC9 cells were reverse transfected with control or STYK1 siRNAs and 48 hours post-transfection 5nM afatinib were added for an additional 24 hours. The number of apoptotic cells was determined using 7‑AAD/Annexin V-APC staining (mean ± SEM, n=5). (G) STYK1 expression data (counts per million = CPM) in parental PC9 cells versus gefitinib- or WZ4002-drug tolerant PC9 cells from the publicly available dataset GSE75602.*

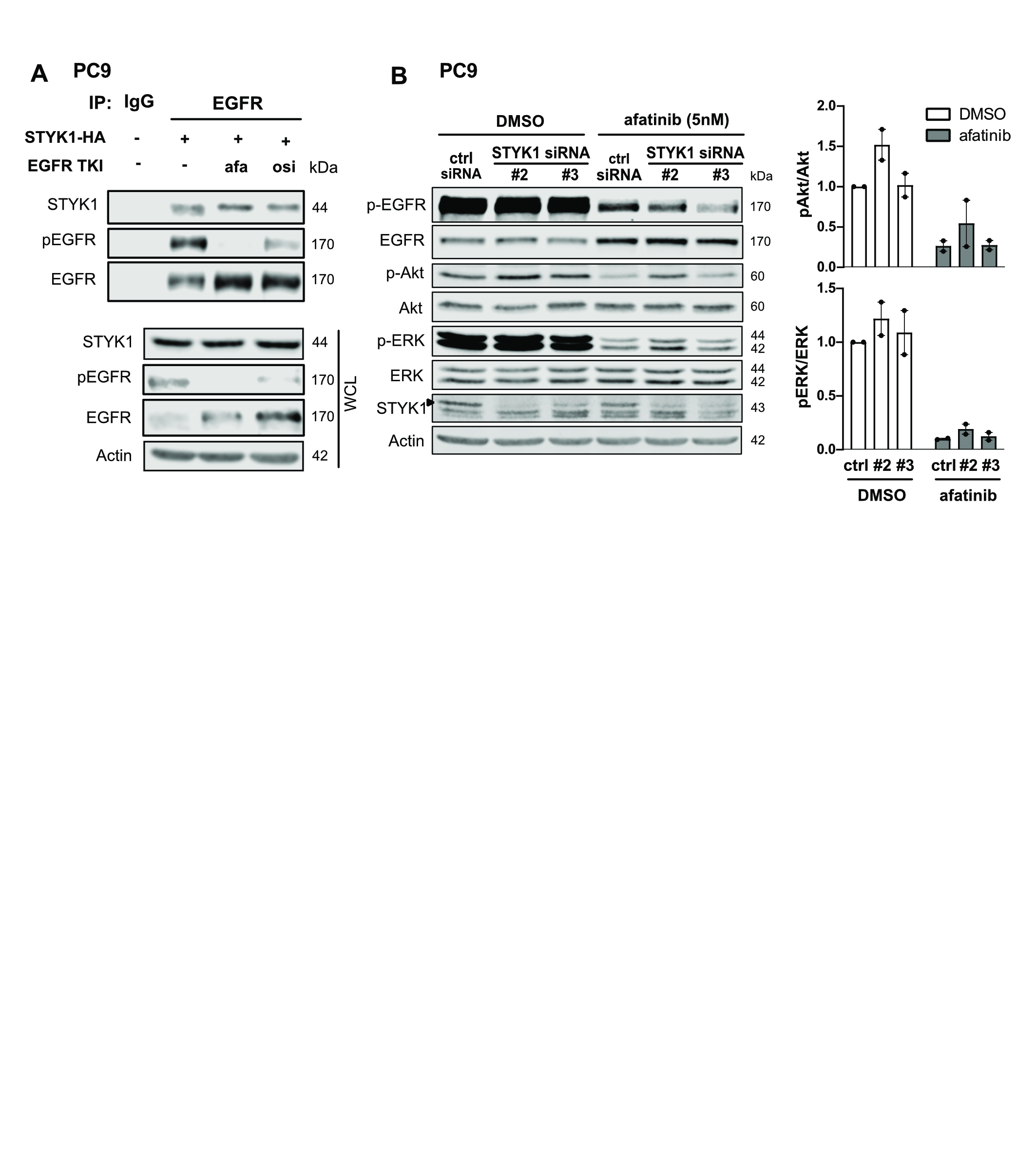

***Supplemental Figure 2: STYK1 interacts with mutant EGFR and STYK1 knockdown does not alter downstream EGFR signaling.*** *(A) STYK1-HA was transfected in PC9 cells. After an overnight treatment with afatinib (10nM) or osimertinib (10nM) and a total incubation of 24 hours post-transfection, lysates were prepared and subjected to immunoprecipitation using anti-EGFR or IgG control antibodies. Immunoprecipitates and WCL were immunoblotted with the indicated antibodies. (B) PC9 cells were reverse transfected with control or STYK1 siRNAs and 48 hours post-transfection 5nM afatinib were added for an additional 24 hours. Total lysates were immunoblotted with indicated antibodies. pAkt/Akt and pERK/ERK levels were quantified by the LI-COR Odyssey software. Normalized values of two independent experiment are shown as mean ± SEM.*

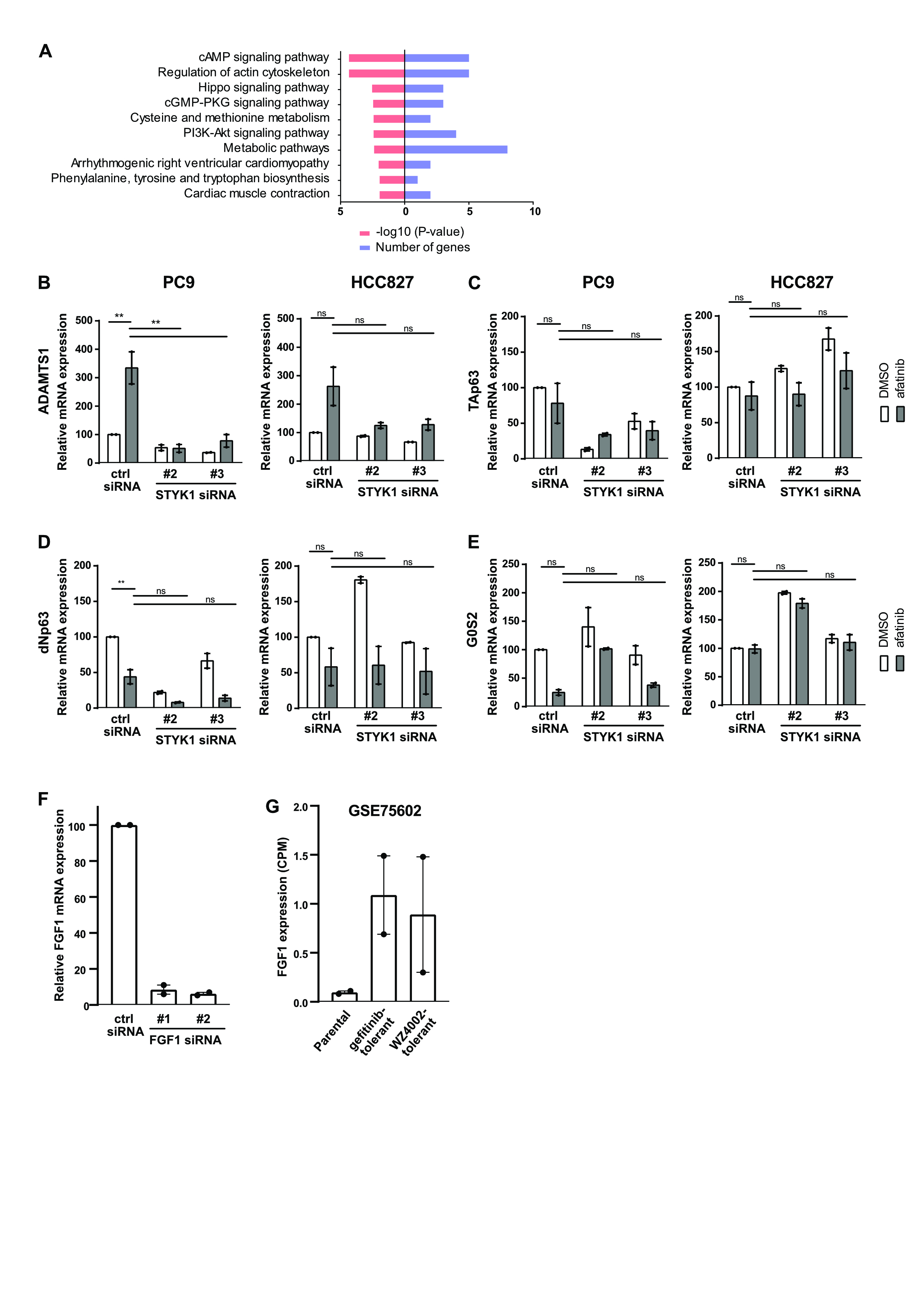

***Supplemental Figure 3:*** ***RNA sequencing KEGG pathway analysis and validation of top hits.*** *(A) KEGG pathway analysis results from the RNAseq data. Red column shows p-value (Log10), and blue column shows number of hit genes. (B-E) Further validation of the RNAseq data by qRT-PCR in PC9 and HCC827 cells upon control or STYK1 siRNA transfection with or without afatinib for ADAMTS1 (B), TAp63 (C), dNp63 (D), and G0S2 (E). Data are presented as mean ± SEM from two independent experiments. (F) PC9 cells were reverse transfected with control or individual FGF siRNAs. qRT-PCR analysis was performed after 24 hours to determine knockdown efficiency (mean ± SEM, n=2). (G) FGF1 expression data (counts per million = CPM) in parental PC9 cells versus gefitinib- or WZ4002-drug tolerant PC9 cells from the publicly available dataset GSE75602.*

***Supplemental Table 1: Kinome RNAi screen data.*** *Data are plate-normalized to the median of the control siRNA-DMSO wells. For each siRNA pool there are 4 wells: DMSO1; DMSO2; AFA1; and AFA2. The average (av) and standard deviation (sd) of duplicates was calculated per siRNA pool. The ratio avAFA/avDMSO and finally a robust Z score were calculated to represent the data.*

*Data are provided as a separate Excel-file: “SUPPL.TABLE1 RNAi SCREEN”*

***Supplemental Table 2: RNA sequencing data.*** *For each gene (identified by Ensembl Gene ID and Gene symbol) the basemean, log2 fold change, and adjusted p-value are shown for STYK1 siRNA2-afatinib and STYK1 siRNA3-afatinib versus control siRNA-afatinib.*

*Data are provided as a separate Excel-file: “SUPPL.TABLE2 RNASEQ”*

| Cloning | XbaI-STYK1-GFP FW | 5'-ATCTCTAGAATGGGCATGACACGGATG-3' |
| --- | --- | --- |
| Cloning | MluI-STYK1-GFP RV | 5'-ATCACGCGTTTACTTGTACAGCTCGTCCA-3' |
| Mutagenesis | NOK P203L (STYK1) | 5'-GGTCCAGAGAAAGCTGAGCAGGTCCCCCTGGG-3' |
| Mutagenesis | NOK P203L (STYK1) | 5'-CCCAGGGGGACCTGCTCAGCTTTCTCTGGACC-3' |
| Mutagenesis | K147A FW | 5'-AGGCCCAAGAGTGTTATTCTCGCGGCTTTAAAAGAACCAGCTG-3' |
| Mutagenesis | K147A RV | 5'-CAGCTGGTTCTTTTAAAGCCGCGAGAATAACACTCTTGGGCTT-3' |
| Mutagenesis | Y191F FW | 5'-TGAAAAGCTGCCACTCTTTATGGTGTTGGAGGATG-3' |
| Mutagenesis | Y191F RV | 5'-CATCCTCCAACACCATAAAGAGTGGCAGCTTTTCA-3' |
| qRT-PCR | STYK1 FW | 5'-GAAGTTTACACCCGAGGGGC-3' |
| qRT-PCR | STYK1 RV | 5'-GGTGCTCCTAGAGTCACCATC-3' |
| qRT-PCR | SDHA FW | 5'-TGGGAACAAGAGGGCATCTG-3' |
| qRT-PCR | SDHA RV | 5'-CCACCACTGCATCAAATTCATG-3' |
| qRT-PCR | TBP FW | 5'-CACGAACCACGGCACTGATT-3' |
| qRT-PCR | TBP RV | 5'-TTTTCTTGCTGCCAGTCTGGAC-3' |
| qRT-PCR | FGF1 FW | 5'-GGAGCGACCAGCACATTCAG-3' |
| qRT-PCR | FGF1 RV | 5'-CCGTATAAAAGCCCGTCGGT-3' |
| qRT-PCR | ADAMTS1 FW | 5'-AGCCCATGAATTAGGCCACG-3' |
| qRT-PCR | ADAMTS1 RV | 5'-TTGACGCCATCATGTGGGAA-3' |
| qRT-PCR | dNp63 FW | 5'-AGCCAGAAGAAAGGACAGCA-3' |
| qRT-PCR | dNp63 RV | 5'-CAGGTTCGTGTACTGTGGCT-3' |
| qRT-PCR | TAp63 FW | 5'-TGTATCCGCATGCAGGACT-3' |
| qRT-PCR | TAp63 RV | 5'-CTGTGTTATAGGGACTGGTGGAC-3' |
| qRT-PCR | G0S2 FW | 5'-GCCGTGCCACTAAGGTCATT-3' |
| qRT-PCR | G0S2 RV | 5'-GATCAGCTCCTGGACCGTTT-3' |

**Supplemental Table 3: Primers sequences used in this study.** FW= forward primer; RV=reverse primer.
