## Supplemental Table 1 for "The EGFR-STYK1-FGF1 axis sustains functional drug tolerance to EGFR inhibitors in EGFR-mutant non-small cell lung cancer"

| Index | PlateID | GeneID | GeneSymbol | category | nor_PC9_DMSO1 | nor_PC9_DMSO2 | nor_PC9_AFA1 | nor_PC9_AFA2 | av PC9DMSO | avPC9AFA | sdPC9DMSO | sdPC9AFA | ratio AFA/DMSO | onlyS1 | median plate | abs dev1 | mad*1.4826_1 | Z*A/D |
|---|---|---|---|---|---|---|---|---|---|---|---|---|---|---|---|---|---|---|
| 1 | Dha_Kin_pool_1 | 6011 | GRK1 | S | 0,52 | 0,66 | 0,30 | 0,36 | 0,59 | 0,33 | 0,10 | 0,04 | 0,56 | 0,56 | 0,62 | 0,06 | 0,14 | -0,40 |
| 2 | Dha_Kin_pool_1 | 80271 | ITPKC | S | 0,50 | 0,49 | 0,15 | 0,16 | 0,49 | 0,15 | 0,01 | 0,01 | 0,31 | 0,31 | 0,62 | 0,31 | 0,14 | -2,17 |
| 3 | Dha_Kin_pool_1 | 2044 | EPHA5 | S | 1,00 | 0,85 | 0,61 | 0,56 | 0,92 | 0,59 | 0,10 | 0,03 | 0,64 | 0,64 | 0,62 | 0,02 | 0,14 | 0,14 |
| 4 | Dha_Kin_pool_1 | 91807 | MLCK | S | 1,00 | 0,85 | 0,63 | 0,62 | 0,93 | 0,63 | 0,11 | 0,01 | 0,68 | 0,68 | 0,62 | 0,06 | 0,14 | 0,43 |
| 5 | Dha_Kin_pool_1 | 5681 | PSKH1 | S | 1,05 | 0,79 | 0,55 | 0,49 | 0,92 | 0,52 | 0,19 | 0,04 | 0,57 | 0,57 | 0,62 | 0,05 | 0,14 | -0,35 |
| 6 | Dha_Kin_pool_1 | 6795 | AURKC | S | 1,24 | 1,04 | 0,61 | 0,63 | 1,14 | 0,62 | 0,14 | 0,01 | 0,55 | 0,55 | 0,62 | 0,07 | 0,14 | -0,48 |
| 7 | Dha_Kin_pool_1 | 5581 | PRKCE | S | 0,85 | 0,82 | 0,48 | 0,53 | 0,83 | 0,50 | 0,02 | 0,04 | 0,60 | 0,60 | 0,62 | 0,01 | 0,14 | -0,10 |
| 8 | Dha_Kin_pool_1 | 203 | AK1 | S | 1,02 | 1,08 | 0,50 | 0,50 | 1,05 | 0,50 | 0,04 | 0,00 | 0,48 | 0,48 | 0,62 | 0,14 | 0,14 | -0,96 |
| 9 | Dha_Kin_pool_1 | 140803 | TRPM6 | S | 0,86 | 0,66 | 0,32 | 0,33 | 0,76 | 0,33 | 0,14 | 0,01 | 0,43 | 0,43 | 0,62 | 0,18 | 0,14 | -1,29 |
| 10 | Dha_Kin_pool_1 | 1195 | CLK1 | S | 1,06 | 0,91 | 0,54 | 0,48 | 0,98 | 0,51 | 0,11 | 0,04 | 0,52 | 0,52 | 0,62 | 0,10 | 0,14 | -0,67 |
| 11 | Dha_Kin_pool_1 | 6199 | RPS6KB2 | S | 0,81 | 0,91 | 0,47 | 0,53 | 0,86 | 0,50 | 0,08 | 0,04 | 0,58 | 0,58 | 0,62 | 0,03 | 0,14 | -0,24 |
| 12 | Dha_Kin_pool_1 | 5218 | PFTK1 | S | 1,11 | 1,08 | 0,49 | 0,54 | 1,09 | 0,51 | 0,02 | 0,03 | 0,47 | 0,47 | 0,62 | 0,15 | 0,14 | -1,02 |
| 13 | Dha_Kin_pool_1 | 8382 | NME5 | S | 1,15 | 0,91 | 0,61 | 0,52 | 1,03 | 0,56 | 0,17 | 0,06 | 0,55 | 0,55 | 0,62 | 0,07 | 0,14 | -0,48 |
| 14 | Dha_Kin_pool_1 | 6714 | SRC | S | 0,74 | 0,67 | 0,49 | 0,39 | 0,71 | 0,44 | 0,05 | 0,08 | 0,62 | 0,62 | 0,62 | 0,01 | 0,14 | 0,05 |
| 15 | Dha_Kin_pool_1 | 6787 | NEK4 | S | 0,92 | 0,86 | 0,63 | 0,55 | 0,89 | 0,59 | 0,05 | 0,06 | 0,66 | 0,66 | 0,62 | 0,05 | 0,14 | 0,34 |
| 16 | Dha_Kin_pool_1 | 53944 | CSNK1G1 | S | 0,73 | 0,65 | 0,17 | 0,20 | 0,69 | 0,19 | 0,06 | 0,02 | 0,28 | 0,28 | 0,62 | 0,34 | 0,14 | -2,39 |
| 17 | Dha_Kin_pool_1 | 83732 | RIOK1 | S | 0,99 | 0,94 | 0,45 | 0,39 | 0,97 | 0,42 | 0,04 | 0,04 | 0,43 | 0,43 | 0,62 | 0,18 | 0,14 | -1,28 |
| 18 | Dha_Kin_pool_1 | 207 | AKT1 | S | 0,68 | 0,38 | 0,14 | 0,16 | 0,53 | 0,15 | 0,21 | 0,02 | 0,28 | 0,28 | 0,62 | 0,34 | 0,14 | -2,36 |
| 19 | Dha_Kin_pool_1 | 54861 | SNRK | S | 1,24 | 1,06 | 0,47 | 0,58 | 1,15 | 0,52 | 0,12 | 0,08 | 0,45 | 0,45 | 0,62 | 0,16 | 0,14 | -1,14 |
| 20 | Dha_Kin_pool_1 | 84446 | KIAA1811 | S | 1,18 | 0,89 | 0,49 | 0,57 | 1,03 | 0,53 | 0,21 | 0,06 | 0,51 | 0,51 | 0,62 | 0,10 | 0,14 | -0,72 |
| 21 | Dha_Kin_pool_1 | 5164 | PDK2 | S | 0,62 | 0,50 | 0,32 | 0,35 | 0,56 | 0,33 | 0,09 | 0,02 | 0,59 | 0,59 | 0,62 | 0,02 | 0,14 | -0,17 |
| 22 | Dha_Kin_pool_1 | 85366 | MYLK2 | S | 0,80 | 0,99 | 0,50 | 0,45 | 0,90 | 0,47 | 0,13 | 0,03 | 0,53 | 0,53 | 0,62 | 0,09 | 0,14 | -0,61 |
| 23 | Dha_Kin_pool_1 | E | E | D | 0,90 | 0,83 | 0,84 | 1,05 | 0,86 | 0,94 | 0,05 | 0,15 | 1,09 | | 0,62 | | 0,14 | 3,35 |
| 24 | Dha_Kin_pool_1 | E | E | K | 0,03 | 0,03 | 0,04 | 0,04 | 0,03 | 0,04 | 0,00 | 0,00 | 1,38 | | 0,62 | | 0,14 | 5,39 |
| 25 | Dha_Kin_pool_1 | 79858 | NEK11 | S | 0,93 | 0,73 | 0,44 | 0,54 | 0,83 | 0,49 | 0,14 | 0,07 | 0,59 | 0,59 | 0,62 | 0,03 | 0,14 | -0,18 |
| 26 | Dha_Kin_pool_1 | 57551 | KIAA1361 | S | 0,86 | 0,89 | 0,54 | 0,64 | 0,87 | 0,59 | 0,02 | 0,07 | 0,68 | 0,68 | 0,62 | 0,06 | 0,14 | 0,44 |
| 27 | Dha_Kin_pool_1 | 150094 | SNF1LK | S | 0,92 | 0,88 | 0,62 | 0,54 | 0,90 | 0,58 | 0,03 | 0,06 | 0,64 | 0,64 | 0,62 | 0,03 | 0,14 | 0,20 |
| 28 | Dha_Kin_pool_1 | 9113 | LATS1 | S | 0,86 | 0,70 | 0,42 | 0,48 | 0,78 | 0,45 | 0,11 | 0,04 | 0,57 | 0,57 | 0,62 | 0,04 | 0,14 | -0,29 |
| 29 | Dha_Kin_pool_1 | 5214 | PFKP | S | 1,08 | 0,79 | 0,61 | 0,75 | 0,93 | 0,68 | 0,20 | 0,10 | 0,73 | 0,73 | 0,62 | 0,11 | 0,14 | 0,78 |
| 30 | Dha_Kin_pool_1 | 5592 | PRKG1 | S | 1,05 | 0,94 | 0,58 | 0,60 | 0,99 | 0,59 | 0,08 | 0,02 | 0,59 | 0,59 | 0,62 | 0,02 | 0,14 | -0,15 |
| 31 | Dha_Kin_pool_1 | 55781 | RIOK2 | S | 0,82 | 0,70 | 0,40 | 0,43 | 0,76 | 0,42 | 0,08 | 0,02 | 0,55 | 0,55 | 0,62 | 0,07 | 0,14 | -0,50 |
| 32 | Dha_Kin_pool_1 | 2475 | FRAP1 | S | 0,66 | 0,81 | 0,44 | 0,47 | 0,73 | 0,45 | 0,11 | 0,02 | 0,62 | 0,62 | 0,62 | 0,00 | 0,14 | 0,03 |
| 33 | Dha_Kin_pool_1 | 5157 | PDGFRL | S | 0,59 | 0,55 | 0,14 | 0,15 | 0,57 | 0,15 | 0,03 | 0,00 | 0,26 | 0,26 | 0,62 | 0,36 | 0,14 | -2,53 |
| 34 | Dha_Kin_pool_1 | 122481 | AK7 | S | 0,80 | 0,86 | 0,51 | 0,42 | 0,83 | 0,47 | 0,04 | 0,06 | 0,56 | 0,56 | 0,62 | 0,05 | 0,14 | -0,37 |
| 35 | Dha_Kin_pool_1 | 9262 | STK17B | S | 0,72 | 0,85 | 0,52 | 0,61 | 0,79 | 0,57 | 0,09 | 0,07 | 0,72 | 0,72 | 0,62 | 0,11 | 0,14 | 0,74 |
| 36 | Dha_Kin_pool_1 | 8428 | STK24 | S | 0,82 | 0,95 | 0,47 | 0,45 | 0,89 | 0,46 | 0,09 | 0,01 | 0,52 | 0,52 | 0,62 | 0,10 | 0,14 | -0,71 |
| 37 | Dha_Kin_pool_1 | 1607 | DGKB | S | 1,02 | 0,97 | 0,58 | 0,61 | 0,99 | 0,60 | 0,04 | 0,02 | 0,60 | 0,60 | 0,62 | 0,02 | 0,14 | -0,12 |
| 38 | Dha_Kin_pool_1 | 10494 | STK25 | S | 0,68 | 0,99 | 0,58 | 0,66 | 0,83 | 0,62 | 0,22 | 0,05 | 0,75 | 0,75 | 0,62 | 0,13 | 0,14 | 0,91 |
| 39 | Dha_Kin_pool_1 | 8317 | CDC7 | S | 1,00 | 0,75 | 0,38 | 0,40 | 0,87 | 0,39 | 0,18 | 0,02 | 0,45 | 0,45 | 0,62 | 0,17 | 0,14 | -1,18 |
| 40 | Dha_Kin_pool_1 | 818 | CAMK2G | S | 0,92 | 0,72 | 0,49 | 0,50 | 0,82 | 0,49 | 0,14 | 0,00 | 0,60 | 0,60 | 0,62 | 0,01 | 0,14 | -0,10 |
| 41 | Dha_Kin_pool_1 | 54963 | URKL1 | S | 0,31 | 0,23 | 0,13 | 0,16 | 0,27 | 0,14 | 0,06 | 0,02 | 0,53 | 0,53 | 0,62 | 0,09 | 0,14 | -0,61 |
| 42 | Dha_Kin_pool_1 | 4293 | MAP3K9 | S | 0,86 | 0,77 | 0,52 | 0,55 | 0,82 | 0,54 | 0,06 | 0,02 | 0,66 | 0,66 | 0,62 | 0,04 | 0,14 | 0,29 |
| 43 | Dha_Kin_pool_1 | 10645 | CAMKK2 | S | 0,81 | 0,71 | 0,49 | 0,48 | 0,76 | 0,48 | 0,07 | 0,01 | 0,64 | 0,64 | 0,62 | 0,02 | 0,14 | 0,15 |
| 44 | Dha_Kin_pool_1 | 55450 | CAMKIINALPHA | S | 0,86 | 0,87 | 0,45 | 0,51 | 0,87 | 0,48 | 0,01 | 0,04 | 0,55 | 0,55 | 0,62 | 0,06 | 0,14 | -0,43 |
| 45 | Dha_Kin_pool_1 | 5585 | PRKCL1 | S | 0,68 | 0,89 | 0,59 | 0,73 | 0,78 | 0,66 | 0,14 | 0,10 | 0,84 | 0,84 | 0,62 | 0,22 | 0,14 | 1,57 |
| 46 | Dha_Kin_pool_1 | 8767 | RIPK2 | S | 0,65 | 0,67 | 0,38 | 0,36 | 0,66 | 0,37 | 0,01 | 0,01 | 0,56 | 0,56 | 0,62 | 0,05 | 0,14 | -0,37 |
| 47 | Dha_Kin_pool_1 | E | E | D | 0,83 | 0,96 | 0,84 | 1,00 | 0,89 | 0,92 | 0,09 | 0,12 | 1,03 | | 0,62 | | 0,14 | 2,88 |
| 48 | Dha_Kin_pool_1 | E | E | D | 1,00 | 0,95 | 1,08 | 1,06 | 0,98 | 1,07 | 0,04 | 0,02 | 1,09 | | 0,62 | | 0,14 | 3,36 |
| 49 | Dha_Kin_pool_1 | 2872 | MKNK2 | S | 0,93 | 0,96 | 0,49 | 0,59 | 0,95 | 0,54 | 0,02 | 0,07 | 0,57 | 0,57 | 0,62 | 0,05 | 0,14 | -0,36 |
| 50 | Dha_Kin_pool_1 | 51086 | TNNI3K | S | 0,61 | 0,59 | 0,27 | 0,28 | 0,60 | 0,28 | 0,01 | 0,00 | 0,46 | 0,46 | 0,62 | 0,15 | 0,14 | -1,08 |
| 51 | Dha_Kin_pool_1 | 65061 | ALS2CR7 | S | 0,75 | 0,78 | 0,32 | 0,37 | 0,76 | 0,34 | 0,02 | 0,03 | 0,45 | 0,45 | 0,62 | 0,17 | 0,14 | -1,16 |
| 52 | Dha_Kin_pool_1 | 3055 | HCK | S | 0,73 | 0,71 | 0,57 | 0,67 | 0,72 | 0,62 | 0,01 | 0,07 | 0,86 | 0,86 | 0,62 | 0,25 | 0,14 | 1,73 |
| 53 | Dha_Kin_pool_1 | 55589 | BMP2K | S | 1,07 | 1,01 | 0,62 | 0,66 | 1,04 | 0,64 | 0,04 | 0,02 | 0,62 | 0,62 | 0,62 | 0,00 | 0,14 | 0,02 |
| 54 | Dha_Kin_pool_1 | 5257 | PHKB | S | 1,07 | 0,97 | 0,73 | 0,86 | 1,02 | 0,79 | 0,07 | 0,10 | 0,78 | 0,78 | 0,62 | 0,16 | 0,14 | 1,13 |

| | | | | | | | | | | | | | | | | | | | |
|---|---|---|---|---|---|---|---|---|---|---|---|---|---|---|---|---|---|---|---|
| 55 | Dha_Kin_pool_1 | | 7204 | TRIO | S | 0,75 | 1,04 | 0,55 | 0,67 | 0,90 | 0,61 | 0,20 | 0,08 | 0,68 | 0,68 | 0,62 | 0,07 | 0,14 | 0,46 |
| 56 | Dha_Kin_pool_1 | | 55500 | ETNK1 | S | 0,97 | 1,12 | 0,66 | 0,77 | 1,04 | 0,72 | 0,10 | 0,08 | 0,69 | 0,69 | 0,62 | 0,07 | 0,14 | 0,49 |
| 57 | Dha_Kin_pool_1 | | 225689 | ERK8 | S | 0,95 | 0,98 | 0,42 | 0,50 | 0,96 | 0,46 | 0,02 | 0,06 | 0,48 | 0,48 | 0,62 | 0,14 | 0,14 | -0,97 |
| 58 | Dha_Kin_pool_1 | | 83440 | ADP-GK | S | 0,84 | 0,99 | 0,59 | 0,56 | 0,92 | 0,57 | 0,10 | 0,02 | 0,63 | 0,63 | 0,62 | 0,01 | 0,14 | 0,07 |
| 59 | Dha_Kin_pool_1 | | 5294 | PIK3CG | S | 0,96 | 0,91 | 0,78 | 0,79 | 0,93 | 0,79 | 0,03 | 0,01 | 0,84 | 0,84 | 0,62 | 0,22 | 0,14 | 1,58 |
| 60 | Dha_Kin_pool_1 | | 2261 | FGFR3 | S | 0,83 | 0,92 | 0,68 | 0,60 | 0,88 | 0,64 | 0,06 | 0,05 | 0,73 | 0,73 | 0,62 | 0,11 | 0,14 | 0,79 |
| 61 | Dha_Kin_pool_1 | | 51135 | IRAK4 | S | 0,79 | 0,73 | 0,51 | 0,50 | 0,76 | 0,50 | 0,04 | 0,01 | 0,66 | 0,66 | 0,62 | 0,05 | 0,14 | 0,32 |
| 62 | Dha_Kin_pool_1 | | 5129 | PCTK3 | S | 1,01 | 1,05 | 0,67 | 0,72 | 1,03 | 0,69 | 0,03 | 0,03 | 0,67 | 0,67 | 0,62 | 0,06 | 0,14 | 0,40 |
| 63 | Dha_Kin_pool_1 | | 10110 | SGK2 | S | 0,71 | 0,54 | 0,45 | 0,47 | 0,63 | 0,46 | 0,12 | 0,01 | 0,73 | 0,73 | 0,62 | 0,11 | 0,14 | 0,79 |
| 64 | Dha_Kin_pool_1 | | 8798 | DYRK4 | S | 0,74 | 0,79 | 0,55 | 0,60 | 0,76 | 0,57 | 0,03 | 0,04 | 0,75 | 0,75 | 0,62 | 0,14 | 0,14 | 0,95 |
| 65 | Dha_Kin_pool_1 | | 55229 | PANK4 | S | 0,57 | 0,66 | 0,34 | 0,44 | 0,62 | 0,39 | 0,07 | 0,07 | 0,63 | 0,63 | 0,62 | 0,01 | 0,14 | 0,10 |
| 66 | Dha_Kin_pool_1 | | 5568 | PRKACG | S | 0,14 | 0,13 | 0,06 | 0,06 | 0,14 | 0,06 | 0,01 | 0,00 | 0,42 | 0,42 | 0,62 | 0,20 | 0,14 | -1,41 |
| 67 | Dha_Kin_pool_1 | | 2321 | FLT1 | S | 0,78 | 0,79 | 0,50 | 0,58 | 0,78 | 0,54 | 0,01 | 0,05 | 0,69 | 0,69 | 0,62 | 0,08 | 0,14 | 0,53 |
| 68 | Dha_Kin_pool_1 | | 10746 | MAP3K2 | S | 1,12 | 1,00 | 0,53 | 0,59 | 1,06 | 0,56 | 0,08 | 0,04 | 0,53 | 0,53 | 0,62 | 0,08 | 0,14 | -0,58 |
| 69 | Dha_Kin_pool_1 | | 26524 | LATS2 | S | 0,58 | 0,47 | 0,22 | 0,26 | 0,53 | 0,24 | 0,08 | 0,03 | 0,46 | 0,46 | 0,62 | 0,16 | 0,14 | -1,13 |
| 70 | Dha_Kin_pool_1 | | 7272 | TTK | S | 0,66 | 0,63 | 0,27 | 0,27 | 0,65 | 0,27 | 0,02 | 0,00 | 0,42 | 0,42 | 0,62 | 0,20 | 0,14 | -1,38 |
| 71 | Dha_Kin_pool_1 | E | | E | D | 0,92 | 0,93 | 1,10 | 1,05 | 0,93 | 1,08 | 0,00 | 0,03 | 1,16 | | 0,62 | | 0,14 | 3,81 |
| 72 | Dha_Kin_pool_1 | E | | E | D | 1,09 | 1,11 | 1,22 | 0,98 | 1,10 | 1,10 | 0,02 | 0,17 | 1,00 | | 0,62 | | 0,14 | 2,70 |
| 73 | Dha_Kin_pool_1 | | 6850 | SYK | S | 0,39 | 0,31 | 0,06 | 0,08 | 0,35 | 0,07 | 0,06 | 0,02 | 0,19 | 0,19 | 0,62 | 0,42 | 0,14 | -2,97 |
| 74 | Dha_Kin_pool_1 | | 2046 | EPHA8 | S | 0,82 | 0,87 | 0,46 | 0,63 | 0,85 | 0,54 | 0,03 | 0,13 | 0,64 | 0,64 | 0,62 | 0,03 | 0,14 | 0,20 |
| 75 | Dha_Kin_pool_1 | | 1760 | DMPK | S | 0,20 | 0,20 | 0,07 | 0,08 | 0,20 | 0,07 | 0,00 | 0,01 | 0,37 | 0,37 | 0,62 | 0,25 | 0,14 | -1,74 |
| 76 | Dha_Kin_pool_1 | | 140469 | MYO3B | S | 0,85 | 0,98 | 0,61 | 0,65 | 0,92 | 0,63 | 0,09 | 0,03 | 0,69 | 0,69 | 0,62 | 0,07 | 0,14 | 0,50 |
| 77 | Dha_Kin_pool_1 | | 65268 | PRKWNK2 | S | 0,69 | 0,76 | 0,40 | 0,44 | 0,73 | 0,42 | 0,05 | 0,03 | 0,58 | 0,58 | 0,62 | 0,04 | 0,14 | -0,25 |
| 78 | Dha_Kin_pool_1 | | 5747 | PTK2 | S | 0,65 | 0,67 | 0,52 | 0,65 | 0,66 | 0,58 | 0,01 | 0,09 | 0,89 | 0,89 | 0,62 | 0,27 | 0,14 | 1,90 |
| 79 | Dha_Kin_pool_1 | | 2241 | FER | S | 0,59 | 0,95 | 0,63 | 0,58 | 0,77 | 0,60 | 0,26 | 0,03 | 0,78 | 0,78 | 0,62 | 0,17 | 0,14 | 1,16 |
| 80 | Dha_Kin_pool_1 | | 816 | CAMK2B | S | 0,60 | 0,52 | 0,16 | 0,18 | 0,56 | 0,17 | 0,05 | 0,01 | 0,30 | 0,30 | 0,62 | 0,31 | 0,14 | -2,20 |
| 81 | Dha_Kin_pool_1 | | 1608 | DGKG | S | 0,50 | 0,70 | 0,22 | 0,22 | 0,60 | 0,22 | 0,14 | 0,00 | 0,37 | 0,37 | 0,62 | 0,25 | 0,14 | -1,73 |
| 82 | Dha_Kin_pool_1 | | 23031 | MAST3 | S | 0,48 | 0,66 | 0,38 | 0,37 | 0,57 | 0,37 | 0,13 | 0,01 | 0,66 | 0,66 | 0,62 | 0,04 | 0,14 | 0,27 |
| 83 | Dha_Kin_pool_1 | | 10295 | BCKDK | S | 0,76 | 1,16 | 0,76 | 0,74 | 0,96 | 0,75 | 0,28 | 0,01 | 0,78 | 0,78 | 0,62 | 0,17 | 0,14 | 1,17 |
| 84 | Dha_Kin_pool_1 | | 3706 | ITPKA | S | 0,79 | 0,86 | 0,55 | 0,53 | 0,82 | 0,54 | 0,05 | 0,02 | 0,65 | 0,65 | 0,62 | 0,04 | 0,14 | 0,26 |
| 85 | Dha_Kin_pool_1 | | 10783 | NEK6 | S | 0,99 | 1,04 | 0,71 | 0,79 | 1,01 | 0,75 | 0,04 | 0,06 | 0,74 | 0,74 | 0,62 | 0,13 | 0,14 | 0,89 |
| 86 | Dha_Kin_pool_1 | | 640 | BLK | S | 0,91 | 0,89 | 0,54 | 0,65 | 0,90 | 0,60 | 0,02 | 0,08 | 0,66 | 0,66 | 0,62 | 0,05 | 0,14 | 0,32 |
| 87 | Dha_Kin_pool_1 | | 139189 | DGKK | S | 0,87 | 0,82 | 0,49 | 0,56 | 0,84 | 0,52 | 0,03 | 0,05 | 0,62 | 0,62 | 0,62 | 0,01 | 0,14 | 0,04 |
| 88 | Dha_Kin_pool_1 | | 4355 | MPP2 | S | 0,74 | 0,66 | 0,37 | 0,40 | 0,70 | 0,38 | 0,06 | 0,02 | 0,55 | 0,55 | 0,62 | 0,07 | 0,14 | -0,48 |
| 89 | Dha_Kin_pool_1 | | 94 | ACVRL1 | S | 0,73 | 0,56 | 0,37 | 0,43 | 0,64 | 0,40 | 0,12 | 0,04 | 0,62 | 0,62 | 0,62 | 0,00 | 0,14 | 0,00 |
| 90 | Dha_Kin_pool_1 | | 4067 | LYN | S | 0,60 | 0,17 | 0,34 | 0,36 | 0,38 | 0,35 | 0,30 | 0,01 | 0,91 | 0,91 | 0,62 | 0,30 | 0,14 | 2,10 |
| 91 | Dha_Kin_pool_1 | | 5596 | MAPK4 | S | 1,11 | 0,75 | 0,73 | 0,70 | 0,93 | 0,71 | 0,25 | 0,02 | 0,77 | 0,77 | 0,62 | 0,15 | 0,14 | 1,06 |
| 92 | Dha_Kin_pool_1 | | 84930 | MASTL | S | 1,07 | 0,85 | 0,52 | 0,48 | 0,96 | 0,50 | 0,16 | 0,03 | 0,52 | 0,52 | 0,62 | 0,09 | 0,14 | -0,64 |
| 93 | Dha_Kin_pool_1 | | 197259 | FLJ34389 | S | 0,83 | 0,75 | 0,48 | 0,43 | 0,79 | 0,45 | 0,06 | 0,04 | 0,57 | 0,57 | 0,62 | 0,05 | 0,14 | -0,32 |
| 94 | Dha_Kin_pool_1 | | 3098 | HK1 | S | 0,54 | 0,72 | 0,27 | 0,25 | 0,63 | 0,26 | 0,13 | 0,01 | 0,41 | 0,41 | 0,62 | 0,21 | 0,14 | -1,45 |
| 95 | Dha_Kin_pool_1 | E | | E | D | 0,96 | 0,84 | 0,90 | 1,04 | 0,90 | 0,97 | 0,08 | 0,10 | 1,07 | | 0,62 | | 0,14 | 3,22 |
| 96 | Dha_Kin_pool_1 | E | | E | D | 1,06 | 1,04 | 0,77 | 0,80 | 1,05 | 0,78 | 0,02 | 0,02 | 0,75 | | 0,62 | | 0,14 | 0,93 |
| 97 | Dha_Kin_pool_1 | | 699 | BUB1 | S | 0,13 | 0,14 | 0,07 | 0,08 | 0,14 | 0,07 | 0,00 | 0,01 | 0,53 | 0,53 | 0,62 | 0,08 | 0,14 | -0,57 |
| 98 | Dha_Kin_pool_1 | | 127933 | UHMK1 | S | 0,99 | 1,07 | 0,80 | 0,94 | 1,03 | 0,87 | 0,06 | 0,09 | 0,85 | 0,85 | 0,62 | 0,23 | 0,14 | 1,62 |
| 99 | Dha_Kin_pool_1 | | 5587 | PRKCM | S | 0,64 | 0,85 | 0,65 | 0,40 | 0,74 | 0,52 | 0,14 | 0,18 | 0,70 | 0,70 | 0,62 | 0,09 | 0,14 | 0,62 |
| 100 | Dha_Kin_pool_1 | | 5298 | PIK4CB | S | 0,51 | 0,64 | 0,20 | 0,21 | 0,57 | 0,21 | 0,09 | 0,01 | 0,36 | 0,36 | 0,62 | 0,25 | 0,14 | -1,77 |
| 101 | Dha_Kin_pool_1 | | 695 | BTK | S | 0,90 | 0,85 | 0,48 | 0,45 | 0,88 | 0,47 | 0,03 | 0,02 | 0,53 | 0,53 | 0,62 | 0,08 | 0,14 | -0,60 |
| 102 | Dha_Kin_pool_1 | | 2931 | GSK3A | S | 0,83 | 0,97 | 0,72 | 0,92 | 0,90 | 0,82 | 0,10 | 0,14 | 0,91 | 0,91 | 0,62 | 0,29 | 0,14 | 2,06 |
| 103 | Dha_Kin_pool_1 | | 1454 | CSNK1E | S | 0,74 | 0,74 | 0,49 | 0,48 | 0,74 | 0,49 | 0,00 | 0,01 | 0,66 | 0,66 | 0,62 | 0,04 | 0,14 | 0,29 |
| 104 | Dha_Kin_pool_1 | | 23729 | CARKL | S | 0,73 | 0,65 | 0,50 | 0,30 | 0,69 | 0,40 | 0,06 | 0,15 | 0,58 | 0,58 | 0,62 | 0,03 | 0,14 | -0,24 |
| 105 | Dha_Kin_pool_1 | | 132 | ADK | S | 0,62 | 0,91 | 0,59 | 0,74 | 0,76 | 0,66 | 0,20 | 0,10 | 0,87 | 0,87 | 0,62 | 0,25 | 0,14 | 1,77 |
| 106 | Dha_Kin_pool_1 | | 27 | ABL2 | S | 0,82 | 0,86 | 0,39 | 0,35 | 0,84 | 0,37 | 0,03 | 0,02 | 0,44 | 0,44 | 0,62 | 0,18 | 0,14 | -1,24 |
| 107 | Dha_Kin_pool_1 | | 5347 | PLK1 | S | 0,08 | 0,09 | 0,05 | 0,06 | 0,09 | 0,05 | 0,01 | 0,01 | 0,60 | 0,60 | 0,62 | 0,01 | 0,14 | -0,10 |
| 108 | Dha_Kin_pool_1 | | 50488 | MINK | S | 0,83 | 0,72 | 0,71 | 0,63 | 0,78 | 0,67 | 0,08 | 0,06 | 0,86 | 0,86 | 0,62 | 0,24 | 0,14 | 1,71 |
| 109 | Dha_Kin_pool_1 | | 51265 | CDKL3 | S | 0,78 | 0,93 | 0,45 | 0,46 | 0,85 | 0,45 | 0,11 | 0,01 | 0,53 | 0,53 | 0,62 | 0,08 | 0,14 | -0,58 |

| | | | | | | | | | | | | | | | | | | | |
|---|---|---|---|---|---|---|---|---|---|---|---|---|---|---|---|---|---|---|---|
| 110 | Dha_Kin_pool_1 | | 9020 | MAP3K14 | S | 0,75 | 0,73 | 0,41 | 0,51 | 0,74 | 0,46 | 0,01 | 0,07 | 0,62 | 0,62 | 0,62 | 0,01 | 0,14 | 0,04 |
| 111 | Dha_Kin_pool_1 | | 6885 | MAP3K7 | S | 0,81 | 0,73 | 0,53 | 0,60 | 0,77 | 0,56 | 0,06 | 0,05 | 0,73 | 0,73 | 0,62 | 0,12 | 0,14 | 0,81 |
| 112 | Dha_Kin_pool_1 | | 1152 | CKB | S | 0,84 | 0,94 | 0,67 | 0,87 | 0,89 | 0,77 | 0,07 | 0,14 | 0,86 | 0,86 | 0,62 | 0,25 | 0,14 | 1,75 |
| 113 | Dha_Kin_pool_1 | | 23552 | CCRK | S | 0,61 | 0,71 | 0,49 | 0,57 | 0,66 | 0,53 | 0,08 | 0,05 | 0,80 | 0,80 | 0,62 | 0,19 | 0,14 | 1,32 |
| 114 | Dha_Kin_pool_1 | | 5601 | MAPK9 | S | 0,85 | 0,90 | 0,58 | 0,62 | 0,87 | 0,60 | 0,04 | 0,03 | 0,68 | 0,68 | 0,62 | 0,07 | 0,14 | 0,48 |
| 115 | Dha_Kin_pool_1 | | 10188 | TNK2 | S | 0,86 | 0,82 | 0,55 | 0,62 | 0,84 | 0,58 | 0,03 | 0,04 | 0,69 | 0,69 | 0,62 | 0,08 | 0,14 | 0,54 |
| 116 | Dha_Kin_pool_1 | | 149420 | PDIK1L | S | 1,08 | 1,20 | 0,74 | 0,74 | 1,14 | 0,74 | 0,09 | 0,00 | 0,65 | 0,65 | 0,62 | 0,03 | 0,14 | 0,21 |
| 117 | Dha_Kin_pool_1 | | 558 | AXL | S | 0,83 | 0,89 | 0,33 | 0,38 | 0,86 | 0,36 | 0,04 | 0,03 | 0,41 | 0,41 | 0,62 | 0,20 | 0,14 | -1,43 |
| 118 | Dha_Kin_pool_1 | | 54101 | ANKRD3 | S | 1,06 | 0,79 | 0,46 | 0,47 | 0,93 | 0,46 | 0,19 | 0,01 | 0,50 | 0,50 | 0,62 | 0,12 | 0,14 | -0,81 |
| 119 | Dha_Kin_pool_1 | E | | E | D | 1,04 | 1,21 | 1,09 | 0,82 | 1,13 | 0,96 | 0,13 | 0,19 | 0,85 | | 0,62 | | 0,14 | 1,65 |
| 120 | Dha_Kin_pool_1 | E | | E | D | 1,15 | 1,03 | 1,07 | 1,08 | 1,09 | 1,08 | 0,09 | 0,01 | 0,99 | | 0,62 | | 0,14 | 2,60 |
| 121 | Dha_Kin_pool_1 | | 8737 | RIPK1 | S | 1,04 | 1,02 | 0,73 | 0,88 | 1,03 | 0,80 | 0,02 | 0,10 | 0,78 | 0,78 | 0,62 | 0,16 | 0,14 | 1,14 |
| 122 | Dha_Kin_pool_1 | | 9578 | CDC42BPB | S | 1,01 | 1,06 | 0,81 | 0,83 | 1,04 | 0,82 | 0,03 | 0,01 | 0,79 | 0,79 | 0,62 | 0,18 | 0,14 | 1,23 |
| 123 | Dha_Kin_pool_1 | | 3718 | JAK3 | S | 0,62 | 0,78 | 0,35 | 0,41 | 0,70 | 0,38 | 0,11 | 0,04 | 0,54 | 0,54 | 0,62 | 0,07 | 0,14 | -0,50 |
| 124 | Dha_Kin_pool_1 | | 83942 | STK22D | S | 0,88 | 0,78 | 0,44 | 0,51 | 0,83 | 0,48 | 0,07 | 0,05 | 0,57 | 0,57 | 0,62 | 0,04 | 0,14 | -0,31 |
| 125 | Dha_Kin_pool_1 | | 983 | CDC2 | S | 0,53 | 0,63 | 0,21 | 0,26 | 0,58 | 0,24 | 0,07 | 0,03 | 0,40 | 0,40 | 0,62 | 0,21 | 0,14 | -1,49 |
| 126 | Dha_Kin_pool_1 | | 11040 | PIM2 | S | 0,56 | 0,62 | 0,18 | 0,25 | 0,59 | 0,21 | 0,04 | 0,05 | 0,36 | 0,36 | 0,62 | 0,25 | 0,14 | -1,78 |
| 127 | Dha_Kin_pool_1 | | 120892 | LRRK2 | S | 1,10 | 1,16 | 0,45 | 0,60 | 1,13 | 0,52 | 0,05 | 0,11 | 0,46 | 0,46 | 0,62 | 0,15 | 0,14 | -1,07 |
| 128 | Dha_Kin_pool_1 | | 3099 | HK2 | S | 0,48 | 0,49 | 0,19 | 0,23 | 0,48 | 0,21 | 0,01 | 0,03 | 0,44 | 0,44 | 0,62 | 0,18 | 0,14 | -1,27 |
| 129 | Dha_Kin_pool_1 | | 64122 | FN3K | S | 0,68 | 0,91 | 0,45 | 0,59 | 0,79 | 0,52 | 0,16 | 0,09 | 0,65 | 0,65 | 0,62 | 0,04 | 0,14 | 0,26 |
| 130 | Dha_Kin_pool_1 | | 5297 | PIK4CA | S | 0,41 | 0,44 | 0,26 | 0,22 | 0,43 | 0,24 | 0,02 | 0,02 | 0,56 | 0,56 | 0,62 | 0,05 | 0,14 | -0,37 |
| 131 | Dha_Kin_pool_1 | | 83983 | SSTK | S | 0,69 | 0,81 | 0,43 | 0,54 | 0,75 | 0,48 | 0,08 | 0,07 | 0,64 | 0,64 | 0,62 | 0,03 | 0,14 | 0,20 |
| 132 | Dha_Kin_pool_1 | | 9344 | TAO1 | S | 0,69 | 0,81 | 0,54 | 0,53 | 0,75 | 0,54 | 0,08 | 0,01 | 0,72 | 0,72 | 0,62 | 0,10 | 0,14 | 0,70 |
| 133 | Dha_Kin_pool_1 | | 4832 | NME3 | S | 0,60 | 0,67 | 0,43 | 0,44 | 0,64 | 0,44 | 0,05 | 0,01 | 0,69 | 0,69 | 0,62 | 0,07 | 0,14 | 0,49 |
| 134 | Dha_Kin_pool_1 | | 2260 | FGFR1 | S | 0,37 | 0,41 | 0,23 | 0,19 | 0,39 | 0,21 | 0,03 | 0,03 | 0,52 | 0,52 | 0,62 | 0,09 | 0,14 | -0,65 |
| 135 | Dha_Kin_pool_1 | | 11284 | PNKP | S | 0,93 | 0,68 | 0,56 | 0,55 | 0,80 | 0,55 | 0,17 | 0,01 | 0,69 | 0,69 | 0,62 | 0,07 | 0,14 | 0,51 |
| 136 | Dha_Kin_pool_1 | | 2064 | ERBB2 | S | 0,99 | 0,89 | 0,54 | 0,67 | 0,94 | 0,61 | 0,07 | 0,09 | 0,64 | 0,64 | 0,62 | 0,03 | 0,14 | 0,20 |
| 137 | Dha_Kin_pool_1 | | 55224 | FLJ10761 | S | 0,80 | 0,95 | 0,60 | 0,66 | 0,88 | 0,63 | 0,11 | 0,05 | 0,72 | 0,72 | 0,62 | 0,10 | 0,14 | 0,72 |
| 138 | Dha_Kin_pool_1 | | 91754 | NEK9 | S | 0,75 | 0,88 | 0,53 | 0,56 | 0,81 | 0,55 | 0,10 | 0,02 | 0,67 | 0,67 | 0,62 | 0,06 | 0,14 | 0,40 |
| 139 | Dha_Kin_pool_1 | | 84197 | FLJ23356 | S | 0,87 | 1,00 | 0,61 | 0,62 | 0,94 | 0,62 | 0,09 | 0,01 | 0,66 | 0,66 | 0,62 | 0,04 | 0,14 | 0,30 |
| 140 | Dha_Kin_pool_1 | | 2322 | FLT3 | S | 0,91 | 0,85 | 0,30 | 0,28 | 0,88 | 0,29 | 0,04 | 0,01 | 0,33 | 0,33 | 0,62 | 0,29 | 0,14 | -2,03 |
| 141 | Dha_Kin_pool_1 | | 5305 | PIP5K2A | S | 1,27 | 0,96 | 0,58 | 0,66 | 1,11 | 0,62 | 0,22 | 0,06 | 0,56 | 0,56 | 0,62 | 0,06 | 0,14 | -0,40 |
| 142 | Dha_Kin_pool_1 | | 2869 | GRK5 | S | 0,63 | 0,77 | 0,50 | 0,56 | 0,70 | 0,53 | 0,10 | 0,05 | 0,76 | 0,76 | 0,62 | 0,14 | 0,14 | 0,99 |
| 143 | Dha_Kin_pool_1 | E | | E | D | 1,00 | 1,00 | 1,07 | 1,11 | 1,00 | 1,09 | 0,00 | 0,03 | 1,09 | | 0,62 | | 0,14 | 3,33 |
| 144 | Dha_Kin_pool_1 | E | | E | D | 0,96 | 0,96 | 1,00 | 0,94 | 0,96 | 0,97 | 0,01 | 0,04 | 1,01 | | 0,62 | | 0,14 | 2,77 |
| 145 | Dha_Kin_pool_1 | | 3984 | LIMK1 | S | 1,19 | 1,08 | 0,73 | 0,66 | 1,14 | 0,69 | 0,08 | 0,04 | 0,61 | 0,61 | 0,62 | 0,01 | 0,14 | -0,04 |
| 146 | Dha_Kin_pool_1 | | 7371 | UMPK | S | 1,11 | 1,26 | 0,75 | 0,83 | 1,18 | 0,79 | 0,10 | 0,05 | 0,67 | 0,67 | 0,62 | 0,05 | 0,14 | 0,38 |
| 147 | Dha_Kin_pool_1 | | 8844 | KSR | S | 0,36 | 0,42 | 0,10 | 0,12 | 0,39 | 0,11 | 0,04 | 0,02 | 0,28 | 0,28 | 0,62 | 0,34 | 0,14 | -2,36 |
| 148 | Dha_Kin_pool_1 | | 9448 | MAP4K4 | S | 0,82 | 0,86 | 0,53 | 0,59 | 0,84 | 0,56 | 0,03 | 0,04 | 0,67 | 0,67 | 0,62 | 0,05 | 0,14 | 0,36 |
| 149 | Dha_Kin_pool_1 | | 23683 | PRKCN | S | 1,03 | 1,02 | 0,53 | 0,61 | 1,02 | 0,57 | 0,01 | 0,06 | 0,56 | 0,56 | 0,62 | 0,06 | 0,14 | -0,41 |
| 150 | Dha_Kin_pool_1 | | 84630 | TTBK1 | S | 0,84 | 0,91 | 0,50 | 0,55 | 0,87 | 0,52 | 0,05 | 0,03 | 0,60 | 0,60 | 0,62 | 0,02 | 0,14 | -0,13 |
| 151 | Dha_Kin_pool_1 | | 6093 | ROCK1 | S | 0,68 | 0,74 | 0,44 | 0,35 | 0,71 | 0,39 | 0,04 | 0,06 | 0,56 | 0,56 | 0,62 | 0,06 | 0,14 | -0,42 |
| 152 | Dha_Kin_pool_1 | | 1612 | DAPK1 | S | 0,75 | 0,88 | 0,43 | 0,55 | 0,81 | 0,49 | 0,09 | 0,09 | 0,61 | 0,61 | 0,62 | 0,01 | 0,14 | -0,06 |
| 153 | Dha_Kin_pool_1 | | 10922 | FASTK | S | 0,96 | 1,00 | 0,69 | 0,71 | 0,98 | 0,70 | 0,03 | 0,01 | 0,71 | 0,71 | 0,62 | 0,10 | 0,14 | 0,68 |
| 154 | Dha_Kin_pool_1 | | 5593 | PRKG2 | S | 0,61 | 0,91 | 0,58 | 0,67 | 0,76 | 0,62 | 0,22 | 0,07 | 0,82 | 0,82 | 0,62 | 0,21 | 0,14 | 1,44 |
| 155 | Dha_Kin_pool_1 | | 4354 | MPP1 | S | 0,56 | 1,13 | 0,69 | 0,76 | 0,84 | 0,73 | 0,40 | 0,05 | 0,86 | 0,86 | 0,62 | 0,24 | 0,14 | 1,71 |
| 156 | Dha_Kin_pool_1 | | 1436 | CSF1R | S | 0,48 | 0,49 | 0,28 | 0,26 | 0,48 | 0,27 | 0,01 | 0,01 | 0,56 | 0,56 | 0,62 | 0,06 | 0,14 | -0,42 |
| 157 | Dha_Kin_pool_1 | | 8445 | DYRK2 | S | 0,71 | 0,69 | 0,32 | 0,29 | 0,70 | 0,31 | 0,01 | 0,02 | 0,44 | 0,44 | 0,62 | 0,18 | 0,14 | -1,25 |
| 158 | Dha_Kin_pool_1 | | 5586 | PRKCL2 | S | 0,58 | 0,54 | 0,31 | 0,30 | 0,56 | 0,31 | 0,03 | 0,00 | 0,55 | 0,55 | 0,62 | 0,07 | 0,14 | -0,48 |
| 159 | Dha_Kin_pool_1 | | 6197 | RPS6KA3 | S | 0,68 | 0,70 | 0,47 | 0,52 | 0,69 | 0,49 | 0,02 | 0,03 | 0,71 | 0,71 | 0,62 | 0,09 | 0,14 | 0,67 |
| 160 | Dha_Kin_pool_1 | | 7010 | TEK | S | 0,74 | 0,78 | 0,58 | 0,70 | 0,76 | 0,64 | 0,02 | 0,08 | 0,84 | 0,84 | 0,62 | 0,23 | 0,14 | 1,59 |
| 161 | Dha_Kin_pool_1 | | 53354 | PANK1 | S | 0,85 | 0,85 | 0,47 | 0,51 | 0,85 | 0,49 | 0,00 | 0,03 | 0,58 | 0,58 | 0,62 | 0,04 | 0,14 | -0,28 |
| 162 | Dha_Kin_pool_1 | | 1456 | CSNK1G3 | S | 0,89 | 1,03 | 0,78 | 0,57 | 0,96 | 0,68 | 0,10 | 0,15 | 0,70 | 0,70 | 0,62 | 0,09 | 0,14 | 0,61 |
| 163 | Dha_Kin_pool_1 | | 5261 | PHKG2 | S | 0,30 | 0,40 | 0,29 | 0,34 | 0,35 | 0,32 | 0,08 | 0,03 | 0,91 | 0,91 | 0,62 | 0,29 | 0,14 | 2,04 |
| 164 | Dha_Kin_pool_1 | | 5598 | MAPK7 | S | 0,84 | 0,83 | 0,53 | 0,65 | 0,83 | 0,59 | 0,01 | 0,09 | 0,71 | 0,71 | 0,62 | 0,09 | 0,14 | 0,66 |

| | | | | | | | | | | | | | | | | | | | |
|---|---|---|---|---|---|---|---|---|---|---|---|---|---|---|---|---|---|---|---|
| 165 | Dha_Kin_pool_1 | | 817 | CAMK2D | S | 1,00 | 0,98 | 0,42 | 0,60 | 0,99 | 0,51 | 0,02 | 0,12 | 0,51 | 0,51 | 0,62 | 0,10 | 0,14 | -0,71 |
| 166 | Dha_Kin_pool_1 | | 11183 | MAP4K5 | S | 0,90 | 0,89 | 0,78 | 0,96 | 0,89 | 0,87 | 0,01 | 0,13 | 0,97 | 0,97 | 0,62 | 0,36 | 0,14 | 2,52 |
| 167 | Dha_Kin_pool_1 | E | | E | D | 1,09 | 1,03 | 0,88 | 0,93 | 1,06 | 0,90 | 0,05 | 0,04 | 0,85 | | 0,62 | | 0,14 | 1,67 |
| 168 | Dha_Kin_pool_1 | E | | E | D | 1,06 | 1,02 | 0,85 | 0,87 | 1,04 | 0,86 | 0,03 | 0,01 | 0,83 | | 0,62 | | 0,14 | 1,49 |
| 169 | Dha_Kin_pool_1 | | 29941 | PKN3 | S | 0,89 | 0,92 | 0,72 | 0,69 | 0,90 | 0,70 | 0,02 | 0,02 | 0,78 | 0,78 | 0,62 | 0,16 | 0,14 | 1,15 |
| 170 | Dha_Kin_pool_1 | | 57143 | ADCK1 | S | 0,91 | 0,70 | 0,48 | 0,63 | 0,80 | 0,56 | 0,15 | 0,10 | 0,69 | 0,69 | 0,62 | 0,08 | 0,14 | 0,53 |
| 171 | Dha_Kin_pool_1 | | 1326 | MAP3K8 | S | 0,77 | 0,94 | 0,42 | 0,48 | 0,85 | 0,45 | 0,12 | 0,04 | 0,53 | 0,53 | 0,62 | 0,09 | 0,14 | -0,61 |
| 172 | Dha_Kin_pool_1 | | 1024 | CDK8 | S | 0,88 | 0,76 | 0,42 | 0,47 | 0,82 | 0,44 | 0,08 | 0,03 | 0,54 | 0,54 | 0,62 | 0,07 | 0,14 | -0,52 |
| 173 | Dha_Kin_pool_1 | | 8569 | MKNK1 | S | 0,94 | 0,74 | 0,56 | 0,57 | 0,84 | 0,56 | 0,14 | 0,01 | 0,67 | 0,67 | 0,62 | 0,05 | 0,14 | 0,35 |
| 174 | Dha_Kin_pool_1 | | 5288 | PIK3C2G | S | 0,76 | 0,72 | 0,47 | 0,50 | 0,74 | 0,48 | 0,03 | 0,02 | 0,66 | 0,66 | 0,62 | 0,04 | 0,14 | 0,28 |
| 175 | Dha_Kin_pool_1 | | 1019 | CDK4 | S | 0,73 | 0,72 | 0,22 | 0,32 | 0,73 | 0,27 | 0,01 | 0,07 | 0,37 | 0,37 | 0,62 | 0,24 | 0,14 | -1,70 |
| 176 | Dha_Kin_pool_1 | | 5292 | PIM1 | S | 0,94 | 0,99 | 0,45 | 0,45 | 0,97 | 0,45 | 0,03 | 0,00 | 0,47 | 0,47 | 0,62 | 0,15 | 0,14 | -1,05 |
| 177 | Dha_Kin_pool_1 | | 1609 | DGKQ | S | 0,83 | 0,79 | 0,41 | 0,42 | 0,81 | 0,42 | 0,03 | 0,00 | 0,51 | 0,51 | 0,62 | 0,10 | 0,14 | -0,71 |
| 178 | Dha_Kin_pool_1 | | 7786 | MAP3K12 | S | 0,41 | 0,35 | 0,06 | 0,07 | 0,38 | 0,07 | 0,04 | 0,01 | 0,17 | 0,17 | 0,62 | 0,44 | 0,14 | -3,12 |
| 179 | Dha_Kin_pool_1 | | 81629 | STK22C | S | 0,72 | 0,95 | 0,42 | 0,46 | 0,84 | 0,44 | 0,16 | 0,03 | 0,52 | 0,52 | 0,62 | 0,09 | 0,14 | -0,65 |
| 180 | Dha_Kin_pool_1 | | 472 | ATM | S | 0,86 | 1,09 | 0,58 | 0,60 | 0,97 | 0,59 | 0,16 | 0,02 | 0,61 | 0,61 | 0,62 | 0,01 | 0,14 | -0,05 |
| 181 | Dha_Kin_pool_1 | | 5256 | PHKA2 | S | 0,89 | 0,79 | 0,64 | 0,69 | 0,84 | 0,66 | 0,07 | 0,03 | 0,79 | 0,79 | 0,62 | 0,18 | 0,14 | 1,23 |
| 182 | Dha_Kin_pool_1 | | 55750 | MULK | S | 0,55 | 0,51 | 0,22 | 0,20 | 0,53 | 0,21 | 0,03 | 0,01 | 0,40 | 0,40 | 0,62 | 0,22 | 0,14 | -1,54 |
| 183 | Dha_Kin_pool_1 | | 2870 | GRK6 | S | 0,74 | 0,73 | 0,40 | 0,47 | 0,73 | 0,43 | 0,01 | 0,05 | 0,59 | 0,59 | 0,62 | 0,03 | 0,14 | -0,18 |
| 184 | Dha_Kin_pool_1 | | 658 | BMPR1B | S | 0,87 | 1,03 | 0,57 | 0,63 | 0,95 | 0,60 | 0,12 | 0,04 | 0,63 | 0,63 | 0,62 | 0,01 | 0,14 | 0,10 |
| 185 | Dha_Kin_pool_1 | | 65975 | STK33 | S | 0,98 | 1,03 | 0,71 | 0,76 | 1,00 | 0,74 | 0,04 | 0,03 | 0,73 | 0,73 | 0,62 | 0,12 | 0,14 | 0,83 |
| 186 | Dha_Kin_pool_1 | | 57538 | MIDORI | S | 0,24 | 0,27 | 0,16 | 0,16 | 0,26 | 0,16 | 0,02 | 0,00 | 0,64 | 0,64 | 0,62 | 0,02 | 0,14 | 0,14 |
| 187 | Dha_Kin_pool_1 | | 8576 | STK16 | S | 0,68 | 0,64 | 0,43 | 0,49 | 0,66 | 0,46 | 0,03 | 0,04 | 0,70 | 0,70 | 0,62 | 0,08 | 0,14 | 0,58 |
| 188 | Dha_Kin_pool_1 | | 55361 | PI4KII | S | 0,82 | 1,03 | 0,54 | 0,64 | 0,92 | 0,59 | 0,15 | 0,07 | 0,63 | 0,63 | 0,62 | 0,02 | 0,14 | 0,13 |
| 189 | Dha_Kin_pool_1 | | 57172 | CAMK1G | S | 0,96 | 1,07 | 0,68 | 0,83 | 1,02 | 0,75 | 0,08 | 0,11 | 0,74 | 0,74 | 0,62 | 0,13 | 0,14 | 0,88 |
| 190 | Dha_Kin_pool_1 | | 55561 | HSMDPKIN | S | 0,73 | 0,92 | 0,88 | 0,68 | 0,83 | 0,78 | 0,13 | 0,14 | 0,95 | 0,95 | 0,62 | 0,33 | 0,14 | 2,34 |
| 191 | Dha_Kin_pool_1 | E | | E | D | 0,86 | 0,97 | 1,02 | 1,20 | 0,92 | 1,11 | 0,08 | 0,13 | 1,22 | | 0,62 | | 0,14 | 4,21 |
| 192 | Dha_Kin_pool_1 | E | | E | D | 0,98 | 1,10 | 0,85 | 0,74 | 1,04 | 0,80 | 0,09 | 0,08 | 0,77 | | 0,62 | | 0,14 | 1,05 |
| 193 | Dha_Kin_pool_1 | | 6793 | STK10 | S | 0,56 | 0,57 | 0,24 | 0,39 | 0,57 | 0,31 | 0,01 | 0,11 | 0,55 | 0,55 | 0,62 | 0,07 | 0,14 | -0,46 |
| 194 | Dha_Kin_pool_1 | | 9863 | AIP1 | S | 0,46 | 0,48 | 0,15 | 0,24 | 0,47 | 0,19 | 0,02 | 0,07 | 0,41 | 0,41 | 0,62 | 0,21 | 0,14 | -1,45 |
| 195 | Dha_Kin_pool_1 | | 25 | ABL1 | S | 0,64 | 0,95 | 0,38 | 0,47 | 0,80 | 0,42 | 0,22 | 0,07 | 0,53 | 0,53 | 0,62 | 0,09 | 0,14 | -0,60 |
| 196 | Dha_Kin_pool_1 | | 56924 | PAK6 | S | 0,78 | 1,16 | 0,40 | 0,41 | 0,97 | 0,41 | 0,27 | 0,01 | 0,42 | 0,42 | 0,62 | 0,20 | 0,14 | -1,38 |
| 197 | Dha_Kin_pool_1 | | 79672 | FN3KRP | S | 0,66 | 0,76 | 0,35 | 0,28 | 0,71 | 0,32 | 0,07 | 0,05 | 0,45 | 0,45 | 0,62 | 0,17 | 0,14 | -1,20 |
| 198 | Dha_Kin_pool_1 | | 208 | AKT2 | S | 0,68 | 0,63 | 0,34 | 0,45 | 0,65 | 0,39 | 0,04 | 0,08 | 0,60 | 0,60 | 0,62 | 0,01 | 0,14 | -0,10 |
| 199 | Dha_Kin_pool_1 | | 138429 | PIP5KL1 | S | 0,57 | 0,79 | 0,56 | 0,65 | 0,68 | 0,60 | 0,15 | 0,06 | 0,89 | 0,89 | 0,62 | 0,28 | 0,14 | 1,95 |
| 200 | Dha_Kin_pool_1 | | 341676 | NEK5 | S | 0,83 | 0,91 | 0,43 | 0,46 | 0,87 | 0,45 | 0,06 | 0,02 | 0,51 | 0,51 | 0,62 | 0,10 | 0,14 | -0,71 |
| 201 | Dha_Kin_pool_1 | | 10769 | PLK2 | S | 0,87 | 0,67 | 0,44 | 0,46 | 0,77 | 0,45 | 0,14 | 0,02 | 0,58 | 0,58 | 0,62 | 0,03 | 0,14 | -0,24 |
| 202 | Dha_Kin_pool_1 | | 1455 | CSNK1G2 | S | 1,19 | 1,27 | 0,74 | 0,82 | 1,23 | 0,78 | 0,06 | 0,06 | 0,63 | 0,63 | 0,62 | 0,02 | 0,14 | 0,12 |
| 203 | Dha_Kin_pool_1 | | 2011 | MARK2 | S | 0,84 | 0,84 | 0,48 | 0,48 | 0,84 | 0,48 | 0,01 | 0,00 | 0,57 | 0,57 | 0,62 | 0,04 | 0,14 | -0,29 |
| 204 | Dha_Kin_pool_1 | | 23043 | TNIK | S | 1,02 | 1,10 | 0,84 | 0,90 | 1,06 | 0,87 | 0,06 | 0,04 | 0,82 | 0,82 | 0,62 | 0,20 | 0,14 | 1,43 |
| 205 | Dha_Kin_pool_1 | | 5315 | PKM2 | S | 0,83 | 0,80 | 0,43 | 0,39 | 0,81 | 0,41 | 0,02 | 0,03 | 0,51 | 0,51 | 0,62 | 0,11 | 0,14 | -0,76 |
| 206 | Dha_Kin_pool_1 | | 6792 | CDKL5 | S | 1,01 | 1,04 | 0,73 | 0,60 | 1,02 | 0,67 | 0,03 | 0,09 | 0,65 | 0,65 | 0,62 | 0,04 | 0,14 | 0,25 |
| 207 | Dha_Kin_pool_1 | | 5290 | PIK3CA | S | 0,13 | 0,13 | 0,08 | 0,08 | 0,13 | 0,08 | 0,01 | 0,00 | 0,61 | 0,61 | 0,62 | 0,00 | 0,14 | -0,01 |
| 208 | Dha_Kin_pool_1 | | 23387 | KIAA0999 | S | 0,93 | 1,15 | 0,64 | 0,82 | 1,04 | 0,73 | 0,16 | 0,13 | 0,70 | 0,70 | 0,62 | 0,08 | 0,14 | 0,58 |
| 209 | Dha_Kin_pool_1 | | 10733 | PLK4 | S | 0,26 | 0,38 | 0,18 | 0,26 | 0,32 | 0,22 | 0,09 | 0,06 | 0,69 | 0,69 | 0,62 | 0,07 | 0,14 | 0,51 |
| 210 | Dha_Kin_pool_1 | | 10114 | HIPK3 | S | 0,92 | 0,84 | 0,54 | 0,36 | 0,88 | 0,45 | 0,05 | 0,13 | 0,52 | 0,52 | 0,62 | 0,10 | 0,14 | -0,70 |
| 211 | Dha_Kin_pool_1 | | 7301 | TYRO3 | S | 0,49 | 0,46 | 0,11 | 0,17 | 0,47 | 0,14 | 0,02 | 0,04 | 0,30 | 0,30 | 0,62 | 0,32 | 0,14 | -2,22 |
| 212 | Dha_Kin_pool_1 | | 5595 | MAPK3 | S | 0,73 | 0,79 | 0,17 | 0,15 | 0,76 | 0,16 | 0,05 | 0,01 | 0,21 | 0,21 | 0,62 | 0,41 | 0,14 | -2,85 |
| 213 | Dha_Kin_pool_1 | | 7525 | YES1 | S | 0,88 | 0,81 | 0,35 | 0,37 | 0,84 | 0,36 | 0,05 | 0,01 | 0,43 | 0,43 | 0,62 | 0,19 | 0,14 | -1,33 |
| 214 | Dha_Kin_pool_1 | | 81602 | CDADC1 | S | 1,08 | 0,97 | 0,37 | 0,52 | 1,02 | 0,45 | 0,08 | 0,11 | 0,44 | 0,44 | 0,62 | 0,18 | 0,14 | -1,25 |
| 215 | Dha_Kin_pool_1 | E | | E | A | 0,65 | 0,75 | 0,77 | 0,85 | 0,70 | 0,81 | 0,07 | 0,06 | 1,16 | | 0,62 | | 0,14 | 3,85 |
| 216 | Dha_Kin_pool_1 | E | | E | A | 0,71 | 0,67 | 0,63 | 0,76 | 0,69 | 0,69 | 0,03 | 0,09 | 1,01 | | 0,62 | | 0,14 | 2,74 |
| 217 | Dha_Kin_pool_1 | | 56164 | STK31 | S | 1,16 | 1,21 | 0,61 | 0,75 | 1,18 | 0,68 | 0,03 | 0,09 | 0,58 | 0,58 | 0,62 | 0,04 | 0,14 | -0,28 |
| 218 | Dha_Kin_pool_1 | | 10000 | AKT3 | S | 1,03 | 0,89 | 0,47 | 0,61 | 0,96 | 0,54 | 0,10 | 0,10 | 0,56 | 0,56 | 0,62 | 0,05 | 0,14 | -0,38 |
| 219 | Dha_Kin_pool_1 | | 84451 | KIAA1804 | S | 0,76 | 0,85 | 0,50 | 0,51 | 0,81 | 0,50 | 0,06 | 0,00 | 0,63 | 0,63 | 0,62 | 0,01 | 0,14 | 0,07 |

| | | | | | | | | | | | | | | | | | | | |
|---|---|---|---|---|---|---|---|---|---|---|---|---|---|---|---|---|---|---|---|
| 220 | Dha_Kin_pool_1 | | 55577 | NAGK | S | 0,70 | 0,91 | 0,48 | 0,54 | 0,80 | 0,51 | 0,15 | 0,04 | 0,64 | 0,64 | 0,62 | 0,02 | 0,14 | 0,15 |
| 221 | Dha_Kin_pool_1 | | 3654 | IRAK1 | S | 0,63 | 0,48 | 0,31 | 0,37 | 0,55 | 0,34 | 0,11 | 0,05 | 0,61 | 0,61 | 0,62 | 0,00 | 0,14 | -0,02 |
| 222 | Dha_Kin_pool_1 | | 29904 | EEF2K | S | 0,92 | 0,66 | 0,29 | 0,26 | 0,79 | 0,27 | 0,19 | 0,02 | 0,34 | 0,34 | 0,62 | 0,27 | 0,14 | -1,91 |
| 223 | Dha_Kin_pool_1 | | 255239 | ANKK1 | S | 0,55 | 0,63 | 0,25 | 0,29 | 0,59 | 0,27 | 0,05 | 0,03 | 0,46 | 0,46 | 0,62 | 0,16 | 0,14 | -1,09 |
| 224 | Dha_Kin_pool_1 | | 4217 | MAP3K5 | S | 0,81 | 0,84 | 0,51 | 0,61 | 0,82 | 0,56 | 0,02 | 0,06 | 0,68 | 0,68 | 0,62 | 0,06 | 0,14 | 0,44 |
| 225 | Dha_Kin_pool_1 | | 5562 | PRKAA1 | S | 0,84 | 1,06 | 0,12 | 0,16 | 0,95 | 0,14 | 0,15 | 0,03 | 0,15 | 0,15 | 0,62 | 0,47 | 0,14 | -3,29 |
| 226 | Dha_Kin_pool_1 | | 8986 | RPS6KA4 | S | 0,63 | 0,56 | 0,25 | 0,29 | 0,59 | 0,27 | 0,05 | 0,03 | 0,46 | 0,46 | 0,62 | 0,15 | 0,14 | -1,08 |
| 227 | Dha_Kin_pool_1 | | 152110 | FLJ32685 | S | 0,77 | 0,76 | 0,45 | 0,50 | 0,77 | 0,48 | 0,01 | 0,03 | 0,62 | 0,62 | 0,62 | 0,01 | 0,14 | 0,04 |
| 228 | Dha_Kin_pool_1 | | 51701 | NLK | S | 0,98 | 1,00 | 0,52 | 0,51 | 0,99 | 0,51 | 0,02 | 0,01 | 0,52 | 0,52 | 0,62 | 0,10 | 0,14 | -0,68 |
| 229 | Dha_Kin_pool_1 | | 10420 | TESK2 | S | 0,90 | 1,03 | 0,65 | 0,61 | 0,96 | 0,63 | 0,09 | 0,03 | 0,65 | 0,65 | 0,62 | 0,03 | 0,14 | 0,24 |
| 230 | Dha_Kin_pool_1 | | 2045 | EPHA7 | S | 0,85 | 0,89 | 0,51 | 0,74 | 0,87 | 0,62 | 0,03 | 0,16 | 0,71 | 0,71 | 0,62 | 0,10 | 0,14 | 0,67 |
| 231 | Dha_Kin_pool_1 | | 140901 | STK35 | S | 0,81 | 0,74 | 0,51 | 0,42 | 0,77 | 0,46 | 0,05 | 0,06 | 0,60 | 0,60 | 0,62 | 0,02 | 0,14 | -0,13 |
| 232 | Dha_Kin_pool_1 | | 6794 | STK11 | S | 0,58 | 0,65 | 0,35 | 0,41 | 0,62 | 0,38 | 0,04 | 0,04 | 0,61 | 0,61 | 0,62 | 0,00 | 0,14 | -0,01 |
| 233 | Dha_Kin_pool_1 | | 3717 | JAK2 | S | 0,80 | 0,73 | 0,37 | 0,34 | 0,77 | 0,35 | 0,05 | 0,02 | 0,46 | 0,46 | 0,62 | 0,16 | 0,14 | -1,10 |
| 234 | Dha_Kin_pool_1 | | 5606 | MAP2K3 | S | 0,56 | 0,65 | 0,35 | 0,38 | 0,61 | 0,36 | 0,06 | 0,02 | 0,60 | 0,60 | 0,62 | 0,02 | 0,14 | -0,13 |
| 235 | Dha_Kin_pool_1 | | 3932 | LCK | S | 0,66 | 0,52 | 0,27 | 0,29 | 0,59 | 0,28 | 0,10 | 0,02 | 0,47 | 0,47 | 0,62 | 0,14 | 0,14 | -1,00 |
| 236 | Dha_Kin_pool_1 | | 22848 | AAK1 | S | 0,62 | 0,85 | 0,44 | 0,42 | 0,73 | 0,43 | 0,16 | 0,01 | 0,58 | 0,58 | 0,62 | 0,03 | 0,14 | -0,24 |
| 237 | Dha_Kin_pool_1 | | 3656 | IRAK2 | S | 0,79 | 0,70 | 0,53 | 0,45 | 0,75 | 0,49 | 0,06 | 0,06 | 0,65 | 0,65 | 0,62 | 0,04 | 0,14 | 0,27 |
| 238 | Dha_Kin_pool_1 | | 130399 | ACVR1C | S | 0,92 | 0,95 | 0,58 | 0,68 | 0,93 | 0,63 | 0,02 | 0,07 | 0,67 | 0,67 | 0,62 | 0,06 | 0,14 | 0,41 |
| 239 | Dha_Kin_pool_1 | E | | E | A | 0,55 | 0,61 | 0,60 | 0,67 | 0,58 | 0,63 | 0,04 | 0,04 | 1,09 | | 0,62 | | 0,14 | 3,32 |
| 240 | Dha_Kin_pool_1 | E | | E | A | 0,74 | 0,58 | 0,59 | 0,61 | 0,66 | 0,60 | 0,11 | 0,01 | 0,91 | | 0,62 | | 0,14 | 2,07 |
| 241 | Dha_Kin_pool_1 | | 91 | ACVR1B | S | 1,14 | 0,95 | 0,63 | 0,71 | 1,05 | 0,67 | 0,13 | 0,06 | 0,64 | 0,64 | 0,62 | 0,02 | 0,14 | 0,17 |
| 242 | Dha_Kin_pool_1 | | 8895 | CPNE3 | S | 1,01 | 1,14 | 0,63 | 0,79 | 1,08 | 0,71 | 0,10 | 0,11 | 0,66 | 0,66 | 0,62 | 0,04 | 0,14 | 0,30 |
| 243 | Dha_Kin_pool_1 | | 8536 | CAMK1 | S | 0,77 | 0,92 | 0,36 | 0,30 | 0,85 | 0,33 | 0,11 | 0,05 | 0,39 | 0,39 | 0,62 | 0,23 | 0,14 | -1,60 |
| 244 | Dha_Kin_pool_1 | | 4140 | MARK3 | S | 0,48 | 0,71 | 0,22 | 0,26 | 0,60 | 0,24 | 0,16 | 0,02 | 0,40 | 0,40 | 0,62 | 0,21 | 0,14 | -1,49 |
| 245 | Dha_Kin_pool_1 | | 5584 | PRKCI | S | 0,64 | 0,62 | 0,33 | 0,50 | 0,63 | 0,41 | 0,02 | 0,12 | 0,65 | 0,65 | 0,62 | 0,04 | 0,14 | 0,26 |
| 246 | Dha_Kin_pool_1 | | 9641 | IKBKE | S | 0,81 | 1,01 | 0,59 | 0,57 | 0,91 | 0,58 | 0,14 | 0,01 | 0,64 | 0,64 | 0,62 | 0,02 | 0,14 | 0,14 |
| 247 | Dha_Kin_pool_1 | | 122011 | CSNK1A1L | S | 0,87 | 0,83 | 0,43 | 0,55 | 0,85 | 0,49 | 0,02 | 0,08 | 0,58 | 0,58 | 0,62 | 0,04 | 0,14 | -0,27 |
| 248 | Dha_Kin_pool_1 | | 8394 | PIP5K1A | S | 0,63 | 0,72 | 0,43 | 0,58 | 0,67 | 0,50 | 0,06 | 0,11 | 0,75 | 0,75 | 0,62 | 0,13 | 0,14 | 0,91 |
| 249 | Dha_Kin_pool_1 | | 9263 | STK17A | S | 0,97 | 1,04 | 0,66 | 0,80 | 1,01 | 0,73 | 0,05 | 0,10 | 0,73 | 0,73 | 0,62 | 0,11 | 0,14 | 0,78 |
| 250 | Dha_Kin_pool_1 | | 56155 | TEX14 | S | 0,79 | 1,05 | 0,64 | 0,59 | 0,92 | 0,62 | 0,19 | 0,03 | 0,67 | 0,67 | 0,62 | 0,05 | 0,14 | 0,36 |
| 251 | Dha_Kin_pool_1 | | 9261 | MAPKAPK2 | S | 0,86 | 0,83 | 0,53 | 0,52 | 0,84 | 0,53 | 0,02 | 0,01 | 0,63 | 0,63 | 0,62 | 0,01 | 0,14 | 0,08 |
| 252 | Dha_Kin_pool_1 | | 4058 | LTK | S | 0,77 | 0,98 | 0,61 | 0,66 | 0,87 | 0,64 | 0,15 | 0,04 | 0,73 | 0,73 | 0,62 | 0,11 | 0,14 | 0,79 |
| 253 | Dha_Kin_pool_1 | | 8573 | CASK | S | 0,84 | 0,96 | 0,64 | 0,66 | 0,90 | 0,65 | 0,09 | 0,02 | 0,72 | 0,72 | 0,62 | 0,11 | 0,14 | 0,76 |
| 254 | Dha_Kin_pool_1 | | 9149 | DYRK1B | S | 0,30 | 0,32 | 0,24 | 0,21 | 0,31 | 0,23 | 0,02 | 0,02 | 0,73 | 0,73 | 0,62 | 0,11 | 0,14 | 0,78 |
| 255 | Dha_Kin_pool_1 | | 167359 | MGC42105 | S | 0,93 | 1,08 | 0,47 | 0,46 | 1,01 | 0,46 | 0,10 | 0,01 | 0,46 | 0,46 | 0,62 | 0,16 | 0,14 | -1,11 |
| 256 | Dha_Kin_pool_1 | | 1111 | CHEK1 | S | 0,23 | 0,34 | 0,18 | 0,17 | 0,28 | 0,18 | 0,08 | 0,00 | 0,62 | 0,62 | 0,62 | 0,01 | 0,14 | 0,06 |
| 257 | Dha_Kin_pool_1 | | 728642 | CDC2L2 | S | 0,13 | 0,17 | 0,08 | 0,11 | 0,15 | 0,10 | 0,03 | 0,02 | 0,66 | 0,66 | 0,62 | 0,05 | 0,14 | 0,33 |
| 258 | Dha_Kin_pool_1 | | 6788 | STK3 | S | 0,52 | 0,53 | 0,27 | 0,31 | 0,52 | 0,29 | 0,01 | 0,03 | 0,56 | 0,56 | 0,62 | 0,05 | 0,14 | -0,38 |
| 259 | Dha_Kin_pool_1 | | 984 | CDC2L1 | S | 0,41 | 0,39 | 0,19 | 0,21 | 0,40 | 0,20 | 0,01 | 0,01 | 0,49 | 0,49 | 0,62 | 0,12 | 0,14 | -0,87 |
| 260 | Dha_Kin_pool_1 | | 5609 | MAP2K7 | S | 0,92 | 0,97 | 0,76 | 0,90 | 0,94 | 0,83 | 0,03 | 0,10 | 0,88 | 0,88 | 0,62 | 0,27 | 0,14 | 1,88 |
| 261 | Dha_Kin_pool_1 | | 2051 | EPHB6 | S | 1,04 | 0,86 | 0,66 | 0,78 | 0,95 | 0,72 | 0,12 | 0,08 | 0,76 | 0,76 | 0,62 | 0,14 | 0,14 | 1,00 |
| 262 | Dha_Kin_pool_1 | | 4139 | MARK1 | S | 0,91 | 0,88 | 0,67 | 0,75 | 0,89 | 0,71 | 0,02 | 0,06 | 0,80 | 0,80 | 0,62 | 0,18 | 0,14 | 1,29 |
| 263 | Dha_Kin_pool_1 | E | | E | A | 0,64 | 0,72 | 0,66 | 0,69 | 0,68 | 0,67 | 0,06 | 0,02 | 0,99 | | 0,62 | | 0,14 | 2,62 |
| 264 | Dha_Kin_pool_1 | E | | E | A | 0,64 | 0,67 | 0,64 | 0,85 | 0,65 | 0,75 | 0,02 | 0,15 | 1,14 | | 0,62 | | 0,14 | 3,69 |
| 265 | Dha_Kin_pool_1 | | 388228 | SBK1 | S | 0,70 | 0,72 | 0,37 | 0,42 | 0,71 | 0,39 | 0,01 | 0,03 | 0,56 | 0,56 | 0,62 | 0,06 | 0,14 | -0,43 |
| 266 | Dha_Kin_pool_1 | | 5127 | PCTK1 | S | 0,46 | 0,43 | 0,27 | 0,27 | 0,44 | 0,27 | 0,02 | 0,01 | 0,61 | 0,61 | 0,62 | 0,01 | 0,14 | -0,04 |
| 267 | Dha_Kin_pool_1 | | 4216 | MAP3K4 | S | 0,78 | 0,77 | 0,09 | 0,12 | 0,78 | 0,11 | 0,01 | 0,02 | 0,14 | 0,14 | 0,62 | 0,48 | 0,14 | -3,34 |
| 268 | Dha_Kin_pool_1 | | 23012 | STK38L | S | 0,81 | 0,90 | 0,51 | 0,56 | 0,85 | 0,53 | 0,06 | 0,04 | 0,62 | 0,62 | 0,62 | 0,01 | 0,14 | 0,05 |
| 269 | Dha_Kin_pool_1 | | 83931 | MGC4796 | S | 0,91 | 1,04 | 0,45 | 0,47 | 0,97 | 0,46 | 0,10 | 0,02 | 0,47 | 0,47 | 0,62 | 0,14 | 0,14 | -1,01 |
| 270 | Dha_Kin_pool_1 | | 64080 | RBKS | S | 0,73 | 0,90 | 0,45 | 0,44 | 0,81 | 0,45 | 0,12 | 0,01 | 0,55 | 0,55 | 0,62 | 0,07 | 0,14 | -0,46 |
| 271 | Dha_Kin_pool_1 | | 5255 | PHKA1 | S | 0,79 | 0,79 | 0,61 | 0,60 | 0,79 | 0,61 | 0,00 | 0,01 | 0,77 | 0,77 | 0,62 | 0,15 | 0,14 | 1,07 |
| 272 | Dha_Kin_pool_1 | | 2081 | ERN1 | S | 0,89 | 0,71 | 0,56 | 0,66 | 0,80 | 0,61 | 0,13 | 0,07 | 0,76 | 0,76 | 0,62 | 0,15 | 0,14 | 1,03 |
| 273 | Dha_Kin_pool_1 | | 55359 | STYK1 | S | 1,01 | 1,03 | 0,31 | 0,35 | 1,02 | 0,33 | 0,01 | 0,03 | 0,32 | 0,32 | 0,62 | 0,29 | 0,14 | -2,06 |
| 274 | Dha_Kin_pool_1 | | 65018 | PINK1 | S | 0,88 | 0,79 | 0,36 | 0,54 | 0,84 | 0,45 | 0,06 | 0,13 | 0,53 | 0,53 | 0,62 | 0,08 | 0,14 | -0,58 |

| | | | | | | | | | | | | | | | | | | | |
|---|---|---|---|---|---|---|---|---|---|---|---|---|---|---|---|---|---|---|---|
| 275 | Dha_Kin_pool_1 | | 4593 | MUSK | S | 0,53 | 0,66 | 0,33 | 0,26 | 0,60 | 0,30 | 0,09 | 0,05 | 0,50 | 0,50 | 0,62 | 0,12 | 0,14 | -0,82 |
| 276 | Dha_Kin_pool_1 | | 9706 | ULK2 | S | 0,57 | 0,72 | 0,52 | 0,41 | 0,65 | 0,46 | 0,11 | 0,08 | 0,71 | 0,71 | 0,62 | 0,10 | 0,14 | 0,69 |
| 277 | Dha_Kin_pool_1 | | 79012 | MGC8407 | S | 0,96 | 0,88 | 0,51 | 0,55 | 0,92 | 0,53 | 0,06 | 0,03 | 0,57 | 0,57 | 0,62 | 0,04 | 0,14 | -0,29 |
| 278 | Dha_Kin_pool_1 | | 6790 | AURKA | S | 0,22 | 0,29 | 0,11 | 0,14 | 0,25 | 0,13 | 0,04 | 0,02 | 0,49 | 0,49 | 0,62 | 0,13 | 0,14 | -0,88 |
| 279 | Dha_Kin_pool_1 | | 10298 | PAK4 | S | 0,71 | 0,78 | 0,63 | 0,60 | 0,75 | 0,62 | 0,05 | 0,02 | 0,82 | 0,82 | 0,62 | 0,21 | 0,14 | 1,46 |
| 280 | Dha_Kin_pool_1 | | 5594 | MAPK1 | S | 0,74 | 1,13 | 0,61 | 0,82 | 0,93 | 0,71 | 0,27 | 0,14 | 0,77 | 0,77 | 0,62 | 0,15 | 0,14 | 1,06 |
| 281 | Dha_Kin_pool_1 | | 375449 | MAST4 | S | 0,82 | 0,78 | 0,45 | 0,53 | 0,80 | 0,49 | 0,03 | 0,06 | 0,62 | 0,62 | 0,62 | 0,00 | 0,14 | 0,00 |
| 282 | Dha_Kin_pool_1 | | 1160 | CKMT2 | S | 0,75 | 0,74 | 0,58 | 0,54 | 0,74 | 0,56 | 0,01 | 0,02 | 0,75 | 0,75 | 0,62 | 0,14 | 0,14 | 0,97 |
| 283 | Dha_Kin_pool_1 | | 9891 | ARK5 | S | 0,61 | 0,88 | 0,62 | 0,53 | 0,75 | 0,58 | 0,20 | 0,06 | 0,77 | 0,77 | 0,62 | 0,16 | 0,14 | 1,11 |
| 284 | Dha_Kin_pool_1 | | 8395 | PIP5K1B | S | 0,91 | 1,11 | 0,50 | 0,58 | 1,01 | 0,54 | 0,14 | 0,05 | 0,54 | 0,54 | 0,62 | 0,08 | 0,14 | -0,57 |
| 285 | Dha_Kin_pool_1 | | 1716 | DGUOK | S | 0,69 | 0,60 | 0,28 | 0,42 | 0,64 | 0,35 | 0,07 | 0,10 | 0,55 | 0,55 | 0,62 | 0,07 | 0,14 | -0,47 |
| 286 | Dha_Kin_pool_1 | | 79646 | PANK3 | S | 1,20 | 1,24 | 0,85 | 0,92 | 1,22 | 0,89 | 0,03 | 0,05 | 0,73 | 0,73 | 0,62 | 0,11 | 0,14 | 0,79 |
| 287 | Dha_Kin_pool_1 | E | | E | A | 0,60 | 0,77 | 0,70 | 0,71 | 0,68 | 0,70 | 0,12 | 0,01 | 1,03 | | 0,62 | | 0,14 | 2,87 |
| 288 | Dha_Kin_pool_1 | E | | E | A | 0,66 | 0,56 | 0,64 | 0,74 | 0,61 | 0,69 | 0,07 | 0,07 | 1,13 | | 0,62 | | 0,14 | 3,63 |
| 289 | Dha_Kin_pool_1 | | 6733 | SRPK2 | S | 1,33 | 1,13 | 0,82 | 0,91 | 1,23 | 0,86 | 0,14 | 0,06 | 0,70 | 0,70 | 0,62 | 0,09 | 0,14 | 0,61 |
| 290 | Dha_Kin_pool_1 | | 1196 | CLK2 | S | 0,72 | 0,58 | 0,15 | 0,34 | 0,65 | 0,25 | 0,10 | 0,14 | 0,38 | 0,38 | 0,62 | 0,24 | 0,14 | -1,66 |
| 291 | Dha_Kin_pool_1 | | 7048 | TGFBR2 | S | 0,70 | 0,86 | 0,23 | 0,23 | 0,78 | 0,23 | 0,11 | 0,00 | 0,30 | 0,30 | 0,62 | 0,32 | 0,14 | -2,23 |
| 292 | Dha_Kin_pool_1 | | 23617 | STK22B | S | 0,49 | 0,62 | 0,16 | 0,17 | 0,56 | 0,17 | 0,09 | 0,01 | 0,30 | 0,30 | 0,62 | 0,32 | 0,14 | -2,23 |
| 293 | Dha_Kin_pool_1 | | 2580 | GAK | S | 0,83 | 0,95 | 0,36 | 0,36 | 0,89 | 0,36 | 0,08 | 0,00 | 0,41 | 0,41 | 0,62 | 0,21 | 0,14 | -1,45 |
| 294 | Dha_Kin_pool_1 | | 6196 | RPS6KA2 | S | 0,88 | 0,84 | 0,29 | 0,36 | 0,86 | 0,32 | 0,03 | 0,05 | 0,38 | 0,38 | 0,62 | 0,24 | 0,14 | -1,69 |
| 295 | Dha_Kin_pool_1 | | 92 | ACVR2 | S | 0,78 | 0,65 | 0,44 | 0,46 | 0,71 | 0,45 | 0,10 | 0,02 | 0,63 | 0,63 | 0,62 | 0,01 | 0,14 | 0,09 |
| 296 | Dha_Kin_pool_1 | | 65267 | PRKWNK3 | S | 0,91 | 1,05 | 0,69 | 0,70 | 0,98 | 0,69 | 0,10 | 0,01 | 0,70 | 0,70 | 0,62 | 0,09 | 0,14 | 0,62 |
| 297 | Dha_Kin_pool_1 | | 5590 | PRKCZ | S | 0,67 | 0,85 | 0,56 | 0,74 | 0,76 | 0,65 | 0,13 | 0,13 | 0,86 | 0,86 | 0,62 | 0,24 | 0,14 | 1,69 |
| 298 | Dha_Kin_pool_1 | | 3702 | ITK | S | 0,86 | 1,07 | 0,56 | 0,58 | 0,97 | 0,57 | 0,15 | 0,02 | 0,59 | 0,59 | 0,62 | 0,03 | 0,14 | -0,18 |
| 299 | Dha_Kin_pool_1 | | 84254 | CAMKK1 | S | 0,95 | 1,04 | 0,46 | 0,69 | 1,00 | 0,57 | 0,06 | 0,16 | 0,58 | 0,58 | 0,62 | 0,04 | 0,14 | -0,28 |
| 300 | Dha_Kin_pool_1 | | 27148 | STK36 | S | 0,63 | 0,80 | 0,30 | 0,34 | 0,71 | 0,32 | 0,12 | 0,03 | 0,45 | 0,45 | 0,62 | 0,16 | 0,14 | -1,14 |
| 301 | Dha_Kin_pool_1 | | 5599 | MAPK8 | S | 0,84 | 0,76 | 0,27 | 0,35 | 0,80 | 0,31 | 0,06 | 0,06 | 0,39 | 0,39 | 0,62 | 0,23 | 0,14 | -1,60 |
| 302 | Dha_Kin_pool_1 | | 7046 | TGFBR1 | S | 1,06 | 1,16 | 0,89 | 0,97 | 1,11 | 0,93 | 0,07 | 0,05 | 0,84 | 0,84 | 0,62 | 0,22 | 0,14 | 1,54 |
| 303 | Dha_Kin_pool_1 | | 5597 | MAPK6 | S | 0,72 | 0,77 | 0,60 | 0,57 | 0,74 | 0,59 | 0,03 | 0,02 | 0,79 | 0,79 | 0,62 | 0,17 | 0,14 | 1,22 |
| 304 | Dha_Kin_pool_1 | | 131890 | GRK7 | S | 0,99 | 0,98 | 0,95 | 0,93 | 0,98 | 0,94 | 0,00 | 0,01 | 0,95 | 0,95 | 0,62 | 0,34 | 0,14 | 2,36 |
| 305 | Dha_Kin_pool_1 | | 156 | ADRBK1 | S | 0,61 | 0,80 | 0,79 | 0,72 | 0,70 | 0,76 | 0,14 | 0,05 | 1,07 | 1,07 | 0,62 | 0,46 | 0,14 | 3,22 |
| 306 | Dha_Kin_pool_1 | | 8558 | CDK10 | S | 0,55 | 0,90 | 0,70 | 0,55 | 0,72 | 0,62 | 0,25 | 0,11 | 0,87 | 0,87 | 0,62 | 0,25 | 0,14 | 1,75 |
| 307 | Dha_Kin_pool_1 | | 1026 | CDKN1A | S | 0,63 | 0,55 | 0,54 | 0,70 | 0,59 | 0,62 | 0,06 | 0,11 | 1,04 | 1,04 | 0,62 | 0,43 | 0,14 | 3,00 |
| 308 | Dha_Kin_pool_1 | | 80025 | PANK2 | S | 0,88 | 1,00 | 0,67 | 0,73 | 0,94 | 0,70 | 0,09 | 0,04 | 0,75 | 0,75 | 0,62 | 0,13 | 0,14 | 0,93 |
| 309 | Dha_Kin_pool_1 | | 2042 | EPHA3 | S | 0,95 | 1,04 | 0,81 | 0,94 | 0,99 | 0,88 | 0,06 | 0,10 | 0,88 | 0,88 | 0,62 | 0,26 | 0,14 | 1,86 |
| 310 | Dha_Kin_pool_1 | | 9807 | IHPK1 | S | 0,93 | 0,92 | 0,77 | 0,77 | 0,93 | 0,77 | 0,01 | 0,00 | 0,83 | 0,83 | 0,62 | 0,22 | 0,14 | 1,52 |
| 311 | Dha_Kin_pool_1 | E | | E | A | 0,66 | 0,72 | 0,67 | 0,88 | 0,69 | 0,78 | 0,04 | 0,15 | 1,12 | | 0,62 | | 0,14 | 3,56 |
| 312 | Dha_Kin_pool_1 | E | | E | A | 0,64 | 0,61 | 0,71 | 0,81 | 0,63 | 0,76 | 0,02 | 0,07 | 1,22 | | 0,62 | | 0,14 | 4,22 |
| 313 | Dha_Kin_pool_1 | | 91461 | LOC91461 | S | 0,67 | 0,66 | 0,42 | 0,45 | 0,67 | 0,44 | 0,00 | 0,02 | 0,66 | 0,66 | 0,62 | 0,04 | 0,14 | 0,28 |
| 314 | Dha_Kin_pool_1 | | 157285 | DKFZP761P0423 | S | 0,96 | 1,07 | 0,82 | 1,04 | 1,02 | 0,93 | 0,08 | 0,15 | 0,91 | 0,91 | 0,62 | 0,30 | 0,14 | 2,09 |
| 315 | Dha_Kin_pool_1 | | 25778 | DUSTYPK | S | 0,92 | 0,83 | 0,57 | 0,68 | 0,88 | 0,63 | 0,07 | 0,07 | 0,71 | 0,71 | 0,62 | 0,10 | 0,14 | 0,69 |
| 316 | Dha_Kin_pool_1 | | 5165 | PDK3 | S | 0,70 | 0,92 | 0,62 | 0,74 | 0,81 | 0,68 | 0,16 | 0,08 | 0,84 | 0,84 | 0,62 | 0,22 | 0,14 | 1,58 |
| 317 | Dha_Kin_pool_1 | | 1606 | DGKA | S | 0,29 | 0,48 | 0,41 | 0,35 | 0,39 | 0,38 | 0,14 | 0,05 | 0,98 | 0,98 | 0,62 | 0,37 | 0,14 | 2,59 |
| 318 | Dha_Kin_pool_1 | | 64768 | C9ORF12 | S | 0,38 | 0,60 | 0,29 | 0,39 | 0,49 | 0,34 | 0,16 | 0,07 | 0,69 | 0,69 | 0,62 | 0,07 | 0,14 | 0,48 |
| 319 | Dha_Kin_pool_1 | | 5753 | PTK6 | S | 0,65 | 0,71 | 0,30 | 0,36 | 0,68 | 0,33 | 0,04 | 0,04 | 0,48 | 0,48 | 0,62 | 0,14 | 0,14 | -0,96 |
| 320 | Dha_Kin_pool_1 | | 5058 | PAK1 | S | 0,70 | 0,62 | 0,24 | 0,25 | 0,66 | 0,24 | 0,05 | 0,01 | 0,37 | 0,37 | 0,62 | 0,25 | 0,14 | -1,74 |
| 321 | Dha_Kin_pool_1 | | 6259 | RYK | S | 0,95 | 1,06 | 0,65 | 0,66 | 1,01 | 0,65 | 0,08 | 0,00 | 0,65 | 0,65 | 0,62 | 0,03 | 0,14 | 0,23 |
| 322 | Dha_Kin_pool_1 | | 8566 | PDXK | S | 0,83 | 0,98 | 0,58 | 0,70 | 0,91 | 0,64 | 0,10 | 0,08 | 0,70 | 0,70 | 0,62 | 0,09 | 0,14 | 0,62 |
| 323 | Dha_Kin_pool_1 | | 7006 | TEC | S | 0,88 | 1,04 | 0,72 | 0,71 | 0,96 | 0,71 | 0,11 | 0,01 | 0,74 | 0,74 | 0,62 | 0,13 | 0,14 | 0,89 |
| 324 | Dha_Kin_pool_1 | | 26289 | AK5 | S | 0,62 | 0,86 | 0,24 | 0,31 | 0,74 | 0,28 | 0,17 | 0,05 | 0,38 | 0,38 | 0,62 | 0,24 | 0,14 | -1,68 |
| 325 | Dha_Kin_pool_1 | | 23139 | MAST2 | S | 0,80 | 0,90 | 0,62 | 0,61 | 0,85 | 0,61 | 0,07 | 0,00 | 0,72 | 0,72 | 0,62 | 0,10 | 0,14 | 0,73 |
| 326 | Dha_Kin_pool_1 | | 7465 | WEE1 | S | 0,24 | 0,24 | 0,09 | 0,11 | 0,24 | 0,10 | 0,00 | 0,02 | 0,43 | 0,43 | 0,62 | 0,19 | 0,14 | -1,33 |
| 327 | Dha_Kin_pool_1 | | 1021 | CDK6 | S | 0,66 | 0,98 | 0,49 | 0,50 | 0,82 | 0,50 | 0,23 | 0,01 | 0,60 | 0,60 | 0,62 | 0,01 | 0,14 | -0,09 |
| 328 | Dha_Kin_pool_1 | | 5588 | PRKCQ | S | 1,07 | 1,20 | 0,69 | 0,71 | 1,14 | 0,70 | 0,09 | 0,01 | 0,62 | 0,62 | 0,62 | 0,00 | 0,14 | 0,01 |
| 329 | Dha_Kin_pool_1 | | 284086 | NEK8 | S | 0,80 | 0,96 | 0,50 | 0,54 | 0,88 | 0,52 | 0,12 | 0,03 | 0,59 | 0,59 | 0,62 | 0,03 | 0,14 | -0,20 |

| | | | | | | | | | | | | | | | | | | | |
|---|---|---|---|---|---|---|---|---|---|---|---|---|---|---|---|---|---|---|---|
| 330 | Dha_Kin_pool_1 | | 2710 | GK | S | 0,82 | 1,02 | 0,53 | 0,65 | 0,92 | 0,59 | 0,14 | 0,09 | 0,64 | 0,64 | 0,62 | 0,02 | 0,14 | 0,14 |
| 331 | Dha_Kin_pool_1 | | 283455 | KSR2 | S | 0,68 | 0,72 | 0,54 | 0,67 | 0,70 | 0,61 | 0,03 | 0,09 | 0,87 | 0,87 | 0,62 | 0,25 | 0,14 | 1,76 |
| 332 | Dha_Kin_pool_1 | | 4920 | ROR2 | S | 1,14 | 0,99 | 0,76 | 0,92 | 1,06 | 0,84 | 0,10 | 0,12 | 0,79 | 0,79 | 0,62 | 0,18 | 0,14 | 1,23 |
| 333 | Dha_Kin_pool_1 | | 79837 | PIP5K2C | S | 0,85 | 0,89 | 0,61 | 0,98 | 0,87 | 0,79 | 0,03 | 0,26 | 0,91 | 0,91 | 0,62 | 0,30 | 0,14 | 2,08 |
| 334 | Dha_Kin_pool_1 | | 90 | ACVR1 | S | 0,69 | 1,00 | 0,56 | 0,67 | 0,85 | 0,62 | 0,22 | 0,08 | 0,73 | 0,73 | 0,62 | 0,11 | 0,14 | 0,81 |
| 335 | Dha_Kin_pool_1 | E | | E | A | 0,67 | 0,66 | 0,57 | 0,57 | 0,67 | 0,57 | 0,01 | 0,00 | 0,86 | | 0,62 | | 0,14 | 1,70 |
| 336 | Dha_Kin_pool_1 | E | | E | A | 0,59 | 0,66 | 0,59 | 0,74 | 0,63 | 0,67 | 0,05 | 0,10 | 1,06 | | 0,62 | | 0,14 | 3,12 |
| 337 | Dha_Kin_pool_1 | | 5567 | PRKACB | S | 1,18 | 1,09 | 0,69 | 0,74 | 1,13 | 0,71 | 0,06 | 0,04 | 0,63 | 0,63 | 0,62 | 0,01 | 0,14 | 0,08 |
| 338 | Dha_Kin_pool_1 | | 27347 | STK39 | S | 1,15 | 1,22 | 0,61 | 0,67 | 1,18 | 0,64 | 0,05 | 0,05 | 0,54 | 0,54 | 0,62 | 0,07 | 0,14 | -0,51 |
| 339 | Dha_Kin_pool_1 | | 4215 | MAP3K3 | S | 0,31 | 0,45 | 0,13 | 0,13 | 0,38 | 0,13 | 0,09 | 0,00 | 0,35 | 0,35 | 0,62 | 0,27 | 0,14 | -1,87 |
| 340 | Dha_Kin_pool_1 | | 4830 | NME1 | S | 0,90 | 0,78 | 0,49 | 0,55 | 0,84 | 0,52 | 0,08 | 0,05 | 0,62 | 0,62 | 0,62 | 0,00 | 0,14 | 0,02 |
| 341 | Dha_Kin_pool_1 | | 8711 | TNK1 | S | 0,67 | 1,02 | 0,61 | 0,61 | 0,85 | 0,61 | 0,24 | 0,00 | 0,72 | 0,72 | 0,62 | 0,11 | 0,14 | 0,75 |
| 342 | Dha_Kin_pool_1 | | 9024 | STK29 | S | 0,80 | 1,18 | 0,57 | 0,63 | 0,99 | 0,60 | 0,27 | 0,04 | 0,61 | 0,61 | 0,62 | 0,01 | 0,14 | -0,07 |
| 343 | Dha_Kin_pool_1 | | 4916 | NTRK3 | S | 0,86 | 0,97 | 0,56 | 0,69 | 0,91 | 0,63 | 0,07 | 0,09 | 0,68 | 0,68 | 0,62 | 0,07 | 0,14 | 0,48 |
| 344 | Dha_Kin_pool_1 | | 22853 | LMTK2 | S | 0,56 | 0,53 | 0,19 | 0,18 | 0,54 | 0,18 | 0,02 | 0,01 | 0,34 | 0,34 | 0,62 | 0,28 | 0,14 | -1,97 |
| 345 | Dha_Kin_pool_1 | | 1841 | DTYMK | S | 0,93 | 0,88 | 0,54 | 0,56 | 0,90 | 0,55 | 0,03 | 0,02 | 0,61 | 0,61 | 0,62 | 0,01 | 0,14 | -0,06 |
| 346 | Dha_Kin_pool_1 | | 4145 | MATK | S | 0,58 | 0,68 | 0,33 | 0,39 | 0,63 | 0,36 | 0,07 | 0,04 | 0,58 | 0,58 | 0,62 | 0,04 | 0,14 | -0,28 |
| 347 | Dha_Kin_pool_1 | | 6198 | RPS6KB1 | S | 0,85 | 0,69 | 0,32 | 0,38 | 0,77 | 0,35 | 0,11 | 0,04 | 0,46 | 0,46 | 0,62 | 0,16 | 0,14 | -1,12 |
| 348 | Dha_Kin_pool_1 | | 55312 | RFK | S | 0,92 | 0,84 | 0,55 | 0,79 | 0,88 | 0,67 | 0,06 | 0,17 | 0,76 | 0,76 | 0,62 | 0,15 | 0,14 | 1,02 |
| 349 | Dha_Kin_pool_1 | | 146057 | TTBK2 | S | 1,13 | 0,80 | 0,47 | 0,41 | 0,96 | 0,44 | 0,23 | 0,04 | 0,46 | 0,46 | 0,62 | 0,16 | 0,14 | -1,11 |
| 350 | Dha_Kin_pool_1 | | 8476 | CDC42BPA | S | 0,58 | 0,53 | 0,27 | 0,32 | 0,55 | 0,29 | 0,04 | 0,04 | 0,53 | 0,53 | 0,62 | 0,09 | 0,14 | -0,62 |
| 351 | Dha_Kin_pool_1 | | 83903 | GSG2 | S | 0,74 | 0,84 | 0,40 | 0,37 | 0,79 | 0,39 | 0,07 | 0,02 | 0,49 | 0,49 | 0,62 | 0,13 | 0,14 | -0,89 |
| 352 | Dha_Kin_pool_1 | | 204851 | HIPK1 | S | 0,87 | 0,87 | 0,51 | 0,61 | 0,87 | 0,56 | 0,00 | 0,07 | 0,65 | 0,65 | 0,62 | 0,03 | 0,14 | 0,22 |
| 353 | Dha_Kin_pool_1 | | 4296 | MAP3K11 | S | 0,70 | 0,89 | 0,49 | 0,50 | 0,80 | 0,50 | 0,14 | 0,00 | 0,62 | 0,62 | 0,62 | 0,01 | 0,14 | 0,06 |
| 354 | Dha_Kin_pool_1 | | 9212 | AURKB | S | 0,40 | 0,50 | 0,17 | 0,15 | 0,45 | 0,16 | 0,07 | 0,01 | 0,36 | 0,36 | 0,62 | 0,26 | 0,14 | -1,81 |
| 355 | Dha_Kin_pool_1 | | 3985 | LIMK2 | S | 0,86 | 0,73 | 0,64 | 0,58 | 0,80 | 0,61 | 0,09 | 0,04 | 0,77 | 0,77 | 0,62 | 0,15 | 0,14 | 1,07 |
| 356 | Dha_Kin_pool_1 | | 2645 | GCK | S | 0,66 | 0,59 | 0,32 | 0,47 | 0,63 | 0,39 | 0,05 | 0,11 | 0,63 | 0,63 | 0,62 | 0,01 | 0,14 | 0,08 |
| 357 | Dha_Kin_pool_1 | | 2048 | EPHB2 | S | 0,29 | 0,22 | 0,15 | 0,15 | 0,25 | 0,15 | 0,05 | 0,00 | 0,60 | 0,60 | 0,62 | 0,01 | 0,14 | -0,08 |
| 358 | Dha_Kin_pool_1 | | 23678 | SGKL | S | 0,86 | 0,92 | 0,70 | 0,72 | 0,89 | 0,71 | 0,04 | 0,01 | 0,80 | 0,80 | 0,62 | 0,19 | 0,14 | 1,33 |
| 359 | Dha_Kin_pool_1 | E | | E | A | 0,54 | 0,65 | 0,61 | 0,53 | 0,60 | 0,57 | 0,08 | 0,06 | 0,96 | | 0,62 | | 0,14 | 2,39 |
| 360 | Dha_Kin_pool_1 | E | | E | A | 0,54 | 0,54 | 0,54 | 0,57 | 0,54 | 0,56 | 0,00 | 0,02 | 1,04 | | 0,62 | | 0,14 | 2,97 |
| 361 | Dha_Kin_pool_1 | | 5979 | RET | S | 1,00 | 0,84 | 0,28 | 0,23 | 0,92 | 0,26 | 0,11 | 0,03 | 0,28 | 0,28 | 0,62 | 0,34 | 0,14 | -2,36 |
| 362 | Dha_Kin_pool_1 | | 5608 | MAP2K6 | S | 1,06 | 1,07 | 0,49 | 0,48 | 1,07 | 0,49 | 0,01 | 0,01 | 0,45 | 0,45 | 0,62 | 0,16 | 0,14 | -1,13 |
| 363 | Dha_Kin_pool_1 | | 26750 | RPS6KC1 | S | 0,23 | 0,27 | 0,07 | 0,11 | 0,25 | 0,09 | 0,03 | 0,03 | 0,36 | 0,36 | 0,62 | 0,26 | 0,14 | -1,82 |
| 364 | Dha_Kin_pool_1 | | 29922 | NME7 | S | 0,91 | 1,05 | 0,50 | 0,48 | 0,98 | 0,49 | 0,10 | 0,01 | 0,50 | 0,50 | 0,62 | 0,12 | 0,14 | -0,84 |
| 365 | Dha_Kin_pool_1 | | 4294 | MAP3K10 | S | 1,10 | 0,85 | 0,53 | 0,73 | 0,98 | 0,63 | 0,17 | 0,14 | 0,64 | 0,64 | 0,62 | 0,03 | 0,14 | 0,20 |
| 366 | Dha_Kin_pool_1 | | 83549 | UCK1 | S | 1,13 | 1,04 | 0,76 | 0,74 | 1,09 | 0,75 | 0,06 | 0,02 | 0,69 | 0,69 | 0,62 | 0,07 | 0,14 | 0,52 |
| 367 | Dha_Kin_pool_1 | | 2868 | GRK4 | S | 0,59 | 0,54 | 0,19 | 0,18 | 0,57 | 0,18 | 0,04 | 0,00 | 0,33 | 0,33 | 0,62 | 0,29 | 0,14 | -2,04 |
| 368 | Dha_Kin_pool_1 | | 340156 | LOC340156 | S | 0,35 | 0,37 | 0,20 | 0,21 | 0,36 | 0,21 | 0,01 | 0,00 | 0,58 | 0,58 | 0,62 | 0,04 | 0,14 | -0,29 |
| 369 | Dha_Kin_pool_1 | | 5613 | PRKX | S | 0,80 | 0,73 | 0,27 | 0,35 | 0,76 | 0,31 | 0,05 | 0,06 | 0,41 | 0,41 | 0,62 | 0,21 | 0,14 | -1,48 |
| 370 | Dha_Kin_pool_1 | | 10020 | GNE | S | 0,68 | 0,78 | 0,52 | 0,45 | 0,73 | 0,48 | 0,07 | 0,05 | 0,66 | 0,66 | 0,62 | 0,05 | 0,14 | 0,33 |
| 371 | Dha_Kin_pool_1 | | 4598 | MVK | S | 0,58 | 0,55 | 0,34 | 0,40 | 0,56 | 0,37 | 0,02 | 0,04 | 0,66 | 0,66 | 0,62 | 0,04 | 0,14 | 0,30 |
| 372 | Dha_Kin_pool_1 | | 1020 | CDK5 | S | 0,80 | 0,64 | 0,47 | 0,55 | 0,72 | 0,51 | 0,12 | 0,06 | 0,71 | 0,71 | 0,62 | 0,09 | 0,14 | 0,64 |
| 373 | Dha_Kin_pool_1 | | 79934 | ADCK4 | S | 0,83 | 0,82 | 0,63 | 0,63 | 0,82 | 0,63 | 0,00 | 0,00 | 0,77 | 0,77 | 0,62 | 0,15 | 0,14 | 1,06 |
| 374 | Dha_Kin_pool_1 | | 27102 | HRI | S | 0,83 | 0,83 | 0,46 | 0,50 | 0,83 | 0,48 | 0,00 | 0,02 | 0,58 | 0,58 | 0,62 | 0,04 | 0,14 | -0,27 |
| 375 | Dha_Kin_pool_1 | | 2263 | FGFR2 | S | 0,59 | 0,73 | 0,42 | 0,38 | 0,66 | 0,40 | 0,09 | 0,03 | 0,60 | 0,60 | 0,62 | 0,01 | 0,14 | -0,08 |
| 376 | Dha_Kin_pool_1 | | 2065 | ERBB3 | S | 0,37 | 0,56 | 0,30 | 0,31 | 0,46 | 0,31 | 0,13 | 0,01 | 0,66 | 0,66 | 0,62 | 0,04 | 0,14 | 0,31 |
| 377 | Dha_Kin_pool_1 | | 4752 | NEK3 | S | 0,75 | 0,73 | 0,38 | 0,44 | 0,74 | 0,41 | 0,01 | 0,04 | 0,55 | 0,55 | 0,62 | 0,06 | 0,14 | -0,44 |
| 378 | Dha_Kin_pool_1 | | 10087 | COL4A3BP | S | 0,47 | 0,69 | 0,13 | 0,18 | 0,58 | 0,15 | 0,15 | 0,04 | 0,26 | 0,26 | 0,62 | 0,35 | 0,14 | -2,47 |
| 379 | Dha_Kin_pool_1 | | 8621 | CDC2L5 | S | 0,98 | 0,94 | 0,68 | 0,70 | 0,96 | 0,69 | 0,03 | 0,01 | 0,72 | 0,72 | 0,62 | 0,10 | 0,14 | 0,74 |
| 380 | Dha_Kin_pool_1 | | 238 | ALK | S | 0,95 | 0,88 | 0,65 | 0,68 | 0,91 | 0,66 | 0,04 | 0,02 | 0,72 | 0,72 | 0,62 | 0,11 | 0,14 | 0,75 |
| 381 | Dha_Kin_pool_1 | | 10290 | SPEG | S | 0,88 | 0,95 | 0,57 | 0,61 | 0,92 | 0,59 | 0,05 | 0,02 | 0,64 | 0,64 | 0,62 | 0,03 | 0,14 | 0,18 |
| 382 | Dha_Kin_pool_1 | | 112858 | TP53RK | S | 1,02 | 0,91 | 0,35 | 0,29 | 0,96 | 0,32 | 0,08 | 0,04 | 0,33 | 0,33 | 0,62 | 0,29 | 0,14 | -2,01 |
| 383 | Dha_Kin_pool_1 | E | | E | A | 0,58 | 0,46 | 0,54 | 0,52 | 0,52 | 0,53 | 0,08 | 0,01 | 1,03 | | 0,62 | | 0,14 | 2,87 |
| 384 | Dha_Kin_pool_1 | E | | E | K | 0,02 | 0,01 | 0,02 | 0,01 | 0,01 | 0,02 | 0,00 | 0,00 | 1,09 | | 0,62 | | 0,14 | 3,36 |

| | | | | | | | | | | | | | | | | | | | |
|---|---|---|---|---|---|---|---|---|---|---|---|---|---|---|---|---|---|---|---|
| 385 | Dha_Kin_pool_2 | | 5603 | MAPK13 | S | 0,70 | 0,59 | 0,43 | 0,27 | 0,65 | 0,35 | 0,08 | 0,11 | 0,54 | 0,54 | 0,64 | 0,10 | 0,13 | -0,73 |
| 386 | Dha_Kin_pool_2 | | 7867 | MAPKAPK3 | S | 0,89 | 0,67 | 0,41 | 0,44 | 0,78 | 0,43 | 0,16 | 0,03 | 0,54 | 0,54 | 0,64 | 0,10 | 0,13 | -0,73 |
| 387 | Dha_Kin_pool_2 | | 93 | ACVR2B | S | 0,77 | 0,74 | 0,23 | 0,29 | 0,76 | 0,26 | 0,03 | 0,04 | 0,34 | 0,34 | 0,64 | 0,30 | 0,13 | -2,26 |
| 388 | Dha_Kin_pool_2 | | 1956 | EGFR | S | 0,79 | 0,76 | 0,69 | 0,82 | 0,77 | 0,76 | 0,03 | 0,09 | 0,98 | 0,98 | 0,64 | 0,33 | 0,13 | 2,46 |
| 389 | Dha_Kin_pool_2 | | 6098 | ROS1 | S | 0,97 | 0,95 | 0,72 | 0,68 | 0,96 | 0,70 | 0,02 | 0,03 | 0,73 | 0,73 | 0,64 | 0,09 | 0,13 | 0,65 |
| 390 | Dha_Kin_pool_2 | | 3101 | HK3 | S | 0,84 | 0,86 | 0,58 | 0,69 | 0,85 | 0,63 | 0,01 | 0,08 | 0,75 | 0,75 | 0,64 | 0,10 | 0,13 | 0,75 |
| 391 | Dha_Kin_pool_2 | | 4921 | DDR2 | S | 0,84 | 0,86 | 0,39 | 0,45 | 0,85 | 0,42 | 0,02 | 0,04 | 0,49 | 0,49 | 0,64 | 0,15 | 0,13 | -1,12 |
| 392 | Dha_Kin_pool_2 | | 23604 | DAPK2 | S | 0,80 | 0,74 | 0,55 | 0,50 | 0,77 | 0,53 | 0,04 | 0,04 | 0,68 | 0,68 | 0,64 | 0,04 | 0,13 | 0,27 |
| 393 | Dha_Kin_pool_2 | | 3705 | ITPK1 | S | 0,73 | 0,65 | 0,47 | 0,40 | 0,69 | 0,43 | 0,05 | 0,05 | 0,62 | 0,62 | 0,64 | 0,02 | 0,13 | -0,16 |
| 394 | Dha_Kin_pool_2 | | 1969 | EPHA2 | S | 0,85 | 0,68 | 0,48 | 0,42 | 0,76 | 0,45 | 0,12 | 0,05 | 0,59 | 0,59 | 0,64 | 0,05 | 0,13 | -0,39 |
| 395 | Dha_Kin_pool_2 | | 5260 | PHKG1 | S | 0,88 | 0,77 | 0,54 | 0,42 | 0,83 | 0,48 | 0,07 | 0,09 | 0,58 | 0,58 | 0,64 | 0,06 | 0,13 | -0,46 |
| 396 | Dha_Kin_pool_2 | | 160851 | DGKH | S | 0,34 | 0,26 | 0,20 | 0,22 | 0,30 | 0,21 | 0,06 | 0,02 | 0,70 | 0,70 | 0,64 | 0,06 | 0,13 | 0,45 |
| 397 | Dha_Kin_pool_2 | | 2444 | FRK | S | 1,09 | 0,94 | 0,32 | 0,62 | 1,01 | 0,47 | 0,11 | 0,21 | 0,46 | 0,46 | 0,64 | 0,18 | 0,13 | -1,35 |
| 398 | Dha_Kin_pool_2 | | 5291 | PIK3CB | S | 0,73 | 0,83 | 0,33 | 0,56 | 0,78 | 0,45 | 0,07 | 0,16 | 0,57 | 0,57 | 0,64 | 0,07 | 0,13 | -0,53 |
| 399 | Dha_Kin_pool_2 | | 8814 | CDKL1 | S | 0,79 | 0,69 | 0,43 | 0,46 | 0,74 | 0,44 | 0,07 | 0,02 | 0,60 | 0,60 | 0,64 | 0,04 | 0,13 | -0,31 |
| 400 | Dha_Kin_pool_2 | | 4915 | NTRK2 | S | 0,56 | 0,54 | 0,33 | 0,32 | 0,55 | 0,32 | 0,01 | 0,00 | 0,59 | 0,59 | 0,64 | 0,06 | 0,13 | -0,42 |
| 401 | Dha_Kin_pool_2 | | 9625 | AATK | S | 0,80 | 0,73 | 0,46 | 0,48 | 0,77 | 0,47 | 0,04 | 0,01 | 0,61 | 0,61 | 0,64 | 0,03 | 0,13 | -0,24 |
| 402 | Dha_Kin_pool_2 | | 5582 | PRKCG | S | 0,87 | 0,77 | 0,31 | 0,33 | 0,82 | 0,32 | 0,07 | 0,01 | 0,39 | 0,39 | 0,64 | 0,25 | 0,13 | -1,87 |
| 403 | Dha_Kin_pool_2 | | 11329 | STK38 | S | 0,78 | 0,67 | 0,27 | 0,34 | 0,72 | 0,30 | 0,08 | 0,05 | 0,42 | 0,42 | 0,64 | 0,23 | 0,13 | -1,67 |
| 404 | Dha_Kin_pool_2 | | 197258 | FUK | S | 0,69 | 0,59 | 0,36 | 0,31 | 0,64 | 0,34 | 0,07 | 0,04 | 0,52 | 0,52 | 0,64 | 0,12 | 0,13 | -0,89 |
| 405 | Dha_Kin_pool_2 | | 205 | AK3 | S | 0,98 | 0,90 | 0,46 | 0,50 | 0,94 | 0,48 | 0,06 | 0,03 | 0,51 | 0,51 | 0,64 | 0,13 | 0,13 | -0,99 |
| 406 | Dha_Kin_pool_2 | | 1459 | CSNK2A2 | S | 0,72 | 0,73 | 0,50 | 0,45 | 0,72 | 0,47 | 0,00 | 0,04 | 0,65 | 0,65 | 0,64 | 0,01 | 0,13 | 0,06 |
| 407 | Dha_Kin_pool_2 | E | | E | D | 0,91 | 0,89 | 0,95 | 0,92 | 0,90 | 0,94 | 0,01 | 0,02 | 1,04 | | 0,64 | | 0,13 | 2,96 |
| 408 | Dha_Kin_pool_2 | E | | E | K | 0,04 | 0,05 | 0,04 | 0,04 | 0,04 | 0,04 | 0,01 | 0,00 | 0,92 | | 0,64 | | 0,13 | 2,02 |
| 409 | Dha_Kin_pool_2 | | 7444 | VRK2 | S | 1,15 | 0,83 | 0,61 | 0,55 | 0,99 | 0,58 | 0,23 | 0,04 | 0,58 | 0,58 | 0,64 | 0,06 | 0,13 | -0,45 |
| 410 | Dha_Kin_pool_2 | | 369 | ARAF1 | S | 0,61 | 0,60 | 0,28 | 0,32 | 0,60 | 0,30 | 0,01 | 0,03 | 0,50 | 0,50 | 0,64 | 0,14 | 0,13 | -1,04 |
| 411 | Dha_Kin_pool_2 | | 613 | BCR | S | 0,86 | 0,80 | 0,42 | 0,61 | 0,83 | 0,51 | 0,04 | 0,14 | 0,62 | 0,62 | 0,64 | 0,02 | 0,13 | -0,18 |
| 412 | Dha_Kin_pool_2 | | 1633 | DCK | S | 0,75 | 0,76 | 0,30 | 0,45 | 0,75 | 0,38 | 0,01 | 0,11 | 0,50 | 0,50 | 0,64 | 0,15 | 0,13 | -1,08 |
| 413 | Dha_Kin_pool_2 | | 93627 | MGC16169 | S | 1,00 | 0,94 | 0,45 | 0,59 | 0,97 | 0,52 | 0,04 | 0,10 | 0,53 | 0,53 | 0,64 | 0,11 | 0,13 | -0,83 |
| 414 | Dha_Kin_pool_2 | | 9162 | DGKI | S | 0,80 | 0,77 | 0,48 | 0,50 | 0,79 | 0,49 | 0,02 | 0,01 | 0,62 | 0,62 | 0,64 | 0,02 | 0,13 | -0,16 |
| 415 | Dha_Kin_pool_2 | | 10654 | PMVK | S | 0,73 | 0,73 | 0,53 | 0,44 | 0,73 | 0,48 | 0,00 | 0,07 | 0,66 | 0,66 | 0,64 | 0,01 | 0,13 | 0,11 |
| 416 | Dha_Kin_pool_2 | | 2041 | EPHA1 | S | 0,85 | 0,92 | 0,74 | 0,59 | 0,88 | 0,66 | 0,05 | 0,11 | 0,75 | 0,75 | 0,64 | 0,11 | 0,13 | 0,80 |
| 417 | Dha_Kin_pool_2 | | 5894 | RAF1 | S | 0,84 | 0,76 | 0,65 | 0,52 | 0,80 | 0,59 | 0,06 | 0,09 | 0,73 | 0,73 | 0,64 | 0,09 | 0,13 | 0,68 |
| 418 | Dha_Kin_pool_2 | | 5580 | PRKCD | S | 0,79 | 0,76 | 0,59 | 0,35 | 0,78 | 0,47 | 0,02 | 0,17 | 0,60 | 0,60 | 0,64 | 0,04 | 0,13 | -0,30 |
| 419 | Dha_Kin_pool_2 | | 545 | ATR | S | 0,66 | 0,68 | 0,41 | 0,38 | 0,67 | 0,39 | 0,01 | 0,02 | 0,59 | 0,59 | 0,64 | 0,06 | 0,13 | -0,41 |
| 420 | Dha_Kin_pool_2 | | 815 | CAMK2A | S | 0,52 | 0,43 | 0,37 | 0,29 | 0,48 | 0,33 | 0,06 | 0,06 | 0,70 | 0,70 | 0,64 | 0,06 | 0,13 | 0,43 |
| 421 | Dha_Kin_pool_2 | | 3791 | KDR | S | 0,73 | 0,64 | 0,59 | 0,52 | 0,69 | 0,55 | 0,06 | 0,05 | 0,81 | 0,81 | 0,64 | 0,16 | 0,13 | 1,22 |
| 422 | Dha_Kin_pool_2 | | 23097 | CDK11 | S | 0,58 | 0,69 | 0,47 | 0,35 | 0,64 | 0,41 | 0,08 | 0,09 | 0,65 | 0,65 | 0,64 | 0,00 | 0,13 | 0,02 |
| 423 | Dha_Kin_pool_2 | | 1445 | CSK | S | 0,72 | 0,70 | 0,57 | 0,42 | 0,71 | 0,49 | 0,02 | 0,11 | 0,70 | 0,70 | 0,64 | 0,05 | 0,13 | 0,40 |
| 424 | Dha_Kin_pool_2 | | 157 | ADRBK2 | S | 0,80 | 0,81 | 0,49 | 0,53 | 0,80 | 0,51 | 0,01 | 0,03 | 0,63 | 0,63 | 0,64 | 0,01 | 0,13 | -0,09 |
| 425 | Dha_Kin_pool_2 | | 22858 | ICK | S | 0,44 | 0,46 | 0,22 | 0,21 | 0,45 | 0,22 | 0,01 | 0,00 | 0,48 | 0,48 | 0,64 | 0,16 | 0,13 | -1,18 |
| 426 | Dha_Kin_pool_2 | | 90956 | ADCK2 | S | 0,60 | 0,63 | 0,61 | 0,45 | 0,62 | 0,53 | 0,02 | 0,11 | 0,86 | 0,86 | 0,64 | 0,22 | 0,13 | 1,61 |
| 427 | Dha_Kin_pool_2 | | 55300 | PI4K2B | S | 0,87 | 0,86 | 0,45 | 0,46 | 0,87 | 0,46 | 0,00 | 0,01 | 0,53 | 0,53 | 0,64 | 0,12 | 0,13 | -0,86 |
| 428 | Dha_Kin_pool_2 | | 80122 | FLJ23074 | S | 0,98 | 0,84 | 0,70 | 0,58 | 0,91 | 0,64 | 0,10 | 0,08 | 0,70 | 0,70 | 0,64 | 0,06 | 0,13 | 0,44 |
| 429 | Dha_Kin_pool_2 | | 28996 | HIPK2 | S | 0,78 | 0,80 | 0,45 | 0,40 | 0,79 | 0,43 | 0,02 | 0,04 | 0,54 | 0,54 | 0,64 | 0,10 | 0,13 | -0,74 |
| 430 | Dha_Kin_pool_2 | | 81788 | SNARK | S | 0,46 | 0,41 | 0,19 | 0,19 | 0,43 | 0,19 | 0,04 | 0,00 | 0,44 | 0,44 | 0,64 | 0,20 | 0,13 | -1,48 |
| 431 | Dha_Kin_pool_2 | E | | E | D | 0,98 | 1,00 | 1,10 | 0,94 | 0,99 | 1,02 | 0,01 | 0,11 | 1,03 | | 0,64 | | 0,13 | 2,87 |
| 432 | Dha_Kin_pool_2 | E | | E | D | 0,94 | 1,06 | 0,79 | 0,93 | 1,00 | 0,86 | 0,08 | 0,10 | 0,86 | | 0,64 | | 0,13 | 1,63 |
| 433 | Dha_Kin_pool_2 | | 55351 | STK32B | S | 0,79 | 0,78 | 0,42 | 0,45 | 0,78 | 0,43 | 0,01 | 0,02 | 0,55 | 0,55 | 0,64 | 0,09 | 0,13 | -0,67 |
| 434 | Dha_Kin_pool_2 | | 5610 | PRKR | S | 0,91 | 0,97 | 0,56 | 0,61 | 0,94 | 0,58 | 0,04 | 0,03 | 0,62 | 0,62 | 0,64 | 0,02 | 0,13 | -0,16 |
| 435 | Dha_Kin_pool_2 | | 11035 | RIPK3 | S | 0,60 | 0,50 | 0,27 | 0,32 | 0,55 | 0,30 | 0,07 | 0,04 | 0,54 | 0,54 | 0,64 | 0,10 | 0,13 | -0,77 |
| 436 | Dha_Kin_pool_2 | | 54986 | ULK4 | S | 0,65 | 0,59 | 0,45 | 0,36 | 0,62 | 0,41 | 0,04 | 0,07 | 0,65 | 0,65 | 0,64 | 0,01 | 0,13 | 0,05 |
| 437 | Dha_Kin_pool_2 | | 56848 | SPHK2 | S | 0,56 | 0,47 | 0,15 | 0,16 | 0,51 | 0,15 | 0,07 | 0,01 | 0,30 | 0,30 | 0,64 | 0,34 | 0,13 | -2,55 |
| 438 | Dha_Kin_pool_2 | | 7016 | TESK1 | S | 0,58 | 0,56 | 0,33 | 0,33 | 0,57 | 0,33 | 0,02 | 0,00 | 0,58 | 0,58 | 0,64 | 0,07 | 0,13 | -0,49 |
| 439 | Dha_Kin_pool_2 | | 1453 | CSNK1D | S | 0,71 | 0,60 | 0,55 | 0,42 | 0,66 | 0,48 | 0,07 | 0,09 | 0,73 | 0,73 | 0,64 | 0,09 | 0,13 | 0,67 |

| | | | | | | | | | | | | | | | | | | | |
|---|---|---|---|---|---|---|---|---|---|---|---|---|---|---|---|---|---|---|---|
| 440 | Dha_Kin_pool_2 | | 8444 | DYRK3 | S | 0,64 | 0,56 | 0,45 | 0,29 | 0,60 | 0,37 | 0,05 | 0,11 | 0,62 | 0,62 | 0,64 | 0,02 | 0,13 | -0,14 |
| 441 | Dha_Kin_pool_2 | | 7297 | TYK2 | S | 0,90 | 0,77 | 0,46 | 0,50 | 0,84 | 0,48 | 0,09 | 0,03 | 0,58 | 0,58 | 0,64 | 0,07 | 0,13 | -0,50 |
| 442 | Dha_Kin_pool_2 | | 53904 | MYO3A | S | 1,00 | 0,44 | 0,71 | 0,71 | 0,72 | 0,71 | 0,40 | 0,01 | 0,99 | 0,99 | 0,64 | 0,34 | 0,13 | 2,54 |
| 443 | Dha_Kin_pool_2 | | 8877 | SPHK1 | S | 0,85 | 0,84 | 0,48 | 0,37 | 0,85 | 0,43 | 0,01 | 0,08 | 0,50 | 0,50 | 0,64 | 0,14 | 0,13 | -1,04 |
| 444 | Dha_Kin_pool_2 | | 10926 | ASK | S | 0,78 | 0,77 | 0,49 | 0,43 | 0,78 | 0,46 | 0,01 | 0,04 | 0,59 | 0,59 | 0,64 | 0,05 | 0,13 | -0,38 |
| 445 | Dha_Kin_pool_2 | | 5159 | PDGFRB | S | 0,50 | 0,36 | 0,26 | 0,18 | 0,43 | 0,22 | 0,10 | 0,06 | 0,51 | 0,51 | 0,64 | 0,13 | 0,13 | -0,98 |
| 446 | Dha_Kin_pool_2 | | 2268 | FGR | S | 0,52 | 0,42 | 0,33 | 0,25 | 0,47 | 0,29 | 0,07 | 0,06 | 0,63 | 0,63 | 0,64 | 0,01 | 0,13 | -0,10 |
| 447 | Dha_Kin_pool_2 | | 4919 | ROR1 | S | 0,76 | 0,63 | 0,76 | 0,51 | 0,70 | 0,64 | 0,09 | 0,18 | 0,92 | 0,92 | 0,64 | 0,27 | 0,13 | 2,03 |
| 448 | Dha_Kin_pool_2 | | 51755 | CRK7 | S | 0,67 | 0,53 | 0,39 | 0,26 | 0,60 | 0,32 | 0,10 | 0,09 | 0,54 | 0,54 | 0,64 | 0,10 | 0,13 | -0,75 |
| 449 | Dha_Kin_pool_2 | | 1460 | CSNK2B | S | 0,76 | 0,61 | 0,42 | 0,36 | 0,69 | 0,39 | 0,11 | 0,04 | 0,57 | 0,57 | 0,64 | 0,07 | 0,13 | -0,55 |
| 450 | Dha_Kin_pool_2 | | 2066 | ERBB4 | S | 0,65 | 0,64 | 0,46 | 0,31 | 0,64 | 0,38 | 0,00 | 0,10 | 0,60 | 0,60 | 0,64 | 0,05 | 0,13 | -0,34 |
| 451 | Dha_Kin_pool_2 | | 9088 | PKMYT1 | S | 0,22 | 0,21 | 0,15 | 0,09 | 0,21 | 0,12 | 0,01 | 0,04 | 0,56 | 0,56 | 0,64 | 0,08 | 0,13 | -0,58 |
| 452 | Dha_Kin_pool_2 | | 3551 | IKBKB | S | 0,48 | 0,45 | 0,34 | 0,32 | 0,46 | 0,33 | 0,02 | 0,02 | 0,71 | 0,71 | 0,64 | 0,07 | 0,13 | 0,53 |
| 453 | Dha_Kin_pool_2 | | 9175 | MAP3K13 | S | 0,74 | 0,68 | 0,34 | 0,27 | 0,71 | 0,30 | 0,04 | 0,05 | 0,43 | 0,43 | 0,64 | 0,22 | 0,13 | -1,59 |
| 454 | Dha_Kin_pool_2 | | 57118 | CAMK1D | S | 1,01 | 0,90 | 0,63 | 0,51 | 0,95 | 0,57 | 0,07 | 0,09 | 0,60 | 0,60 | 0,64 | 0,05 | 0,13 | -0,35 |
| 455 | Dha_Kin_pool_2 | E | | E | D | 1,04 | 0,95 | 1,00 | 0,74 | 0,99 | 0,87 | 0,06 | 0,18 | 0,88 | | 0,64 | | 0,13 | 1,73 |
| 456 | Dha_Kin_pool_2 | E | | E | D | 0,89 | 0,98 | 0,90 | 1,05 | 0,94 | 0,98 | 0,06 | 0,11 | 1,04 | | 0,64 | | 0,13 | 2,96 |
| 457 | Dha_Kin_pool_2 | | 11213 | IRAK3 | S | 1,04 | 1,14 | 0,64 | 0,67 | 1,09 | 0,65 | 0,07 | 0,02 | 0,60 | 0,60 | 0,64 | 0,04 | 0,13 | -0,31 |
| 458 | Dha_Kin_pool_2 | | 4342 | MOS | S | 0,80 | 0,84 | 0,54 | 0,42 | 0,82 | 0,48 | 0,03 | 0,08 | 0,59 | 0,59 | 0,64 | 0,05 | 0,13 | -0,39 |
| 459 | Dha_Kin_pool_2 | | 1027 | CDKN1B | S | 0,66 | 0,54 | 0,31 | 0,24 | 0,60 | 0,27 | 0,09 | 0,04 | 0,46 | 0,46 | 0,64 | 0,19 | 0,13 | -1,37 |
| 460 | Dha_Kin_pool_2 | | 1613 | DAPK3 | S | 0,67 | 0,56 | 0,39 | 0,30 | 0,61 | 0,34 | 0,08 | 0,07 | 0,56 | 0,56 | 0,64 | 0,08 | 0,13 | -0,61 |
| 461 | Dha_Kin_pool_2 | | 4214 | MAP3K1 | S | 0,80 | 0,79 | 0,62 | 0,49 | 0,79 | 0,56 | 0,01 | 0,09 | 0,70 | 0,70 | 0,64 | 0,06 | 0,13 | 0,42 |
| 462 | Dha_Kin_pool_2 | | 7535 | ZAP70 | S | 1,14 | 0,85 | 0,79 | 0,62 | 1,00 | 0,71 | 0,21 | 0,12 | 0,71 | 0,71 | 0,64 | 0,07 | 0,13 | 0,50 |
| 463 | Dha_Kin_pool_2 | | 26576 | STK23 | S | 0,84 | 0,75 | 0,56 | 0,49 | 0,79 | 0,53 | 0,06 | 0,05 | 0,66 | 0,66 | 0,64 | 0,02 | 0,13 | 0,13 |
| 464 | Dha_Kin_pool_2 | | 5754 | PTK7 | S | 0,74 | 0,73 | 0,61 | 0,48 | 0,74 | 0,55 | 0,00 | 0,09 | 0,74 | 0,74 | 0,64 | 0,10 | 0,13 | 0,72 |
| 465 | Dha_Kin_pool_2 | | 83694 | RPS6KL1 | S | 0,42 | 0,41 | 0,15 | 0,10 | 0,42 | 0,12 | 0,01 | 0,04 | 0,29 | 0,29 | 0,64 | 0,35 | 0,13 | -2,61 |
| 466 | Dha_Kin_pool_2 | | 8396 | PIP5K2B | S | 0,90 | 0,79 | 0,54 | 0,50 | 0,84 | 0,52 | 0,08 | 0,03 | 0,61 | 0,61 | 0,64 | 0,03 | 0,13 | -0,22 |
| 467 | Dha_Kin_pool_2 | | 6789 | STK4 | S | 0,69 | 0,49 | 0,67 | 0,48 | 0,59 | 0,58 | 0,14 | 0,13 | 0,98 | 0,98 | 0,64 | 0,34 | 0,13 | 2,51 |
| 468 | Dha_Kin_pool_2 | | 9223 | BAIAP1 | S | 0,67 | 0,70 | 0,67 | 0,51 | 0,68 | 0,59 | 0,02 | 0,11 | 0,87 | 0,87 | 0,64 | 0,23 | 0,13 | 1,69 |
| 469 | Dha_Kin_pool_2 | | 9748 | SLK | S | 0,52 | 0,50 | 0,23 | 0,25 | 0,51 | 0,24 | 0,02 | 0,01 | 0,48 | 0,48 | 0,64 | 0,17 | 0,13 | -1,23 |
| 470 | Dha_Kin_pool_2 | | 23396 | PIP5K1C | S | 0,73 | 0,54 | 0,59 | 0,50 | 0,63 | 0,54 | 0,14 | 0,06 | 0,86 | 0,86 | 0,64 | 0,21 | 0,13 | 1,58 |
| 471 | Dha_Kin_pool_2 | | 5211 | PFKL | S | 0,55 | 0,44 | 0,36 | 0,28 | 0,49 | 0,32 | 0,08 | 0,06 | 0,64 | 0,64 | 0,64 | 0,00 | 0,13 | -0,01 |
| 472 | Dha_Kin_pool_2 | | 5105 | PCK1 | S | 0,49 | 0,47 | 0,55 | 0,37 | 0,48 | 0,46 | 0,01 | 0,13 | 0,96 | 0,96 | 0,64 | 0,32 | 0,13 | 2,36 |
| 473 | Dha_Kin_pool_2 | | 1025 | CDK9 | S | 0,31 | 0,33 | 0,28 | 0,17 | 0,32 | 0,23 | 0,01 | 0,08 | 0,71 | 0,71 | 0,64 | 0,06 | 0,13 | 0,47 |
| 474 | Dha_Kin_pool_2 | | 2264 | FGFR4 | S | 0,71 | 0,68 | 0,44 | 0,47 | 0,70 | 0,46 | 0,02 | 0,02 | 0,65 | 0,65 | 0,64 | 0,01 | 0,13 | 0,09 |
| 475 | Dha_Kin_pool_2 | | 27330 | RPS6KA6 | S | 0,93 | 0,89 | 0,76 | 0,60 | 0,91 | 0,68 | 0,03 | 0,11 | 0,74 | 0,74 | 0,64 | 0,10 | 0,13 | 0,74 |
| 476 | Dha_Kin_pool_2 | | 6732 | SRPK1 | S | 0,95 | 0,99 | 0,65 | 0,60 | 0,97 | 0,63 | 0,02 | 0,03 | 0,65 | 0,65 | 0,64 | 0,00 | 0,13 | 0,02 |
| 477 | Dha_Kin_pool_2 | | 5566 | PRKACA | S | 0,94 | 0,87 | 0,56 | 0,42 | 0,90 | 0,49 | 0,05 | 0,10 | 0,54 | 0,54 | 0,64 | 0,10 | 0,13 | -0,73 |
| 478 | Dha_Kin_pool_2 | | 2185 | PTK2B | S | 0,70 | 0,65 | 0,55 | 0,46 | 0,68 | 0,50 | 0,03 | 0,06 | 0,74 | 0,74 | 0,64 | 0,10 | 0,13 | 0,73 |
| 479 | Dha_Kin_pool_2 | E | | E | D | 1,01 | 1,10 | 1,06 | 1,11 | 1,06 | 1,09 | 0,07 | 0,04 | 1,03 | | 0,64 | | 0,13 | 2,85 |
| 480 | Dha_Kin_pool_2 | E | | E | D | 1,00 | 0,96 | 0,97 | 1,14 | 0,98 | 1,06 | 0,02 | 0,13 | 1,07 | | 0,64 | | 0,13 | 3,19 |
| 481 | Dha_Kin_pool_2 | | 10461 | MERTK | S | 0,93 | 0,95 | 0,56 | 0,48 | 0,94 | 0,52 | 0,01 | 0,05 | 0,55 | 0,55 | 0,64 | 0,09 | 0,13 | -0,69 |
| 482 | Dha_Kin_pool_2 | | 64781 | CERK | S | 0,98 | 0,88 | 0,77 | 0,63 | 0,93 | 0,70 | 0,08 | 0,09 | 0,75 | 0,75 | 0,64 | 0,11 | 0,13 | 0,80 |
| 483 | Dha_Kin_pool_2 | | 64710 | NUCKS | S | 0,98 | 0,67 | 0,45 | 0,45 | 0,83 | 0,45 | 0,22 | 0,00 | 0,54 | 0,54 | 0,64 | 0,10 | 0,13 | -0,76 |
| 484 | Dha_Kin_pool_2 | | 11113 | CIT | S | 0,60 | 0,71 | 0,34 | 0,37 | 0,66 | 0,35 | 0,08 | 0,02 | 0,54 | 0,54 | 0,64 | 0,11 | 0,13 | -0,79 |
| 485 | Dha_Kin_pool_2 | | 344387 | CDKL4 | S | 1,07 | 1,13 | 0,50 | 0,46 | 1,10 | 0,48 | 0,04 | 0,03 | 0,43 | 0,43 | 0,64 | 0,21 | 0,13 | -1,55 |
| 486 | Dha_Kin_pool_2 | | 79834 | KIAA2002 | S | 0,82 | 0,75 | 0,66 | 0,57 | 0,78 | 0,61 | 0,04 | 0,07 | 0,78 | 0,78 | 0,64 | 0,14 | 0,13 | 1,03 |
| 487 | Dha_Kin_pool_2 | | 1399 | CRKL | S | 0,69 | 0,82 | 0,59 | 0,36 | 0,75 | 0,48 | 0,10 | 0,17 | 0,63 | 0,63 | 0,64 | 0,01 | 0,13 | -0,09 |
| 488 | Dha_Kin_pool_2 | | 9252 | RPS6KA5 | S | 0,79 | 0,83 | 0,53 | 0,66 | 0,81 | 0,59 | 0,03 | 0,10 | 0,73 | 0,73 | 0,64 | 0,09 | 0,13 | 0,67 |
| 489 | Dha_Kin_pool_2 | | 282974 | STK32C | S | 0,57 | 0,63 | 0,50 | 0,25 | 0,60 | 0,38 | 0,04 | 0,17 | 0,63 | 0,63 | 0,64 | 0,01 | 0,13 | -0,08 |
| 490 | Dha_Kin_pool_2 | | 25865 | PRKD2 | S | 0,72 | 0,69 | 0,94 | 0,54 | 0,71 | 0,74 | 0,02 | 0,28 | 1,04 | 1,04 | 0,64 | 0,40 | 0,13 | 2,96 |
| 491 | Dha_Kin_pool_2 | | 7084 | TK2 | S | 0,73 | 0,82 | 0,66 | 0,43 | 0,77 | 0,55 | 0,07 | 0,16 | 0,71 | 0,71 | 0,64 | 0,06 | 0,13 | 0,47 |
| 492 | Dha_Kin_pool_2 | | 1457 | CSNK2A1 | S | 0,41 | 0,45 | 0,30 | 0,16 | 0,43 | 0,23 | 0,03 | 0,10 | 0,54 | 0,54 | 0,64 | 0,11 | 0,13 | -0,80 |
| 493 | Dha_Kin_pool_2 | | 5871 | MAP4K2 | S | 0,73 | 0,81 | 0,63 | 0,39 | 0,77 | 0,51 | 0,06 | 0,17 | 0,66 | 0,66 | 0,64 | 0,02 | 0,13 | 0,13 |
| 494 | Dha_Kin_pool_2 | | 9943 | OSR1 | S | 0,89 | 0,61 | 0,47 | 0,36 | 0,75 | 0,41 | 0,20 | 0,08 | 0,55 | 0,55 | 0,64 | 0,09 | 0,13 | -0,69 |

| | | | | | | | | | | | | | | | | | | | |
|---|---|---|---|---|---|---|---|---|---|---|---|---|---|---|---|---|---|---|---|
| 495 | Dha_Kin_pool_2 | | 5891 | RAGE | S | 0,78 | 0,79 | 0,41 | 0,33 | 0,78 | 0,37 | 0,00 | 0,06 | 0,47 | 0,47 | 0,64 | 0,17 | 0,13 | -1,28 |
| 496 | Dha_Kin_pool_2 | | 659 | BMPR2 | S | 0,59 | 0,62 | 0,29 | 0,19 | 0,61 | 0,24 | 0,02 | 0,08 | 0,40 | 0,40 | 0,64 | 0,25 | 0,13 | -1,82 |
| 497 | Dha_Kin_pool_2 | | 340371 | NRBP2 | S | 0,40 | 0,32 | 0,45 | 0,24 | 0,36 | 0,35 | 0,06 | 0,14 | 0,96 | 0,96 | 0,64 | 0,31 | 0,13 | 2,31 |
| 498 | Dha_Kin_pool_2 | | 6416 | MAP2K4 | S | 0,91 | 0,93 | 0,68 | 0,65 | 0,92 | 0,66 | 0,02 | 0,02 | 0,72 | 0,72 | 0,64 | 0,08 | 0,13 | 0,57 |
| 499 | Dha_Kin_pool_2 | | 65220 | FLJ13052 | S | 0,81 | 0,79 | 0,47 | 0,53 | 0,80 | 0,50 | 0,01 | 0,04 | 0,62 | 0,62 | 0,64 | 0,02 | 0,13 | -0,14 |
| 500 | Dha_Kin_pool_2 | | 11011 | TLK2 | S | 1,08 | 0,99 | 0,78 | 0,69 | 1,04 | 0,74 | 0,06 | 0,06 | 0,71 | 0,71 | 0,64 | 0,07 | 0,13 | 0,50 |
| 501 | Dha_Kin_pool_2 | | 9414 | TJP2 | S | 0,62 | 0,49 | 0,45 | 0,26 | 0,55 | 0,36 | 0,09 | 0,14 | 0,64 | 0,64 | 0,64 | 0,00 | 0,13 | 0,00 |
| 502 | Dha_Kin_pool_2 | | 203054 | ADCK5 | S | 0,91 | 0,73 | 0,76 | 0,54 | 0,82 | 0,65 | 0,12 | 0,16 | 0,79 | 0,79 | 0,64 | 0,14 | 0,13 | 1,07 |
| 503 | Dha_Kin_pool_2 | E | | E | D | 1,02 | 1,18 | 0,94 | 0,95 | 1,10 | 0,95 | 0,11 | 0,01 | 0,86 | | 0,64 | | 0,13 | 1,63 |
| 504 | Dha_Kin_pool_2 | E | | E | D | 1,00 | 1,24 | 1,05 | 1,05 | 1,12 | 1,05 | 0,18 | 0,00 | 0,93 | | 0,64 | | 0,13 | 2,15 |
| 505 | Dha_Kin_pool_2 | | 5600 | MAPK11 | S | 0,95 | 0,83 | 0,43 | 0,49 | 0,89 | 0,46 | 0,09 | 0,04 | 0,51 | 0,51 | 0,64 | 0,13 | 0,13 | -0,96 |
| 506 | Dha_Kin_pool_2 | | 5163 | PDK1 | S | 0,78 | 0,77 | 0,45 | 0,38 | 0,77 | 0,42 | 0,01 | 0,04 | 0,54 | 0,54 | 0,64 | 0,11 | 0,13 | -0,79 |
| 507 | Dha_Kin_pool_2 | | 253430 | IPMK | S | 0,71 | 0,70 | 0,48 | 0,28 | 0,71 | 0,38 | 0,00 | 0,15 | 0,54 | 0,54 | 0,64 | 0,10 | 0,13 | -0,77 |
| 508 | Dha_Kin_pool_2 | | 9833 | MELK | S | 0,93 | 0,97 | 0,53 | 0,45 | 0,95 | 0,49 | 0,03 | 0,06 | 0,52 | 0,52 | 0,64 | 0,12 | 0,13 | -0,92 |
| 509 | Dha_Kin_pool_2 | | 5583 | PRKCH | S | 0,99 | 0,88 | 0,67 | 0,38 | 0,94 | 0,53 | 0,08 | 0,20 | 0,56 | 0,56 | 0,64 | 0,08 | 0,13 | -0,61 |
| 510 | Dha_Kin_pool_2 | | 5063 | PAK3 | S | 0,93 | 0,64 | 0,49 | 0,29 | 0,79 | 0,39 | 0,21 | 0,14 | 0,50 | 0,50 | 0,64 | 0,14 | 0,13 | -1,06 |
| 511 | Dha_Kin_pool_2 | | 3645 | INSRR | S | 0,81 | 0,88 | 0,61 | 0,35 | 0,84 | 0,48 | 0,05 | 0,19 | 0,57 | 0,57 | 0,64 | 0,08 | 0,13 | -0,56 |
| 512 | Dha_Kin_pool_2 | | 200576 | PIP5K3 | S | 0,69 | 0,67 | 0,54 | 0,32 | 0,68 | 0,43 | 0,01 | 0,15 | 0,63 | 0,63 | 0,64 | 0,02 | 0,13 | -0,12 |
| 513 | Dha_Kin_pool_2 | | 657 | BMPR1A | S | 0,72 | 0,70 | 0,60 | 0,44 | 0,71 | 0,52 | 0,01 | 0,11 | 0,73 | 0,73 | 0,64 | 0,09 | 0,13 | 0,66 |
| 514 | Dha_Kin_pool_2 | | 5232 | PGK2 | S | 0,54 | 0,53 | 0,43 | 0,27 | 0,54 | 0,35 | 0,01 | 0,11 | 0,66 | 0,66 | 0,64 | 0,01 | 0,13 | 0,10 |
| 515 | Dha_Kin_pool_2 | | 701 | BUB1B | S | 0,40 | 0,43 | 0,25 | 0,13 | 0,41 | 0,19 | 0,02 | 0,09 | 0,47 | 0,47 | 0,64 | 0,18 | 0,13 | -1,32 |
| 516 | Dha_Kin_pool_2 | | 5607 | MAP2K5 | S | 0,73 | 0,65 | 0,61 | 0,46 | 0,69 | 0,53 | 0,06 | 0,10 | 0,78 | 0,78 | 0,64 | 0,13 | 0,13 | 0,98 |
| 517 | Dha_Kin_pool_2 | | 80216 | LAK | S | 0,65 | 0,60 | 0,49 | 0,35 | 0,63 | 0,42 | 0,03 | 0,10 | 0,67 | 0,67 | 0,64 | 0,03 | 0,13 | 0,22 |
| 518 | Dha_Kin_pool_2 | | 1022 | CDK7 | S | 0,79 | 0,68 | 0,53 | 0,45 | 0,74 | 0,49 | 0,08 | 0,06 | 0,67 | 0,67 | 0,64 | 0,02 | 0,13 | 0,18 |
| 519 | Dha_Kin_pool_2 | | 79705 | LRRK1 | S | 0,83 | 0,65 | 0,74 | 0,50 | 0,74 | 0,62 | 0,13 | 0,17 | 0,84 | 0,84 | 0,64 | 0,19 | 0,13 | 1,43 |
| 520 | Dha_Kin_pool_2 | | 124923 | FLJ25006 | S | 0,54 | 0,68 | 0,51 | 0,33 | 0,61 | 0,42 | 0,10 | 0,13 | 0,69 | 0,69 | 0,64 | 0,05 | 0,13 | 0,36 |
| 521 | Dha_Kin_pool_2 | | 1432 | MAPK14 | S | 0,90 | 0,75 | 0,77 | 0,54 | 0,83 | 0,65 | 0,10 | 0,17 | 0,79 | 0,79 | 0,64 | 0,15 | 0,13 | 1,10 |
| 522 | Dha_Kin_pool_2 | | 29110 | TBK1 | S | 0,75 | 0,69 | 0,58 | 0,49 | 0,72 | 0,54 | 0,04 | 0,06 | 0,75 | 0,75 | 0,64 | 0,10 | 0,13 | 0,76 |
| 523 | Dha_Kin_pool_2 | | 10595 | ERN2 | S | 0,90 | 0,85 | 0,71 | 0,59 | 0,88 | 0,65 | 0,03 | 0,09 | 0,75 | 0,75 | 0,64 | 0,10 | 0,13 | 0,76 |
| 524 | Dha_Kin_pool_2 | | 7049 | TGFBR3 | S | 0,80 | 0,79 | 0,78 | 0,58 | 0,80 | 0,68 | 0,01 | 0,14 | 0,86 | 0,86 | 0,64 | 0,21 | 0,13 | 1,58 |
| 525 | Dha_Kin_pool_2 | | 51727 | UMP-CMPK | S | 0,47 | 0,42 | 0,41 | 0,27 | 0,44 | 0,34 | 0,04 | 0,10 | 0,76 | 0,76 | 0,64 | 0,12 | 0,13 | 0,87 |
| 526 | Dha_Kin_pool_2 | | 1119 | CHKA | S | 0,71 | 0,87 | 0,43 | 0,41 | 0,79 | 0,42 | 0,11 | 0,02 | 0,53 | 0,53 | 0,64 | 0,11 | 0,13 | -0,82 |
| 527 | Dha_Kin_pool_2 | E | | E | D | 1,07 | 1,11 | 1,13 | 1,01 | 1,09 | 1,07 | 0,03 | 0,09 | 0,98 | | 0,64 | | 0,13 | 2,52 |
| 528 | Dha_Kin_pool_2 | E | | E | D | 0,97 | 0,93 | 0,91 | 1,08 | 0,95 | 1,00 | 0,03 | 0,12 | 1,05 | | 0,64 | | 0,13 | 3,03 |
| 529 | Dha_Kin_pool_2 | | 65266 | WNK4 | S | 0,89 | 0,81 | 0,51 | 0,51 | 0,85 | 0,51 | 0,05 | 0,00 | 0,60 | 0,60 | 0,64 | 0,04 | 0,13 | -0,30 |
| 530 | Dha_Kin_pool_2 | | 6195 | RPS6KA1 | S | 0,99 | 0,86 | 0,72 | 0,46 | 0,92 | 0,59 | 0,10 | 0,18 | 0,64 | 0,64 | 0,64 | 0,00 | 0,13 | 0,00 |
| 531 | Dha_Kin_pool_2 | | 117283 | IHPK3 | S | 0,45 | 0,42 | 0,19 | 0,12 | 0,43 | 0,15 | 0,02 | 0,05 | 0,35 | 0,35 | 0,64 | 0,30 | 0,13 | -2,19 |
| 532 | Dha_Kin_pool_2 | | 660 | BMX | S | 1,12 | 0,85 | 0,57 | 0,61 | 0,98 | 0,59 | 0,19 | 0,03 | 0,60 | 0,60 | 0,64 | 0,04 | 0,13 | -0,29 |
| 533 | Dha_Kin_pool_2 | | 57396 | CLK4 | S | 0,87 | 0,68 | 0,61 | 0,34 | 0,78 | 0,48 | 0,14 | 0,19 | 0,62 | 0,62 | 0,64 | 0,03 | 0,13 | -0,21 |
| 534 | Dha_Kin_pool_2 | | 5062 | PAK2 | S | 0,71 | 0,55 | 0,51 | 0,31 | 0,63 | 0,41 | 0,11 | 0,14 | 0,65 | 0,65 | 0,64 | 0,00 | 0,13 | 0,02 |
| 535 | Dha_Kin_pool_2 | | 1452 | CSNK1A1 | S | 0,59 | 0,59 | 0,56 | 0,31 | 0,59 | 0,43 | 0,00 | 0,18 | 0,72 | 0,72 | 0,64 | 0,08 | 0,13 | 0,60 |
| 536 | Dha_Kin_pool_2 | | 4831 | NME2 | S | 0,50 | 0,50 | 0,50 | 0,44 | 0,50 | 0,47 | 0,00 | 0,04 | 0,94 | 0,94 | 0,64 | 0,29 | 0,13 | 2,17 |
| 537 | Dha_Kin_pool_2 | | 4117 | MAK | S | 0,50 | 0,52 | 0,34 | 0,22 | 0,51 | 0,28 | 0,01 | 0,08 | 0,56 | 0,56 | 0,64 | 0,09 | 0,13 | -0,64 |
| 538 | Dha_Kin_pool_2 | | 5293 | PIK3CD | S | 0,56 | 0,56 | 0,41 | 0,28 | 0,56 | 0,35 | 0,00 | 0,09 | 0,62 | 0,62 | 0,64 | 0,03 | 0,13 | -0,21 |
| 539 | Dha_Kin_pool_2 | | 9064 | MAP3K6 | S | 0,48 | 0,60 | 0,31 | 0,30 | 0,54 | 0,31 | 0,08 | 0,01 | 0,57 | 0,57 | 0,64 | 0,07 | 0,13 | -0,53 |
| 540 | Dha_Kin_pool_2 | | 440275 | EIF2AK4 | S | 0,58 | 0,53 | 0,32 | 0,24 | 0,56 | 0,28 | 0,03 | 0,06 | 0,50 | 0,50 | 0,64 | 0,14 | 0,13 | -1,06 |
| 541 | Dha_Kin_pool_2 | | 10201 | NME6 | S | 0,79 | 0,67 | 0,56 | 0,48 | 0,73 | 0,52 | 0,09 | 0,06 | 0,71 | 0,71 | 0,64 | 0,06 | 0,13 | 0,48 |
| 542 | Dha_Kin_pool_2 | | 1859 | DYRK1A | S | 0,72 | 0,78 | 0,37 | 0,26 | 0,75 | 0,31 | 0,04 | 0,08 | 0,41 | 0,41 | 0,64 | 0,23 | 0,13 | -1,69 |
| 543 | Dha_Kin_pool_2 | | 814 | CAMK4 | S | 0,87 | 0,93 | 0,69 | 0,53 | 0,90 | 0,61 | 0,04 | 0,12 | 0,68 | 0,68 | 0,64 | 0,03 | 0,13 | 0,25 |
| 544 | Dha_Kin_pool_2 | | 51347 | JIK | S | 0,73 | 0,79 | 0,55 | 0,40 | 0,76 | 0,47 | 0,04 | 0,11 | 0,63 | 0,63 | 0,64 | 0,02 | 0,13 | -0,13 |
| 545 | Dha_Kin_pool_2 | | 5313 | PKLR | S | 0,79 | 0,95 | 0,73 | 0,51 | 0,87 | 0,62 | 0,11 | 0,16 | 0,71 | 0,71 | 0,64 | 0,07 | 0,13 | 0,48 |
| 546 | Dha_Kin_pool_2 | | 57144 | PAK7 | S | 0,45 | 0,52 | 0,38 | 0,26 | 0,48 | 0,32 | 0,05 | 0,08 | 0,67 | 0,67 | 0,64 | 0,03 | 0,13 | 0,19 |
| 547 | Dha_Kin_pool_2 | | 7443 | VRK1 | S | 0,84 | 0,77 | 0,85 | 0,60 | 0,80 | 0,73 | 0,05 | 0,18 | 0,91 | 0,91 | 0,64 | 0,26 | 0,13 | 1,95 |
| 548 | Dha_Kin_pool_2 | | 285220 | EPHA6 | S | 1,10 | 0,96 | 0,90 | 0,73 | 1,03 | 0,81 | 0,10 | 0,12 | 0,79 | 0,79 | 0,64 | 0,14 | 0,13 | 1,05 |
| 549 | Dha_Kin_pool_2 | | 139728 | PNCK | S | 0,84 | 0,80 | 0,58 | 0,39 | 0,82 | 0,49 | 0,03 | 0,13 | 0,59 | 0,59 | 0,64 | 0,05 | 0,13 | -0,39 |

| | | | | | | | | | | | | | | | | | | | |
|---|---|---|---|---|---|---|---|---|---|---|---|---|---|---|---|---|---|---|---|
| 550 | Dha_Kin_pool_2 | | 51231 | VRK3 | S | 1,16 | 0,89 | 0,90 | 0,62 | 1,03 | 0,76 | 0,19 | 0,20 | 0,74 | 0,74 | 0,64 | 0,10 | 0,13 | 0,73 |
| 551 | Dha_Kin_pool_2 | E | | E | D | 1,09 | 1,08 | 1,20 | 1,01 | 1,09 | 1,10 | 0,01 | 0,13 | 1,01 | | 0,64 | | 0,13 | 2,75 |
| 552 | Dha_Kin_pool_2 | E | | E | D | 1,12 | 1,15 | 1,12 | 1,00 | 1,14 | 1,06 | 0,02 | 0,08 | 0,93 | | 0,64 | | 0,13 | 2,14 |
| 553 | Dha_Kin_pool_2 | | 5156 | PDGFRA | S | 0,84 | 0,85 | 0,42 | 0,27 | 0,84 | 0,35 | 0,01 | 0,10 | 0,41 | 0,41 | 0,64 | 0,23 | 0,13 | -1,73 |
| 554 | Dha_Kin_pool_2 | | 780 | DDR1 | S | 0,92 | 0,62 | 0,52 | 0,40 | 0,77 | 0,46 | 0,21 | 0,09 | 0,59 | 0,59 | 0,64 | 0,05 | 0,13 | -0,37 |
| 555 | Dha_Kin_pool_2 | | 5286 | PIK3C2A | S | 0,72 | 0,62 | 0,49 | 0,37 | 0,67 | 0,43 | 0,08 | 0,08 | 0,64 | 0,64 | 0,64 | 0,01 | 0,13 | -0,04 |
| 556 | Dha_Kin_pool_2 | | 202374 | STK32A | S | 0,87 | 0,70 | 0,52 | 0,41 | 0,78 | 0,47 | 0,12 | 0,07 | 0,60 | 0,60 | 0,64 | 0,05 | 0,13 | -0,35 |
| 557 | Dha_Kin_pool_2 | | 55872 | TOPK | S | 0,58 | 0,39 | 0,38 | 0,25 | 0,49 | 0,32 | 0,13 | 0,09 | 0,65 | 0,65 | 0,64 | 0,01 | 0,13 | 0,04 |
| 558 | Dha_Kin_pool_2 | | 51776 | ZAK | S | 0,89 | 0,76 | 0,54 | 0,45 | 0,82 | 0,50 | 0,09 | 0,06 | 0,60 | 0,60 | 0,64 | 0,04 | 0,13 | -0,29 |
| 559 | Dha_Kin_pool_2 | | 5166 | PDK4 | S | 0,59 | 0,71 | 0,48 | 0,33 | 0,65 | 0,41 | 0,08 | 0,11 | 0,63 | 0,63 | 0,64 | 0,01 | 0,13 | -0,11 |
| 560 | Dha_Kin_pool_2 | | 1147 | CHUK | S | 0,80 | 0,77 | 0,58 | 0,47 | 0,79 | 0,52 | 0,02 | 0,08 | 0,66 | 0,66 | 0,64 | 0,02 | 0,13 | 0,16 |
| 561 | Dha_Kin_pool_2 | | 27010 | TPK1 | S | 0,68 | 0,75 | 0,50 | 0,23 | 0,71 | 0,37 | 0,05 | 0,19 | 0,52 | 0,52 | 0,64 | 0,13 | 0,13 | -0,93 |
| 562 | Dha_Kin_pool_2 | | 283629 | C14ORF20 | S | 0,91 | 0,74 | 0,53 | 0,48 | 0,83 | 0,50 | 0,12 | 0,03 | 0,61 | 0,61 | 0,64 | 0,04 | 0,13 | -0,27 |
| 563 | Dha_Kin_pool_2 | | 415116 | PIM3 | S | 0,86 | 0,88 | 0,69 | 0,36 | 0,87 | 0,52 | 0,01 | 0,23 | 0,60 | 0,60 | 0,64 | 0,04 | 0,13 | -0,32 |
| 564 | Dha_Kin_pool_2 | | 114783 | LMTK3 | S | 0,69 | 0,65 | 0,51 | 0,34 | 0,67 | 0,42 | 0,02 | 0,12 | 0,63 | 0,63 | 0,64 | 0,01 | 0,13 | -0,08 |
| 565 | Dha_Kin_pool_2 | | 11200 | CHEK2 | S | 0,64 | 0,71 | 0,40 | 0,29 | 0,68 | 0,34 | 0,05 | 0,07 | 0,51 | 0,51 | 0,64 | 0,14 | 0,13 | -1,02 |
| 566 | Dha_Kin_pool_2 | | 3716 | JAK1 | S | 0,56 | 0,61 | 0,23 | 0,19 | 0,59 | 0,21 | 0,03 | 0,02 | 0,36 | 0,36 | 0,64 | 0,29 | 0,13 | -2,12 |
| 567 | Dha_Kin_pool_2 | | 9451 | EIF2AK3 | S | 0,69 | 0,78 | 0,54 | 0,34 | 0,73 | 0,44 | 0,06 | 0,14 | 0,60 | 0,60 | 0,64 | 0,04 | 0,13 | -0,30 |
| 568 | Dha_Kin_pool_2 | | 5289 | PIK3C3 | S | 0,94 | 0,75 | 0,54 | 0,62 | 0,84 | 0,58 | 0,13 | 0,06 | 0,69 | 0,69 | 0,64 | 0,05 | 0,13 | 0,34 |
| 569 | Dha_Kin_pool_2 | | 9475 | ROCK2 | S | 0,68 | 0,60 | 0,24 | 0,26 | 0,64 | 0,25 | 0,05 | 0,02 | 0,39 | 0,39 | 0,64 | 0,25 | 0,13 | -1,86 |
| 570 | Dha_Kin_pool_2 | | 5287 | PIK3C2B | S | 0,69 | 0,61 | 0,70 | 0,41 | 0,65 | 0,56 | 0,05 | 0,21 | 0,86 | 0,86 | 0,64 | 0,21 | 0,13 | 1,57 |
| 571 | Dha_Kin_pool_2 | | 6446 | SGK | S | 0,82 | 0,70 | 0,81 | 0,40 | 0,76 | 0,60 | 0,09 | 0,29 | 0,79 | 0,79 | 0,64 | 0,15 | 0,13 | 1,12 |
| 572 | Dha_Kin_pool_2 | | 1198 | CLK3 | S | 1,13 | 0,99 | 0,86 | 0,54 | 1,06 | 0,70 | 0,10 | 0,23 | 0,66 | 0,66 | 0,64 | 0,02 | 0,13 | 0,15 |
| 573 | Dha_Kin_pool_2 | | 54899 | PXK | S | 0,94 | 0,84 | 0,86 | 0,68 | 0,89 | 0,77 | 0,07 | 0,12 | 0,86 | 0,86 | 0,64 | 0,22 | 0,13 | 1,64 |
| 574 | Dha_Kin_pool_2 | | 30811 | HUNK | S | 0,89 | 0,90 | 0,52 | 0,57 | 0,90 | 0,55 | 0,00 | 0,03 | 0,61 | 0,61 | 0,64 | 0,04 | 0,13 | -0,26 |
| 575 | Dha_Kin_pool_2 | E | | E | D | 1,18 | 0,84 | 1,21 | 0,89 | 1,01 | 1,05 | 0,24 | 0,22 | 1,04 | | 0,64 | | 0,13 | 2,93 |
| 576 | Dha_Kin_pool_2 | E | | E | D | 0,90 | 0,86 | 0,93 | 0,77 | 0,88 | 0,85 | 0,03 | 0,11 | 0,96 | | 0,64 | | 0,13 | 2,38 |
| 577 | Dha_Kin_pool_2 | | 11184 | MAP4K1 | S | 0,50 | 0,55 | 0,35 | 0,22 | 0,52 | 0,29 | 0,04 | 0,09 | 0,55 | 0,55 | 0,64 | 0,09 | 0,13 | -0,69 |
| 578 | Dha_Kin_pool_2 | | 5128 | PCTK2 | S | 1,01 | 0,71 | 0,69 | 0,50 | 0,86 | 0,59 | 0,21 | 0,13 | 0,69 | 0,69 | 0,64 | 0,05 | 0,13 | 0,35 |
| 579 | Dha_Kin_pool_2 | | 3643 | INSR | S | 0,44 | 0,37 | 0,32 | 0,17 | 0,41 | 0,25 | 0,05 | 0,11 | 0,61 | 0,61 | 0,64 | 0,03 | 0,13 | -0,26 |
| 580 | Dha_Kin_pool_2 | | 147746 | HIPK4 | S | 0,77 | 0,60 | 0,40 | 0,39 | 0,69 | 0,39 | 0,12 | 0,00 | 0,57 | 0,57 | 0,64 | 0,07 | 0,13 | -0,51 |
| 581 | Dha_Kin_pool_2 | | 23235 | SIK2 | S | 0,79 | 0,69 | 0,51 | 0,52 | 0,74 | 0,52 | 0,07 | 0,00 | 0,70 | 0,70 | 0,64 | 0,06 | 0,13 | 0,41 |
| 582 | Dha_Kin_pool_2 | | 4233 | MET | S | 0,55 | 0,55 | 0,44 | 0,26 | 0,55 | 0,35 | 0,00 | 0,12 | 0,63 | 0,63 | 0,64 | 0,01 | 0,13 | -0,07 |
| 583 | Dha_Kin_pool_2 | | 2534 | FYN | S | 0,77 | 0,72 | 0,63 | 0,47 | 0,74 | 0,55 | 0,03 | 0,12 | 0,74 | 0,74 | 0,64 | 0,10 | 0,13 | 0,71 |
| 584 | Dha_Kin_pool_2 | | 25989 | DKFZP434C131 | S | 0,76 | 0,75 | 0,64 | 0,42 | 0,75 | 0,53 | 0,01 | 0,15 | 0,70 | 0,70 | 0,64 | 0,06 | 0,13 | 0,42 |
| 585 | Dha_Kin_pool_2 | | 22983 | SAST | S | 0,23 | 0,20 | 0,12 | 0,05 | 0,21 | 0,09 | 0,02 | 0,05 | 0,40 | 0,40 | 0,64 | 0,24 | 0,13 | -1,79 |
| 586 | Dha_Kin_pool_2 | | 4751 | NEK2 | S | 0,89 | 0,72 | 0,47 | 0,49 | 0,81 | 0,48 | 0,12 | 0,01 | 0,59 | 0,59 | 0,64 | 0,05 | 0,13 | -0,37 |
| 587 | Dha_Kin_pool_2 | | 8550 | MAPKAPK5 | S | 0,51 | 0,37 | 0,42 | 0,25 | 0,44 | 0,34 | 0,10 | 0,12 | 0,77 | 0,77 | 0,64 | 0,12 | 0,13 | 0,91 |
| 588 | Dha_Kin_pool_2 | | 203447 | NRK | S | 0,83 | 0,78 | 0,44 | 0,43 | 0,80 | 0,44 | 0,03 | 0,01 | 0,54 | 0,54 | 0,64 | 0,10 | 0,13 | -0,75 |
| 589 | Dha_Kin_pool_2 | | 57787 | MARK4 | S | 0,16 | 0,20 | 0,19 | 0,12 | 0,18 | 0,15 | 0,03 | 0,04 | 0,87 | 0,87 | 0,64 | 0,22 | 0,13 | 1,66 |
| 590 | Dha_Kin_pool_2 | | 2047 | EPHB1 | S | 0,87 | 0,75 | 0,54 | 0,44 | 0,81 | 0,49 | 0,08 | 0,07 | 0,61 | 0,61 | 0,64 | 0,04 | 0,13 | -0,27 |
| 591 | Dha_Kin_pool_2 | | 260425 | MAGI-3 | S | 1,00 | 0,80 | 0,62 | 0,45 | 0,90 | 0,53 | 0,14 | 0,12 | 0,59 | 0,59 | 0,64 | 0,05 | 0,13 | -0,39 |
| 592 | Dha_Kin_pool_2 | | 8517 | IKBKG | S | 0,81 | 0,84 | 0,45 | 0,40 | 0,83 | 0,42 | 0,02 | 0,03 | 0,51 | 0,51 | 0,64 | 0,13 | 0,13 | -0,97 |
| 593 | Dha_Kin_pool_2 | | 55728 | N4BP2 | S | 0,90 | 0,88 | 0,54 | 0,44 | 0,89 | 0,49 | 0,01 | 0,07 | 0,55 | 0,55 | 0,64 | 0,09 | 0,13 | -0,69 |
| 594 | Dha_Kin_pool_2 | | 4750 | NEK1 | S | 0,97 | 0,82 | 0,63 | 0,55 | 0,89 | 0,59 | 0,10 | 0,06 | 0,66 | 0,66 | 0,64 | 0,02 | 0,13 | 0,16 |
| 595 | Dha_Kin_pool_2 | | 9942 | XYLB | S | 0,63 | 0,58 | 0,34 | 0,16 | 0,61 | 0,25 | 0,04 | 0,12 | 0,41 | 0,41 | 0,64 | 0,23 | 0,13 | -1,70 |
| 596 | Dha_Kin_pool_2 | | 51765 | RP6-213H19.1 | S | 0,88 | 0,85 | 0,64 | 0,61 | 0,87 | 0,63 | 0,03 | 0,02 | 0,72 | 0,72 | 0,64 | 0,08 | 0,13 | 0,61 |
| 597 | Dha_Kin_pool_2 | | 23049 | SMG1 | S | 1,04 | 0,83 | 0,83 | 0,54 | 0,94 | 0,68 | 0,15 | 0,20 | 0,73 | 0,73 | 0,64 | 0,09 | 0,13 | 0,65 |
| 598 | Dha_Kin_pool_2 | | 51447 | IHPK2 | S | 0,75 | 0,61 | 0,41 | 0,35 | 0,68 | 0,38 | 0,10 | 0,04 | 0,55 | 0,55 | 0,64 | 0,09 | 0,13 | -0,66 |
| 599 | Dha_Kin_pool_2 | E | | E | A | 0,67 | 0,64 | 0,80 | 0,67 | 0,65 | 0,74 | 0,02 | 0,10 | 1,13 | | 0,64 | | 0,13 | 3,59 |
| 600 | Dha_Kin_pool_2 | E | | E | A | 0,70 | 0,53 | 0,72 | 0,76 | 0,62 | 0,74 | 0,12 | 0,03 | 1,20 | | 0,64 | | 0,13 | 4,10 |
| 601 | Dha_Kin_pool_2 | | 3611 | ILK | S | 1,01 | 1,17 | 0,53 | 0,56 | 1,09 | 0,55 | 0,11 | 0,02 | 0,50 | 0,50 | 0,64 | 0,14 | 0,13 | -1,05 |
| 602 | Dha_Kin_pool_2 | | 85481 | PSKH2 | S | 1,04 | 0,83 | 0,38 | 0,47 | 0,94 | 0,43 | 0,14 | 0,06 | 0,46 | 0,46 | 0,64 | 0,19 | 0,13 | -1,38 |
| 603 | Dha_Kin_pool_2 | | 29959 | NRBP | S | 0,78 | 0,61 | 0,44 | 0,26 | 0,70 | 0,35 | 0,12 | 0,13 | 0,50 | 0,50 | 0,64 | 0,14 | 0,13 | -1,06 |
| 604 | Dha_Kin_pool_2 | | 8408 | ULK1 | S | 0,68 | 0,55 | 0,30 | 0,13 | 0,61 | 0,21 | 0,09 | 0,12 | 0,34 | 0,34 | 0,64 | 0,30 | 0,13 | -2,22 |

| | | | | | | | | | | | | | | | | | | | |
|---|---|---|---|---|---|---|---|---|---|---|---|---|---|---|---|---|---|---|---|
| 605 | Dha_Kin_pool_2 | | 5578 | PRKCA | S | 0,66 | 0,62 | 0,45 | 0,29 | 0,64 | 0,37 | 0,03 | 0,11 | 0,58 | 0,58 | 0,64 | 0,06 | 0,13 | -0,46 |
| 606 | Dha_Kin_pool_2 | | 8999 | CDKL2 | S | 0,77 | 0,62 | 0,57 | 0,42 | 0,69 | 0,49 | 0,11 | 0,11 | 0,71 | 0,71 | 0,64 | 0,07 | 0,13 | 0,52 |
| 607 | Dha_Kin_pool_2 | | 2242 | FES | S | 0,41 | 0,48 | 0,47 | 0,31 | 0,44 | 0,39 | 0,05 | 0,11 | 0,88 | 0,88 | 0,64 | 0,24 | 0,13 | 1,75 |
| 608 | Dha_Kin_pool_2 | | 115701 | HAK | S | 0,58 | 0,52 | 0,40 | 0,23 | 0,55 | 0,32 | 0,04 | 0,12 | 0,58 | 0,58 | 0,64 | 0,06 | 0,13 | -0,48 |
| 609 | Dha_Kin_pool_2 | | 1017 | CDK2 | S | 0,65 | 0,69 | 0,45 | 0,36 | 0,67 | 0,41 | 0,03 | 0,06 | 0,61 | 0,61 | 0,64 | 0,04 | 0,13 | -0,27 |
| 610 | Dha_Kin_pool_2 | | 5605 | MAP2K2 | S | 0,52 | 0,54 | 0,53 | 0,30 | 0,53 | 0,41 | 0,01 | 0,16 | 0,78 | 0,78 | 0,64 | 0,14 | 0,13 | 1,03 |
| 611 | Dha_Kin_pool_2 | | 5602 | MAPK10 | S | 0,75 | 0,78 | 0,87 | 0,73 | 0,77 | 0,80 | 0,02 | 0,10 | 1,05 | 1,05 | 0,64 | 0,40 | 0,13 | 2,98 |
| 612 | Dha_Kin_pool_2 | | 3795 | KHK | S | 0,73 | 0,69 | 0,54 | 0,41 | 0,71 | 0,47 | 0,02 | 0,09 | 0,66 | 0,66 | 0,64 | 0,02 | 0,13 | 0,16 |
| 613 | Dha_Kin_pool_2 | | 2987 | GUK1 | S | 0,84 | 0,74 | 0,69 | 0,62 | 0,79 | 0,66 | 0,07 | 0,05 | 0,83 | 0,83 | 0,64 | 0,19 | 0,13 | 1,41 |
| 614 | Dha_Kin_pool_2 | | 1158 | CKM | S | 0,61 | 0,82 | 0,49 | 0,24 | 0,71 | 0,37 | 0,15 | 0,18 | 0,51 | 0,51 | 0,64 | 0,13 | 0,13 | -0,95 |
| 615 | Dha_Kin_pool_2 | | 8491 | MAP4K3 | S | 0,78 | 0,71 | 0,63 | 0,32 | 0,74 | 0,47 | 0,05 | 0,22 | 0,64 | 0,64 | 0,64 | 0,00 | 0,13 | -0,03 |
| 616 | Dha_Kin_pool_2 | | 5591 | PRKDC | S | 0,81 | 0,80 | 0,62 | 0,51 | 0,81 | 0,57 | 0,00 | 0,08 | 0,70 | 0,70 | 0,64 | 0,06 | 0,13 | 0,43 |
| 617 | Dha_Kin_pool_2 | | 8780 | RIOK3 | S | 1,01 | 1,00 | 0,68 | 0,45 | 1,01 | 0,56 | 0,01 | 0,16 | 0,56 | 0,56 | 0,64 | 0,08 | 0,13 | -0,61 |
| 618 | Dha_Kin_pool_2 | | 85443 | KIAA1765 | S | 0,99 | 0,88 | 0,59 | 0,56 | 0,93 | 0,57 | 0,08 | 0,02 | 0,61 | 0,61 | 0,64 | 0,03 | 0,13 | -0,23 |
| 619 | Dha_Kin_pool_2 | | 5230 | PGK1 | S | 0,47 | 0,48 | 0,42 | 0,29 | 0,48 | 0,36 | 0,01 | 0,09 | 0,75 | 0,75 | 0,64 | 0,10 | 0,13 | 0,76 |
| 620 | Dha_Kin_pool_2 | | 50808 | AK3L1 | S | 0,96 | 0,88 | 0,87 | 0,88 | 0,92 | 0,88 | 0,06 | 0,01 | 0,95 | 0,95 | 0,64 | 0,31 | 0,13 | 2,31 |
| 621 | Dha_Kin_pool_2 | | 1263 | PLK3 | S | 0,92 | 0,68 | 0,51 | 0,38 | 0,80 | 0,45 | 0,17 | 0,09 | 0,55 | 0,55 | 0,64 | 0,09 | 0,13 | -0,66 |
| 622 | Dha_Kin_pool_2 | | 2049 | EPHB3 | S | 1,05 | 0,89 | 1,03 | 0,73 | 0,97 | 0,88 | 0,11 | 0,21 | 0,90 | 0,90 | 0,64 | 0,26 | 0,13 | 1,93 |
| 623 | Dha_Kin_pool_2 | E | | E | A | 0,67 | 0,52 | 0,79 | 0,47 | 0,60 | 0,63 | 0,11 | 0,23 | 1,06 | | 0,64 | | 0,13 | 3,11 |
| 624 | Dha_Kin_pool_2 | E | | E | A | 0,59 | 0,59 | 0,57 | 0,58 | 0,59 | 0,57 | 0,00 | 0,01 | 0,97 | | 0,64 | | 0,13 | 2,43 |
| 625 | Dha_Kin_pool_2 | | 5579 | PRKCB1 | S | 0,88 | 0,82 | 0,60 | 0,64 | 0,85 | 0,62 | 0,04 | 0,03 | 0,73 | 0,73 | 0,64 | 0,09 | 0,13 | 0,67 |
| 626 | Dha_Kin_pool_2 | | 8527 | DGKD | S | 0,67 | 0,70 | 0,33 | 0,32 | 0,69 | 0,33 | 0,02 | 0,01 | 0,47 | 0,47 | 0,64 | 0,17 | 0,13 | -1,26 |
| 627 | Dha_Kin_pool_2 | | 2050 | EPHB4 | S | 0,89 | 0,73 | 0,53 | 0,38 | 0,81 | 0,46 | 0,11 | 0,11 | 0,56 | 0,56 | 0,64 | 0,08 | 0,13 | -0,58 |
| 628 | Dha_Kin_pool_2 | | 166614 | MGC45428 | S | 0,73 | 0,61 | 0,44 | 0,43 | 0,67 | 0,44 | 0,08 | 0,00 | 0,65 | 0,65 | 0,64 | 0,01 | 0,13 | 0,08 |
| 629 | Dha_Kin_pool_2 | | 1120 | CHKB | S | 0,55 | 0,46 | 0,39 | 0,38 | 0,50 | 0,38 | 0,06 | 0,01 | 0,76 | 0,76 | 0,64 | 0,12 | 0,13 | 0,86 |
| 630 | Dha_Kin_pool_2 | | 5106 | PCK2 | S | 0,53 | 0,58 | 0,43 | 0,30 | 0,55 | 0,36 | 0,03 | 0,09 | 0,66 | 0,66 | 0,64 | 0,01 | 0,13 | 0,10 |
| 631 | Dha_Kin_pool_2 | | 269 | AMHR2 | S | 0,73 | 0,95 | 0,48 | 0,44 | 0,84 | 0,46 | 0,16 | 0,03 | 0,55 | 0,55 | 0,64 | 0,10 | 0,13 | -0,71 |
| 632 | Dha_Kin_pool_2 | | 169436 | C9ORF96 | S | 0,62 | 0,66 | 0,47 | 0,37 | 0,64 | 0,42 | 0,03 | 0,07 | 0,66 | 0,66 | 0,64 | 0,02 | 0,13 | 0,14 |
| 633 | Dha_Kin_pool_2 | | 5604 | MAP2K1 | S | 0,71 | 0,69 | 0,55 | 0,47 | 0,70 | 0,51 | 0,02 | 0,05 | 0,73 | 0,73 | 0,64 | 0,08 | 0,13 | 0,61 |
| 634 | Dha_Kin_pool_2 | | 2585 | GALK2 | S | 0,78 | 0,77 | 0,55 | 0,48 | 0,78 | 0,52 | 0,01 | 0,05 | 0,67 | 0,67 | 0,64 | 0,02 | 0,13 | 0,16 |
| 635 | Dha_Kin_pool_2 | | 4914 | NTRK1 | S | 1,00 | 0,89 | 0,86 | 0,52 | 0,94 | 0,69 | 0,08 | 0,24 | 0,73 | 0,73 | 0,64 | 0,08 | 0,13 | 0,62 |
| 636 | Dha_Kin_pool_2 | | 4833 | NME4 | S | 0,65 | 0,61 | 0,57 | 0,53 | 0,63 | 0,55 | 0,02 | 0,03 | 0,87 | 0,87 | 0,64 | 0,23 | 0,13 | 1,68 |
| 637 | Dha_Kin_pool_2 | | 23178 | PASK | S | 0,88 | 0,46 | 0,35 | 0,24 | 0,67 | 0,30 | 0,30 | 0,08 | 0,45 | 0,45 | 0,64 | 0,20 | 0,13 | -1,45 |
| 638 | Dha_Kin_pool_2 | | 9874 | TLK1 | S | 0,82 | 0,59 | 0,62 | 0,42 | 0,70 | 0,52 | 0,16 | 0,14 | 0,74 | 0,74 | 0,64 | 0,09 | 0,13 | 0,69 |
| 639 | Dha_Kin_pool_2 | | 5213 | PFKM | S | 0,93 | 0,99 | 0,61 | 0,69 | 0,96 | 0,65 | 0,04 | 0,05 | 0,68 | 0,68 | 0,64 | 0,04 | 0,13 | 0,27 |
| 640 | Dha_Kin_pool_2 | | 5563 | PRKAA2 | S | 0,62 | 0,54 | 0,32 | 0,32 | 0,58 | 0,32 | 0,05 | 0,00 | 0,55 | 0,55 | 0,64 | 0,09 | 0,13 | -0,67 |
| 641 | Dha_Kin_pool_2 | | 3707 | ITPKB | S | 0,85 | 0,97 | 0,62 | 0,57 | 0,91 | 0,59 | 0,08 | 0,04 | 0,65 | 0,65 | 0,64 | 0,01 | 0,13 | 0,04 |
| 642 | Dha_Kin_pool_2 | | 4356 | MPP3 | S | 0,72 | 0,80 | 0,38 | 0,26 | 0,76 | 0,32 | 0,05 | 0,08 | 0,42 | 0,42 | 0,64 | 0,23 | 0,13 | -1,69 |
| 643 | Dha_Kin_pool_2 | | 1018 | CDK3 | S | 0,69 | 0,71 | 0,51 | 0,45 | 0,70 | 0,48 | 0,01 | 0,04 | 0,69 | 0,69 | 0,64 | 0,04 | 0,13 | 0,32 |
| 644 | Dha_Kin_pool_2 | | 5170 | PDPK1 | S | 0,41 | 0,40 | 0,50 | 0,42 | 0,41 | 0,46 | 0,01 | 0,06 | 1,12 | 1,12 | 0,64 | 0,48 | 0,13 | 3,57 |
| 645 | Dha_Kin_pool_2 | | 118672 | C10ORF89 | S | 0,99 | 0,85 | 0,83 | 0,72 | 0,92 | 0,78 | 0,10 | 0,07 | 0,85 | 0,85 | 0,64 | 0,20 | 0,13 | 1,50 |
| 646 | Dha_Kin_pool_2 | | 2043 | EPHA4 | S | 0,51 | 0,45 | 0,36 | 0,20 | 0,48 | 0,28 | 0,04 | 0,11 | 0,58 | 0,58 | 0,64 | 0,07 | 0,13 | -0,49 |
| 647 | Dha_Kin_pool_2 | E | | E | A | 0,68 | 0,60 | 0,69 | 0,55 | 0,64 | 0,62 | 0,06 | 0,10 | 0,97 | | 0,64 | | 0,13 | 2,39 |
| 648 | Dha_Kin_pool_2 | E | | E | A | 0,63 | 0,57 | 0,75 | 0,79 | 0,60 | 0,77 | 0,04 | 0,03 | 1,27 | | 0,64 | | 0,13 | 4,68 |
| 649 | Dha_Kin_pool_2 | | 9201 | DCAMKL1 | S | 0,65 | 0,60 | 0,31 | 0,27 | 0,63 | 0,29 | 0,03 | 0,03 | 0,46 | 0,46 | 0,64 | 0,18 | 0,13 | -1,34 |
| 650 | Dha_Kin_pool_2 | | 2712 | GK2 | S | 0,74 | 0,65 | 0,57 | 0,58 | 0,69 | 0,57 | 0,06 | 0,01 | 0,83 | 0,83 | 0,64 | 0,18 | 0,13 | 1,35 |
| 651 | Dha_Kin_pool_2 | | 54822 | TRPM7 | S | 0,49 | 0,43 | 0,28 | 0,22 | 0,46 | 0,25 | 0,04 | 0,04 | 0,54 | 0,54 | 0,64 | 0,10 | 0,13 | -0,76 |
| 652 | Dha_Kin_pool_2 | | 4638 | MYLK | S | 0,32 | 0,28 | 0,13 | 0,08 | 0,30 | 0,11 | 0,03 | 0,04 | 0,36 | 0,36 | 0,64 | 0,29 | 0,13 | -2,11 |
| 653 | Dha_Kin_pool_2 | | 2932 | GSK3B | S | 0,13 | 0,14 | 0,10 | 0,11 | 0,14 | 0,11 | 0,01 | 0,00 | 0,77 | 0,77 | 0,64 | 0,12 | 0,13 | 0,91 |
| 654 | Dha_Kin_pool_2 | | 4486 | MST1R | S | 0,83 | 0,77 | 0,64 | 0,49 | 0,80 | 0,56 | 0,04 | 0,11 | 0,70 | 0,70 | 0,64 | 0,06 | 0,13 | 0,46 |
| 655 | Dha_Kin_pool_2 | | 204 | AK2 | S | 0,77 | 0,87 | 0,65 | 0,50 | 0,82 | 0,58 | 0,07 | 0,11 | 0,70 | 0,70 | 0,64 | 0,06 | 0,13 | 0,44 |
| 656 | Dha_Kin_pool_2 | | 140609 | NEK7 | S | 0,68 | 0,65 | 0,32 | 0,17 | 0,67 | 0,24 | 0,02 | 0,11 | 0,37 | 0,37 | 0,64 | 0,28 | 0,13 | -2,06 |
| 657 | Dha_Kin_pool_2 | | 65125 | PRKWNK1 | S | 0,62 | 0,66 | 0,40 | 0,26 | 0,64 | 0,33 | 0,03 | 0,10 | 0,52 | 0,52 | 0,64 | 0,13 | 0,13 | -0,94 |
| 658 | Dha_Kin_pool_2 | | 60385 | TSKS | S | 0,63 | 0,53 | 0,58 | 0,36 | 0,58 | 0,47 | 0,07 | 0,16 | 0,81 | 0,81 | 0,64 | 0,16 | 0,13 | 1,21 |
| 659 | Dha_Kin_pool_2 | | 8859 | STK19 | S | 0,37 | 0,39 | 0,35 | 0,19 | 0,38 | 0,27 | 0,01 | 0,11 | 0,70 | 0,70 | 0,64 | 0,06 | 0,13 | 0,43 |

| | | | | | | | | | | | | | | | | | | | |
|---|---|---|---|---|---|---|---|---|---|---|---|---|---|---|---|---|---|---|---|
| 660 | Dha_Kin_pool_2 | | 6300 | MAPK12 | S | 0,35 | 0,34 | 0,36 | 0,19 | 0,35 | 0,28 | 0,00 | 0,12 | 0,79 | 0,79 | 0,64 | 0,15 | 0,13 | 1,10 |
| 661 | Dha_Kin_pool_2 | | 2584 | GALK1 | S | 0,38 | 0,33 | 0,44 | 0,30 | 0,35 | 0,37 | 0,03 | 0,10 | 1,05 | 1,05 | 0,64 | 0,41 | 0,13 | 3,04 |
| 662 | Dha_Kin_pool_2 | | 27231 | NMRK2 | S | 0,81 | 0,84 | 0,78 | 0,66 | 0,83 | 0,72 | 0,02 | 0,08 | 0,87 | 0,87 | 0,64 | 0,23 | 0,13 | 1,68 |
| 663 | Dha_Kin_pool_2 | | 7083 | TK1 | S | 1,10 | 1,13 | 0,57 | 0,67 | 1,12 | 0,62 | 0,02 | 0,07 | 0,55 | 0,55 | 0,64 | 0,09 | 0,13 | -0,66 |
| 664 | Dha_Kin_pool_2 | | 9479 | MAPK8IP1 | S | 0,61 | 0,51 | 0,48 | 0,50 | 0,56 | 0,49 | 0,07 | 0,01 | 0,87 | 0,87 | 0,64 | 0,23 | 0,13 | 1,69 |
| 665 | Dha_Kin_pool_2 | | 8525 | DGKZ | S | 0,71 | 0,79 | 0,65 | 0,42 | 0,75 | 0,54 | 0,06 | 0,16 | 0,71 | 0,71 | 0,64 | 0,07 | 0,13 | 0,51 |
| 666 | Dha_Kin_pool_2 | | 8575 | PRKRA | S | 0,87 | 0,69 | 0,53 | 0,40 | 0,78 | 0,46 | 0,12 | 0,09 | 0,60 | 0,60 | 0,64 | 0,05 | 0,13 | -0,36 |
| 667 | Dha_Kin_pool_2 | | 1796 | DOK1 | S | 0,59 | 0,54 | 0,28 | 0,32 | 0,57 | 0,30 | 0,04 | 0,03 | 0,53 | 0,53 | 0,64 | 0,11 | 0,13 | -0,80 |
| 668 | Dha_Kin_pool_2 | | 154043 | CNKSR3 | S | 0,58 | 0,55 | 0,51 | 0,33 | 0,57 | 0,42 | 0,02 | 0,13 | 0,74 | 0,74 | 0,64 | 0,10 | 0,13 | 0,72 |
| 669 | Dha_Kin_pool_2 | | 26007 | DAK | S | 1,10 | 1,20 | 0,93 | 0,86 | 1,15 | 0,89 | 0,07 | 0,04 | 0,78 | 0,78 | 0,64 | 0,14 | 0,13 | 1,01 |
| 670 | Dha_Kin_pool_2 | | 7294 | TXK | S | 1,12 | 1,07 | 0,88 | 0,87 | 1,09 | 0,87 | 0,04 | 0,01 | 0,80 | 0,80 | 0,64 | 0,15 | 0,13 | 1,14 |
| 671 | Dha_Kin_pool_2 | E | | E | A | 0,65 | 0,68 | 0,77 | 0,59 | 0,67 | 0,68 | 0,03 | 0,13 | 1,02 | | 0,64 | | 0,13 | 2,81 |
| 672 | Dha_Kin_pool_2 | E | | E | A | 0,66 | 0,61 | 0,72 | 0,65 | 0,63 | 0,69 | 0,04 | 0,05 | 1,08 | | 0,64 | | 0,13 | 3,26 |
| 673 | Dha_Kin_pool_2 | | 51314 | NME8 | S | 1,00 | 1,06 | 0,66 | 0,72 | 1,03 | 0,69 | 0,04 | 0,04 | 0,67 | 0,67 | 0,64 | 0,02 | 0,13 | 0,18 |
| 674 | Dha_Kin_pool_2 | | 58538 | MPP4 | S | 0,82 | 0,78 | 0,64 | 0,54 | 0,80 | 0,59 | 0,03 | 0,07 | 0,74 | 0,74 | 0,64 | 0,10 | 0,13 | 0,72 |
| 675 | Dha_Kin_pool_2 | | 132158 | GLYCTK | S | 0,73 | 0,65 | 0,62 | 0,53 | 0,69 | 0,57 | 0,05 | 0,07 | 0,83 | 0,83 | 0,64 | 0,19 | 0,13 | 1,41 |
| 676 | Dha_Kin_pool_2 | | 7075 | TIE1 | S | 0,83 | 0,76 | 0,70 | 0,44 | 0,80 | 0,57 | 0,04 | 0,18 | 0,72 | 0,72 | 0,64 | 0,08 | 0,13 | 0,56 |
| 677 | Dha_Kin_pool_2 | | 64398 | MPP5 | S | 0,71 | 0,72 | 0,78 | 0,68 | 0,72 | 0,73 | 0,00 | 0,07 | 1,01 | 1,01 | 0,64 | 0,37 | 0,13 | 2,75 |
| 678 | Dha_Kin_pool_2 | | 80201 | HKDC1 | S | 0,75 | 0,67 | 0,52 | 0,49 | 0,71 | 0,50 | 0,05 | 0,02 | 0,71 | 0,71 | 0,64 | 0,06 | 0,13 | 0,47 |
| 679 | Dha_Kin_pool_2 | | 51678 | MPP6 | S | 0,69 | 0,76 | 0,56 | 0,41 | 0,73 | 0,49 | 0,05 | 0,11 | 0,67 | 0,67 | 0,64 | 0,03 | 0,13 | 0,19 |
| 680 | Dha_Kin_pool_2 | | 9463 | PICK1 | S | 0,65 | 0,59 | 0,67 | 0,49 | 0,62 | 0,58 | 0,04 | 0,12 | 0,94 | 0,94 | 0,64 | 0,29 | 0,13 | 2,18 |
| 681 | Dha_Kin_pool_2 | | 8526 | DGKE | S | 0,85 | 1,02 | 0,79 | 0,59 | 0,94 | 0,69 | 0,12 | 0,14 | 0,73 | 0,73 | 0,64 | 0,09 | 0,13 | 0,66 |
| 682 | Dha_Kin_pool_2 | | 5756 | TWF1 | S | 0,85 | 0,70 | 0,54 | 0,51 | 0,77 | 0,52 | 0,10 | 0,02 | 0,68 | 0,68 | 0,64 | 0,03 | 0,13 | 0,26 |
| 683 | Dha_Kin_pool_2 | | 3815 | KIT | S | 0,79 | 0,73 | 0,78 | 0,40 | 0,76 | 0,59 | 0,05 | 0,27 | 0,78 | 0,78 | 0,64 | 0,13 | 0,13 | 0,98 |
| 684 | Dha_Kin_pool_2 | | 80347 | COASY | S | 0,92 | 0,72 | 0,64 | 0,38 | 0,82 | 0,51 | 0,14 | 0,18 | 0,63 | 0,63 | 0,64 | 0,01 | 0,13 | -0,10 |
| 685 | Dha_Kin_pool_2 | | 23533 | PIK3R5 | S | 0,29 | 0,25 | 0,28 | 0,14 | 0,27 | 0,21 | 0,03 | 0,10 | 0,79 | 0,79 | 0,64 | 0,15 | 0,13 | 1,08 |
| 686 | Dha_Kin_pool_2 | | 10221 | TRIB1 | S | 0,52 | 0,64 | 0,35 | 0,36 | 0,58 | 0,35 | 0,09 | 0,01 | 0,61 | 0,61 | 0,64 | 0,04 | 0,13 | -0,28 |
| 687 | Dha_Kin_pool_2 | | 55437 | STRADB | S | 0,84 | 0,97 | 0,85 | 0,65 | 0,91 | 0,75 | 0,09 | 0,14 | 0,82 | 0,82 | 0,64 | 0,18 | 0,13 | 1,34 |
| 688 | Dha_Kin_pool_2 | | 51232 | CRIM1 | S | 0,93 | 0,68 | 0,31 | 0,23 | 0,81 | 0,27 | 0,17 | 0,05 | 0,34 | 0,34 | 0,64 | 0,31 | 0,13 | -2,27 |
| 689 | Dha_Kin_pool_2 | | 3480 | IGF1R | S | 0,68 | 0,71 | 0,48 | 0,44 | 0,69 | 0,46 | 0,03 | 0,02 | 0,66 | 0,66 | 0,64 | 0,02 | 0,13 | 0,12 |
| 690 | Dha_Kin_pool_2 | | 5296 | PIK3R2 | S | 0,22 | 0,22 | 0,20 | 0,15 | 0,22 | 0,18 | 0,00 | 0,04 | 0,81 | 0,81 | 0,64 | 0,17 | 0,13 | 1,26 |
| 691 | Dha_Kin_pool_2 | | 8631 | SKAP1 | S | 0,80 | 0,62 | 0,31 | 0,21 | 0,71 | 0,26 | 0,13 | 0,07 | 0,36 | 0,36 | 0,64 | 0,28 | 0,13 | -2,07 |
| 692 | Dha_Kin_pool_2 | | 2324 | FLT4 | S | 0,94 | 0,97 | 0,81 | 0,59 | 0,95 | 0,70 | 0,02 | 0,16 | 0,73 | 0,73 | 0,64 | 0,09 | 0,13 | 0,68 |
| 693 | Dha_Kin_pool_2 | | 5576 | PRKAR2A | S | 1,03 | 1,16 | 0,77 | 0,78 | 1,09 | 0,78 | 0,09 | 0,01 | 0,71 | 0,71 | 0,64 | 0,07 | 0,13 | 0,50 |
| 694 | Dha_Kin_pool_2 | | 8503 | PIK3R3 | S | 0,55 | 0,62 | 0,41 | 0,34 | 0,59 | 0,38 | 0,05 | 0,05 | 0,64 | 0,64 | 0,64 | 0,00 | 0,13 | -0,03 |
| 695 | Dha_Kin_pool_2 | E | | E | A | 0,63 | 0,77 | 0,90 | 0,76 | 0,70 | 0,83 | 0,10 | 0,10 | 1,18 | | 0,64 | | 0,13 | 3,98 |
| 696 | Dha_Kin_pool_2 | E | | E | A | 0,65 | 0,63 | 0,63 | 0,61 | 0,64 | 0,62 | 0,01 | 0,02 | 0,97 | | 0,64 | | 0,13 | 2,40 |
| 697 | Dha_Kin_pool_2 | | 8941 | CDK5R2 | S | 0,89 | 0,99 | 0,75 | 0,75 | 0,94 | 0,75 | 0,07 | 0,00 | 0,80 | 0,80 | 0,64 | 0,16 | 0,13 | 1,16 |
| 698 | Dha_Kin_pool_2 | | 5832 | ALDH18A1 | S | 0,86 | 0,82 | 0,51 | 0,52 | 0,84 | 0,51 | 0,02 | 0,01 | 0,61 | 0,61 | 0,64 | 0,03 | 0,13 | -0,24 |
| 699 | Dha_Kin_pool_2 | | 5634 | PRPS2 | S | 0,88 | 0,75 | 0,77 | 0,63 | 0,82 | 0,70 | 0,09 | 0,10 | 0,86 | 0,86 | 0,64 | 0,22 | 0,13 | 1,60 |
| 700 | Dha_Kin_pool_2 | | 8851 | CDK5R1 | S | 0,50 | 0,47 | 0,43 | 0,25 | 0,49 | 0,34 | 0,02 | 0,12 | 0,70 | 0,70 | 0,64 | 0,06 | 0,13 | 0,45 |
| 701 | Dha_Kin_pool_2 | | 5573 | PRKAR1A | S | 0,57 | 0,43 | 0,47 | 0,27 | 0,50 | 0,37 | 0,10 | 0,14 | 0,74 | 0,74 | 0,64 | 0,09 | 0,13 | 0,70 |
| 702 | Dha_Kin_pool_2 | | 30849 | PIK3R4 | S | 0,63 | 0,70 | 0,30 | 0,25 | 0,67 | 0,28 | 0,05 | 0,03 | 0,41 | 0,41 | 0,64 | 0,23 | 0,13 | -1,71 |
| 703 | Dha_Kin_pool_2 | | 5575 | PRKAR1B | S | 0,66 | 0,66 | 0,62 | 0,34 | 0,66 | 0,48 | 0,00 | 0,20 | 0,73 | 0,73 | 0,64 | 0,08 | 0,13 | 0,61 |
| 704 | Dha_Kin_pool_2 | | 28951 | TRIB2 | S | 0,52 | 0,64 | 0,66 | 0,49 | 0,58 | 0,57 | 0,09 | 0,12 | 0,98 | 0,98 | 0,64 | 0,34 | 0,13 | 2,52 |
| 705 | Dha_Kin_pool_2 | | 8518 | IKBKAP | S | 0,83 | 0,62 | 0,80 | 0,46 | 0,73 | 0,63 | 0,15 | 0,24 | 0,87 | 0,87 | 0,64 | 0,23 | 0,13 | 1,68 |
| 706 | Dha_Kin_pool_2 | | 57761 | TRIB3 | S | 0,28 | 0,21 | 0,35 | 0,19 | 0,25 | 0,27 | 0,05 | 0,11 | 1,08 | 1,08 | 0,64 | 0,44 | 0,13 | 3,27 |
| 707 | Dha_Kin_pool_2 | | 5631 | PRPS1 | S | 0,59 | 0,60 | 0,50 | 0,25 | 0,59 | 0,37 | 0,01 | 0,17 | 0,63 | 0,63 | 0,64 | 0,02 | 0,13 | -0,11 |
| 708 | Dha_Kin_pool_2 | | 57410 | SCYL1 | S | 0,91 | 0,86 | 0,68 | 0,60 | 0,89 | 0,64 | 0,04 | 0,06 | 0,72 | 0,72 | 0,64 | 0,08 | 0,13 | 0,56 |
| 709 | Dha_Kin_pool_2 | | 57147 | SCYL3 | S | 0,20 | 0,16 | 0,36 | 0,12 | 0,18 | 0,24 | 0,03 | 0,17 | 1,32 | 1,32 | 0,64 | 0,67 | 0,13 | 4,99 |
| 710 | Dha_Kin_pool_2 | | 5295 | PIK3R1 | S | 0,71 | 0,82 | 0,69 | 0,42 | 0,76 | 0,55 | 0,08 | 0,19 | 0,72 | 0,72 | 0,64 | 0,08 | 0,13 | 0,59 |
| 711 | Dha_Kin_pool_2 | | 92335 | STRADA | S | 0,74 | 0,82 | 0,34 | 0,36 | 0,78 | 0,35 | 0,06 | 0,02 | 0,45 | 0,45 | 0,64 | 0,20 | 0,13 | -1,47 |
| 712 | Dha_Kin_pool_2 | | 6725 | SRMS | S | 0,76 | 0,68 | 0,62 | 0,64 | 0,72 | 0,63 | 0,05 | 0,01 | 0,87 | 0,87 | 0,64 | 0,23 | 0,13 | 1,69 |
| 713 | Dha_Kin_pool_2 | | 221823 | PRPS1L1 | S | 0,91 | 1,07 | 0,82 | 0,56 | 0,99 | 0,69 | 0,11 | 0,19 | 0,70 | 0,70 | 0,64 | 0,05 | 0,13 | 0,39 |
| 714 | Dha_Kin_pool_2 | | 11344 | TWF2 | S | 0,46 | 0,42 | 0,23 | 0,17 | 0,44 | 0,20 | 0,03 | 0,04 | 0,45 | 0,45 | 0,64 | 0,19 | 0,13 | -1,44 |

| | | | | | | | | | | | | | | | | | | | |
|---|---|---|---|---|---|---|---|---|---|---|---|---|---|---|---|---|---|---|---|
| 715 | Dha_Kin_pool_2 | | 8899 | PRPF4B | S | 0,77 | 0,93 | 0,69 | 0,46 | 0,85 | 0,57 | 0,11 | 0,17 | 0,67 | 0,67 | 0,64 | 0,03 | 0,13 | 0,23 |
| 716 | Dha_Kin_pool_2 | | 5577 | PRKAR2B | S | 0,65 | 0,69 | 0,66 | 0,48 | 0,67 | 0,57 | 0,03 | 0,13 | 0,85 | 0,85 | 0,64 | 0,21 | 0,13 | 1,56 |
| 717 | Dha_Kin_pool_2 | | 673 | BRAF | S | 0,69 | 0,95 | 0,37 | 0,33 | 0,82 | 0,35 | 0,19 | 0,03 | 0,43 | 0,43 | 0,64 | 0,22 | 0,13 | -1,61 |
| 718 | Dha_Kin_pool_2 | E | | E | E | 1,25 | 1,39 | 0,94 | 0,85 | 1,32 | 0,89 | 0,10 | 0,06 | 0,68 | | 0,64 | | 0,13 | 0,25 |
| 719 | Dha_Kin_pool_2 | E | | E | A | 0,67 | 0,63 | 0,80 | 0,62 | 0,65 | 0,71 | 0,03 | 0,13 | 1,10 | | 0,64 | | 0,13 | 3,36 |
| 720 | Dha_Kin_pool_2 | E | | E | A | 0,57 | 0,66 | 0,69 | 0,60 | 0,62 | 0,64 | 0,07 | 0,07 | 1,04 | | 0,64 | | 0,13 | 2,94 |
| 721 | Dha_Kin_pool_2 | E | | E | E | 1,20 | 1,05 | 0,81 | 0,76 | 1,13 | 0,79 | 0,11 | 0,03 | 0,70 | | 0,64 | | 0,13 | 0,41 |
| 722 | Dha_Kin_pool_2 | E | | E | E | 1,09 | 1,14 | 0,83 | 0,80 | 1,12 | 0,82 | 0,03 | 0,02 | 0,73 | | 0,64 | | 0,13 | 0,63 |
| 723 | Dha_Kin_pool_2 | E | | E | E | 1,15 | 1,09 | 0,74 | 0,70 | 1,12 | 0,72 | 0,05 | 0,03 | 0,65 | | 0,64 | | 0,13 | 0,02 |
| 724 | Dha_Kin_pool_2 | E | | E | E | 0,98 | 1,05 | 0,85 | 0,75 | 1,02 | 0,80 | 0,05 | 0,07 | 0,79 | | 0,64 | | 0,13 | 1,05 |
| 725 | Dha_Kin_pool_2 | E | | E | E | 1,02 | 0,94 | 0,72 | 0,73 | 0,98 | 0,73 | 0,05 | 0,01 | 0,74 | | 0,64 | | 0,13 | 0,73 |
| 726 | Dha_Kin_pool_2 | E | | E | E | 1,00 | 0,86 | 0,68 | 0,75 | 0,93 | 0,72 | 0,10 | 0,05 | 0,77 | | 0,64 | | 0,13 | 0,95 |
| 727 | Dha_Kin_pool_2 | E | | E | E | 1,03 | 0,88 | 0,73 | 0,84 | 0,95 | 0,78 | 0,11 | 0,08 | 0,82 | | 0,64 | | 0,13 | 1,34 |
| 728 | Dha_Kin_pool_2 | E | | E | E | 1,05 | 0,92 | 0,77 | 0,89 | 0,98 | 0,83 | 0,09 | 0,09 | 0,85 | | 0,64 | | 0,13 | 1,50 |
| 729 | Dha_Kin_pool_2 | E | | E | E | 1,04 | 0,95 | 0,70 | 0,58 | 0,99 | 0,64 | 0,06 | 0,09 | 0,64 | | 0,64 | | 0,13 | 0,00 |
| 730 | Dha_Kin_pool_2 | E | | E | E | 0,91 | 0,88 | 0,76 | 0,74 | 0,89 | 0,75 | 0,02 | 0,01 | 0,84 | | 0,64 | | 0,13 | 1,45 |
| 731 | Dha_Kin_pool_2 | E | | E | E | 1,00 | 1,02 | 0,91 | 0,92 | 1,01 | 0,91 | 0,02 | 0,01 | 0,91 | | 0,64 | | 0,13 | 1,94 |
| 732 | Dha_Kin_pool_2 | E | | E | E | 0,99 | 0,96 | 0,91 | 0,77 | 0,97 | 0,84 | 0,02 | 0,10 | 0,86 | | 0,64 | | 0,13 | 1,61 |
| 733 | Dha_Kin_pool_2 | E | | E | E | 0,96 | 1,02 | 0,96 | 0,81 | 0,99 | 0,89 | 0,04 | 0,10 | 0,90 | | 0,64 | | 0,13 | 1,88 |
| 734 | Dha_Kin_pool_2 | E | | E | E | 1,00 | 1,10 | 1,01 | 0,72 | 1,05 | 0,87 | 0,07 | 0,20 | 0,83 | | 0,64 | | 0,13 | 1,35 |
| 735 | Dha_Kin_pool_2 | E | | E | E | 0,99 | 1,09 | 0,92 | 0,69 | 1,04 | 0,80 | 0,07 | 0,16 | 0,77 | | 0,64 | | 0,13 | 0,96 |
| 736 | Dha_Kin_pool_2 | E | | E | E | 1,08 | 1,15 | 0,88 | 0,71 | 1,11 | 0,79 | 0,05 | 0,12 | 0,71 | | 0,64 | | 0,13 | 0,51 |
| 737 | Dha_Kin_pool_2 | E | | E | E | 1,11 | 1,20 | 0,67 | 0,82 | 1,15 | 0,75 | 0,06 | 0,11 | 0,65 | | 0,64 | | 0,13 | 0,03 |
| 738 | Dha_Kin_pool_2 | E | | E | E | 1,14 | 0,96 | 0,79 | 0,86 | 1,05 | 0,82 | 0,13 | 0,05 | 0,78 | | 0,64 | | 0,13 | 1,04 |
| 739 | Dha_Kin_pool_2 | E | | E | E | 1,27 | 1,26 | 0,97 | 0,89 | 1,26 | 0,93 | 0,01 | 0,06 | 0,74 | | 0,64 | | 0,13 | 0,68 |
| 740 | Dha_Kin_pool_2 | E | | E | E | 1,21 | 1,22 | 0,85 | 0,75 | 1,21 | 0,80 | 0,01 | 0,08 | 0,66 | | 0,64 | | 0,13 | 0,12 |
| 741 | Dha_Kin_pool_2 | E | | E | E | 1,26 | 1,32 | 1,04 | 0,87 | 1,29 | 0,95 | 0,04 | 0,12 | 0,74 | | 0,64 | | 0,13 | 0,70 |
| 742 | Dha_Kin_pool_2 | E | | E | E | 1,30 | 1,27 | 0,94 | 0,77 | 1,28 | 0,86 | 0,02 | 0,12 | 0,67 | | 0,64 | | 0,13 | 0,19 |
| 743 | Dha_Kin_pool_2 | E | | E | A | 0,77 | 0,58 | 0,76 | 0,69 | 0,68 | 0,73 | 0,13 | 0,05 | 1,07 | | 0,64 | | 0,13 | 3,18 |
| 744 | Dha_Kin_pool_2 | E | | E | A | 0,58 | 0,60 | 0,72 | 0,62 | 0,59 | 0,67 | 0,02 | 0,07 | 1,13 | | 0,64 | | 0,13 | 3,61 |
| 745 | Dha_Kin_pool_2 | E | | E | E | 1,00 | 0,95 | 0,77 | 0,48 | 0,98 | 0,63 | 0,03 | 0,20 | 0,64 | | 0,64 | | 0,13 | -0,03 |
| 746 | Dha_Kin_pool_2 | E | | E | E | 1,03 | 1,11 | 0,70 | 0,62 | 1,07 | 0,66 | 0,06 | 0,06 | 0,62 | | 0,64 | | 0,13 | -0,19 |
| 747 | Dha_Kin_pool_2 | E | | E | E | 1,04 | 0,94 | 0,80 | 0,74 | 0,99 | 0,77 | 0,06 | 0,05 | 0,78 | | 0,64 | | 0,13 | 0,98 |
| 748 | Dha_Kin_pool_2 | E | | E | E | 1,12 | 0,95 | 0,85 | 0,72 | 1,03 | 0,79 | 0,12 | 0,09 | 0,76 | | 0,64 | | 0,13 | 0,86 |
| 749 | Dha_Kin_pool_2 | E | | E | E | 1,15 | 0,98 | 0,70 | 0,80 | 1,06 | 0,75 | 0,12 | 0,07 | 0,70 | | 0,64 | | 0,13 | 0,46 |
| 750 | Dha_Kin_pool_2 | E | | E | E | 0,88 | 0,99 | 0,75 | 0,79 | 0,93 | 0,77 | 0,08 | 0,03 | 0,83 | | 0,64 | | 0,13 | 1,37 |
| 751 | Dha_Kin_pool_2 | E | | E | E | 1,09 | 0,98 | 0,83 | 0,58 | 1,03 | 0,70 | 0,08 | 0,18 | 0,68 | | 0,64 | | 0,13 | 0,28 |
| 752 | Dha_Kin_pool_2 | E | | E | E | 0,86 | 1,04 | 0,71 | 0,70 | 0,95 | 0,71 | 0,13 | 0,01 | 0,74 | | 0,64 | | 0,13 | 0,73 |
| 753 | Dha_Kin_pool_2 | E | | E | E | 0,96 | 0,80 | 0,92 | 0,72 | 0,88 | 0,82 | 0,12 | 0,15 | 0,93 | | 0,64 | | 0,13 | 2,12 |
| 754 | Dha_Kin_pool_2 | E | | E | E | 1,19 | 0,97 | 0,76 | 0,60 | 1,08 | 0,68 | 0,16 | 0,11 | 0,63 | | 0,64 | | 0,13 | -0,11 |
| 755 | Dha_Kin_pool_2 | E | | E | E | 0,99 | 0,95 | 0,65 | 0,72 | 0,97 | 0,69 | 0,03 | 0,05 | 0,71 | | 0,64 | | 0,13 | 0,51 |
| 756 | Dha_Kin_pool_2 | E | | E | E | 1,09 | 0,93 | 0,86 | 0,74 | 1,01 | 0,80 | 0,11 | 0,08 | 0,79 | | 0,64 | | 0,13 | 1,12 |
| 757 | Dha_Kin_pool_2 | E | | E | E | 0,92 | 1,03 | 0,89 | 0,82 | 0,97 | 0,85 | 0,08 | 0,05 | 0,88 | | 0,64 | | 0,13 | 1,74 |
| 758 | Dha_Kin_pool_2 | E | | E | E | 0,92 | 1,08 | 0,85 | 0,63 | 1,00 | 0,74 | 0,11 | 0,15 | 0,74 | | 0,64 | | 0,13 | 0,72 |
| 759 | Dha_Kin_pool_2 | E | | E | E | 0,98 | 1,10 | 0,87 | 0,70 | 1,04 | 0,79 | 0,09 | 0,12 | 0,76 | | 0,64 | | 0,13 | 0,85 |
| 760 | Dha_Kin_pool_2 | E | | E | E | 0,99 | 1,02 | 0,73 | 0,59 | 1,01 | 0,66 | 0,02 | 0,10 | 0,66 | | 0,64 | | 0,13 | 0,09 |
| 761 | Dha_Kin_pool_2 | E | | E | E | 1,02 | 0,97 | 0,83 | 0,57 | 0,99 | 0,70 | 0,04 | 0,19 | 0,70 | | 0,64 | | 0,13 | 0,45 |
| 762 | Dha_Kin_pool_2 | E | | E | E | 1,11 | 1,05 | 0,86 | 0,67 | 1,08 | 0,76 | 0,05 | 0,14 | 0,71 | | 0,64 | | 0,13 | 0,46 |
| 763 | Dha_Kin_pool_2 | E | | E | E | 0,99 | 1,03 | 0,93 | 0,67 | 1,01 | 0,80 | 0,03 | 0,18 | 0,79 | | 0,64 | | 0,13 | 1,06 |
| 764 | Dha_Kin_pool_2 | E | | E | E | 1,27 | 0,99 | 0,84 | 0,74 | 1,13 | 0,79 | 0,20 | 0,07 | 0,70 | | 0,64 | | 0,13 | 0,44 |
| 765 | Dha_Kin_pool_2 | E | | E | E | 1,20 | 1,00 | 0,86 | 0,72 | 1,10 | 0,79 | 0,14 | 0,10 | 0,72 | | 0,64 | | 0,13 | 0,55 |
| 766 | Dha_Kin_pool_2 | E | | E | E | 1,17 | 0,98 | 0,89 | 0,87 | 1,08 | 0,88 | 0,13 | 0,01 | 0,82 | | 0,64 | | 0,13 | 1,31 |
| 767 | Dha_Kin_pool_2 | E | | E | A | 0,64 | 0,52 | 0,59 | 0,58 | 0,58 | 0,58 | 0,08 | 0,01 | 1,00 | | 0,64 | | 0,13 | 2,64 |
| 768 | Dha_Kin_pool_2 | E | | E | K | 0,01 | 0,01 | 0,02 | 0,02 | 0,01 | 0,02 | 0,00 | 0,00 | 1,60 | | 0,64 | | 0,13 | 7,10 |
