## Supplemental Table 2 for "The EGFR-STYK1-FGF1 axis sustains functional drug tolerance to EGFR inhibitors in EGFR-mutant non-small cell lung cancer"

| | | siRNA2-AFA vs ctrl-AFA | | | siRNA3-AFA vs ctrl-AFA | | |
|---|---|---|---|---|---|---|---|
| Gene | | Basemean | log2FoldChange | Adjusted p-value | Basemean | log2FoldChange | Adjusted p-value |
| ENSG00000074410.9 | CA12 | 43.0379488587795 | -1.51532896749452 | 3.72052258434022e-08 | 36.2296594233575 | -2.81386026511085 | 6.88419400857947e-16 |
| ENSG00000130595.12 | TNNT3 | 4.59615775579268 | -2.73350683750916 | 0.0113798360116411 | 4.5516626670481 | -2.72473753524772 | 0.0173544147038365 |
| ENSG00000184908.13 | CLCNKB | 11.2952899263227 | -1.77017855475872 | 0.0060920686869027 | 9.97024598211669 | -2.70214709654657 | 0.000662374636327962 |
| ENSG00000060140.4 | STYK1 | 64.8461661956939 | -2.72918870572451 | 7.82183352776054e-27 | 64.6310801995866 | -2.68492711980512 | 5.00914336755756e-26 |
| ENSG00000154734.10 | ADAMTS1 | 25.6608151134994 | -2.11469863796892 | 6.87007902885386e-09 | 24.9012801314993 | -2.28860948300724 | 2.03648201826156e-09 |
| ENSG00000149591.12 | TAGLN | 189.394832162226 | -3.76598270533873 | 3.1936710630402e-106 | 211.046900699864 | -2.28350664895344 | 4.9011768274029e-72 |
| ENSG00000152779.12 | SLC16A12 | 7.35494252249748 | -3.49388022062112 | 4.68043221893903e-05 | 8.12381864029478 | -2.21834887244446 | 0.00228064059881637 |
| ENSG00000072163.14 | LIMS2 | 15.4949421890533 | -3.2611602917794 | 7.24382289803177e-08 | 17.0403353549047 | -2.14023749614995 | 5.74716171853852e-06 |
| ENSG00000115232.9 | ITGA4 | 14.6070931239411 | -2.33635000982425 | 7.93654405902259e-06 | 14.8515610985139 | -2.12222608870065 | 4.40389411903727e-05 |
| ENSG00000101335.5 | MYL9 | 635.940921550798 | -3.1520841281099 | 3.69609972220055e-294 | 703.380366174956 | -2.05655528676916 | 2.84120976774742e-171 |
| ENSG00000157856.6 | DRC1 | 13.3594559033019 | -1.32024757128035 | 0.00883941300273772 | 11.750102225166 | -2.04773397124479 | 0.000347568025112384 |
| ENSG00000142494.9 | SLC47A1 | 9.36251793501993 | -1.42501788482037 | 0.0242644473456356 | 8.47890346891006 | -1.98586720569472 | 0.004626423943988 |
| ENSG00000140682.14 | TGFB1I1 | 160.852835575777 | -2.20997854498339 | 1.47998676762927e-49 | 165.586938296476 | -1.929726851408 | 1.84468177712181e-40 |
| ENSG00000103546.14 | SLC6A2 | 20.1318896887843 | -1.35899216520744 | 0.000609818070670236 | 18.2232846444039 | -1.87043189431881 | 0.000154863580269584 |
| ENSG00000176928.4 | GCNT4 | 79.1669177443431 | -2.21142560969531 | 6.74549577598967e-28 | 83.7920313363285 | -1.75434368627166 | 6.72745638687505e-20 |
| ENSG00000204941.9 | PSG5 | 39.2414679759204 | -3.11571174334792 | 1.23935734500966e-19 | 45.4065606813388 | -1.74725868508034 | 9.04970384523984e-11 |
| ENSG00000102760.12 | RGCC | 14.8877835860182 | -1.51812206610662 | 0.000928642278586593 | 14.310465938972 | -1.69367563995393 | 0.000867529436197998 |
| ENSG00000141750.6 | STAC2 | 8.20747575043649 | -4.44864310243336 | 1.24522074233982e-05 | 10.1974657896162 | -1.68824907540881 | 0.00517123566254375 |
| ENSG00000100346.13 | CACNA1I | 35.2608023900889 | -1.86604557217915 | 9.4284218494719e-10 | 36.0197691958691 | -1.68775921088606 | 1.40837950001902e-07 |
| ENSG00000145934.11 | TENM2 | 10.7475227403228 | -2.00810443062901 | 0.000455367770119611 | 11.2458600794151 | -1.68070833572055 | 0.00635847637171497 |
| ENSG00000130176.3 | CNN1 | 25.0346041790161 | -6.69226336162875 | 1.17439138796551e-09 | 32.28540632335 | -1.67553888648106 | 1.77260088064751e-07 |
| ENSG00000163701.14 | IL17RE | 163.147533138254 | -1.29226058085582 | 3.46487210197078e-21 | 151.527256171414 | -1.65840645959863 | 5.97392525467161e-28 |
| ENSG00000073282.8 | TP63 | 26.3167615419059 | -4.10930505602427 | 1.6106389630313e-15 | 32.6227982388005 | -1.64577194560328 | 5.75580813499662e-07 |
| ENSG00000168685.10 | IL7R | 11.8864273327478 | -1.89482713828421 | 0.000302300312410601 | 12.3499670120439 | -1.61452887350005 | 0.00259690765754081 |
| ENSG00000170142.7 | UBE2E1 | 978.47987470016 | -1.28694303942448 | 1.27741922837276e-82 | 919.484831104727 | -1.57983084310931 | 2.62188733279338e-127 |
| ENSG00000187672.8 | ERC2 | 12.1108734305113 | -2.09406780085152 | 0.000198240858562194 | 12.9381719912088 | -1.57944024487376 | 0.00426410194067687 |
| ENSG00000079393.16 | DUSP13 | 19.8871316570358 | -2.86563572495297 | 1.62785599335611e-10 | 23.3710975867117 | -1.52380664663749 | 4.38803967326899e-05 |
| ENSG00000183668.13 | PSG9 | 8.45876547291646 | -2.42656902139905 | 0.000369613724353446 | 9.57815659088138 | -1.52354238952538 | 0.0249725520651182 |
| ENSG00000112183.10 | RBM24 | 15.0862862380734 | -1.86584036814742 | 6.94456220706807e-05 | 15.9768187555004 | -1.46989602565481 | 0.00127919217785182 |
| ENSG00000118523.5 | CTGF | 218.642096234912 | -2.51986481465963 | 1.08244987513771e-79 | 251.769611065271 | -1.46157086338312 | 2.19696531995805e-42 |
| ENSG00000203727.3 | SAMD5 | 25.4684687906251 | -1.49593853449456 | 8.96021729215721e-05 | 25.5497261624061 | -1.43054324669756 | 0.000110953563975058 |
| ENSG00000170381.8 | SEMA3E | 67.4074574245334 | -2.17219927141915 | 1.90488114173816e-23 | 75.1089954405767 | -1.42805490961973 | 2.23290579490938e-13 |
| ENSG00000145990.6 | GFOD1 | 40.4657851533104 | -1.12657058468596 | 5.33472402470794e-05 | 38.3809721560164 | -1.3515410217803 | 5.41177306875778e-06 |
| ENSG00000173546.7 | CSPG4 | 139.593793041642 | -1.22020789162077 | 6.1816560990003e-15 | 135.093209804859 | -1.34256073576086 | 7.87981145837664e-16 |
| ENSG00000168427.7 | KLHL30 | 27.9620251763451 | -2.10130426432108 | 1.42118091893686e-09 | 31.4926923480576 | -1.32682285219946 | 0.000102186835826595 |
| ENSG00000006747.10 | SCIN | 37.0824148620567 | -1.88356290360621 | 1.55549644267396e-10 | 40.6232982007547 | -1.31621822749124 | 9.14803699915332e-06 |
| ENSG00000127129.5 | EDN2 | 36.9785906625928 | -3.20862462535094 | 1.15493349838224e-19 | 46.5282504514604 | -1.29290931299074 | 6.10019446279621e-07 |
| ENSG00000158865.8 | SLC5A11 | 35.4295682805915 | -2.67270243537965 | 2.84948443544569e-16 | 42.9512586156668 | -1.26532332279434 | 1.49643901499929e-05 |
| ENSG00000115461.4 | IGFBP5 | 220.190653266747 | -4.63415679654846 | 3.42628502649475e-122 | 297.954714832135 | -1.25788760722151 | 1.11570954404295e-31 |

| | | | | | | | |
|---|---|---|---|---|---|---|---|
| ENSG00000091536.12 | MYO15A | 8.91174555323638 | -1.15772708637244 | 0.0746808592627333 | 8.6783364965619 | -1.25594239925852 | 0.0880436782900824 |
| ENSG00000069974.11 | RAB27A | 129.586698515033 | -1.26743953565931 | 2.43381549234763e-14 | 129.137748067056 | -1.2447299233665 | 1.37477864225004e-15 |
| ENSG00000162614.14 | NEXN | 55.480494780917 | -3.56866180048914 | 2.00021018357783e-31 | 72.9628222041293 | -1.19579933740944 | 1.54089477988093e-09 |
| ENSG00000113578.13 | FGF1 | 33.2885312698034 | -4.73543591774618 | 3.22782407619165e-19 | 45.7509589123122 | -1.19004992881935 | 1.0441382338224e-05 |
| ENSG00000188707.4 | ZBED6CL | 213.742022122602 | -2.26644069256667 | 1.68710916289622e-67 | 253.918945853645 | -1.16730218007651 | 1.52716378819454e-22 |
| ENSG00000159339.9 | PADI4 | 34.4696184342589 | -1.63182961899331 | 2.4305685948944e-08 | 37.3238417139016 | -1.16234908790857 | 0.000113012807957326 |
| ENSG00000154153.9 | FAM134B | 287.117743417104 | -2.42664729622185 | 7.19348564284515e-99 | 348.329279342335 | -1.15448875328847 | 8.20497337173076e-34 |
| ENSG00000135604.9 | STX11 | 15.6396356888271 | -2.15021399971333 | 9.33434618253403e-06 | 18.3745220150114 | -1.15281362983305 | 0.00986318803014571 |
| ENSG00000115602.12 | IL1RL1 | 19.2329129560839 | -1.68315045700181 | 3.52695408998774e-05 | 21.1397571101473 | -1.1343530282653 | 0.00406813307941771 |
| ENSG00000065989.11 | PDE4A | 136.964095225158 | -1.75895064341999 | 4.78930834012995e-25 | 152.918470898052 | -1.12619844778893 | 1.41968359457873e-11 |
| ENSG00000158023.5 | WDR66 | 70.8955372292149 | -3.16756156687285 | 8.70286956254592e-37 | 92.3752407863377 | -1.12438167828664 | 4.58330713941622e-09 |
| ENSG00000163827.8 | LRRC2 | 23.6823694160564 | -2.65772522728161 | 5.36012483410964e-10 | 29.6623110627051 | -1.11829338691906 | 0.00243531685479869 |
| ENSG00000144331.14 | ZNF385B | 19.2799528358408 | -2.81873240436814 | 4.03032885214139e-08 | 24.4484182985863 | -1.11595141489258 | 0.00505336331711121 |
| ENSG00000154040.16 | CABYR | 141.237061152964 | -1.88217585644493 | 1.26433653228221e-38 | 161.403160396541 | -1.1124217308047 | 3.58247610202799e-16 |
| ENSG00000074370.13 | ATP2A3 | 66.1300242653914 | -1.59143571731068 | 2.8386951237908e-14 | 72.0069299133571 | -1.10928100442888 | 4.89830753851725e-08 |
| ENSG00000185885.11 | IFITM1 | 17.6879781946654 | -1.83141722925636 | 0.000277173435765123 | 20.1854564911793 | -1.08116267948154 | 0.0257760007122163 |
| ENSG00000074047.16 | GLI2 | 68.2882799625916 | -1.33692059922397 | 3.16949784122293e-11 | 71.6100671543638 | -1.07471927156171 | 5.14333192765849e-07 |
| ENSG00000116761.7 | CTH | 110.039734569374 | -1.53790881095133 | 6.92445996732504e-19 | 120.01334242416 | -1.07153953165692 | 6.18648582199467e-10 |
| ENSG00000197355.6 | UAP1L1 | 54.6304686119377 | -1.12821693612493 | 1.22204622672123e-06 | 55.1393936357488 | -1.05342211128744 | 2.03373094579423e-05 |
| ENSG00000184226.10 | PCDH9 | 45.2017711123757 | -2.49276646652867 | 2.50499786816676e-17 | 56.4680705801871 | -1.05285478539123 | 2.00696641045476e-05 |
| ENSG00000008311.10 | AASS | 68.5346833632663 | -1.45689995805199 | 5.14108354993405e-11 | 74.1032781978611 | -1.04313866443965 | 6.87568710759677e-07 |
| ENSG00000144642.16 | RBMS3 | 52.179387730477 | -1.78339153162578 | 2.040338148938e-13 | 59.7999570147029 | -1.03617150662384 | 4.9479117864768e-05 |
| ENSG00000118596.7 | SLC16A7 | 45.773328038838 | -1.79878520709534 | 1.10382308091801e-10 | 52.5977395152955 | -1.03270098027523 | 0.00012744170410587 |
| ENSG00000180638.13 | SLC47A2 | 115.122606496401 | -2.12182988712819 | 1.40914043921685e-36 | 138.232622722901 | -1.02975438231701 | 4.68490763946567e-13 |
| ENSG00000142619.4 | PADI3 | 14.0009739703535 | -1.7383232300401 | 0.000792654733448388 | 15.9967178691372 | -1.02788981046274 | 0.0376374423468565 |
| ENSG00000165914.10 | TTC7B | 18.0224598477409 | -1.00215406755637 | 0.0889372826145421 | 17.7638082759308 | -1.02260805792446 | 0.043490699805689 |
| ENSG00000064042.13 | LIMCH1 | 3608.82647169129 | -1.82425781784361 | 1.71049637283761e-221 | 4184.03621695621 | -1.00558764155815 | 1.55900609036844e-96 |
| ENSG00000198121.9 | LPAR1 | 82.7524011909762 | -1.02838805733922 | 8.90544822020881e-08 | 82.5829722766058 | -1.00138130697409 | 2.62593535799126e-07 |
| ENSG00000134198.5 | TSPAN2 | 234.170338148573 | 1.02680286008032 | 6.86355418079589e-20 | 230.125932913839 | 1.00663866957663 | 3.22861590889492e-17 |
| ENSG00000185090.10 | MANEAL | 56.555894306414 | 1.4654038570664 | 5.41097198646775e-11 | 45.73510862497 | 1.04103452485431 | 6.84020720150818e-05 |
| ENSG00000182836.5 | PLCXD3 | 84.0439601683929 | 2.06226836761227 | 7.97679219407158e-27 | 49.3926698961154 | 1.04786392606655 | 7.83355747094016e-05 |
| ENSG00000168010.6 | ATG16L2 | 22.220138925745 | 1.12246414454072 | 0.00249843697190442 | 21.5372568027495 | 1.07042516934289 | 0.0143836536286233 |
| ENSG00000198477.3 | ZNF280B | 16.9224389142644 | 1.09314550394229 | 0.0103927541287795 | 17.1531962596833 | 1.12925660576558 | 0.0167075362862 |
| ENSG00000163207.5 | IVL | 31.1239231016348 | 1.0601992612025 | 0.000498510361975021 | 31.9873949904245 | 1.13944198163592 | 0.000301949383757075 |
| ENSG00000132821.7 | VSTM2L | 18.4300634492903 | 1.97947636447893 | 4.63458476110533e-06 | 11.9438934883874 | 1.16139654182007 | 0.0552345773608458 |
| ENSG00000151233.6 | GXYLT1 | 486.85127226174 | 1.53833589674039 | 1.15858128310552e-47 | 400.977199245766 | 1.16310229439665 | 1.041618398894e-32 |
| ENSG00000226085.2 | UQCRFS1P1 | 28.3014447217204 | 1.76361758423673 | 0.000185830615685178 | 21.3399818122712 | 1.22164943809766 | 0.0495890025097573 |
| ENSG00000159247.8 | TUBBP5 | 11.5526035612443 | 1.46489365336228 | 0.00679424157885586 | 10.1471437717437 | 1.22773592911553 | 0.0437263663187479 |
| ENSG00000123689.5 | G0S2 | 40.4717188514774 | 2.19482080872073 | 1.35962066875273e-13 | 24.0509465211174 | 1.2352872741517 | 0.00130880528126841 |
| ENSG00000084444.9 | KIAA1467 | 167.443692048613 | 1.83286859703944 | 4.73798803814856e-42 | 125.984525285825 | 1.29680624689382 | 3.67215186444255e-17 |
| ENSG00000104951.11 | IL4I1 | 77.3542896841352 | 3.51294599815346 | 1.30999636535008e-36 | 21.9873197384429 | 1.36222857326616 | 0.00108194354568 |

| | | | | | | | |
|---|---|---|---|---|---|---|---|
| ENSG00000157950.10 | SSX2B | 18.0465680485124 | 1.49071453190664 | 0.000251426785025265 | 16.8813599888012 | 1.37487486912069 | 0.0036531483764672 |
| ENSG00000141040.10 | ZNF287 | 13.365934918798 | 1.77136291846969 | 0.000774105303613309 | 10.7936418583187 | 1.38240243357589 | 0.0322798081538113 |
| ENSG00000140323.4 | DISP2 | 34.1415414040237 | 2.21091765926303 | 2.33769569691537e-11 | 23.5647950857034 | 1.53981530015197 | 0.000129545974920389 |
| ENSG00000198039.7 | ZNF273 | 19.7439502245088 | 1.98617661617591 | 3.88535586407635e-05 | 16.0055810548972 | 1.60960284181633 | 0.00308192225242147 |
| ENSG00000100505.9 | TRIM9 | 35.727362108799 | 2.58171688243494 | 1.18896204755912e-14 | 21.3234837958401 | 1.67230286778752 | 3.98743937779829e-05 |
| ENSG00000119547.5 | ONECUT2 | 54.969533887957 | 1.1821216816513 | 2.88029241598519e-07 | 71.6219636350437 | 1.72061424416088 | 2.88890028212325e-14 |
| ENSG00000102098.13 | SCML2 | 9.92866981499034 | 2.45146457275033 | 8.56025008404397e-05 | 7.31399087095578 | 1.92811973618438 | 0.0102006262799044 |
